# Negative regulation of APC/C activation by MAPK-mediated attenuation of Cdc20^Slp1^ under stress

**DOI:** 10.1101/2024.04.02.587770

**Authors:** Li Sun, Xuejin Chen, Chunlin Song, Wenjing Shi, Libo Liu, Shuang Bai, Xi Wang, Jiali Chen, Chengyu Jiang, Shuang-min Wang, Zhou-qing Luo, Ruiwen Wang, Yamei Wang, Quan-wen Jin

## Abstract

Mitotic anaphase onset is a key cellular process tightly regulated by multiple kinases. The involvement of mitogen-activated protein kinases (MAPKs) in this process has been established in *Xenopus* egg extracts. However, the detailed regulatory cascade remains elusive, and also it is unknown whether the MAPKs-dependent mitotic regulation is evolutionarily conserved in the single cell eukaryotic organism such as fission yeast (*Schizosaccharomyces pombe*). Here we show that two MAPKs in *S. pombe* indeed act in concert to restrain anaphase-promoting complex/cyclosome (APC/C) activity upon activation of the spindle assembly checkpoint (SAC). One MAPK, Pmk1, binds and phosphorylates Slp1^Cdc20^, the co-activator of APC/C. Phosphorylation of Slp1^Cdc20^ by Pmk1, but not by Cdk1, promotes its subsequent ubiquitylation and degradation. Intriguingly, Pmk1-mediated phosphorylation event is also required to sustain SAC under environmental stress. Thus, our study establishes a new underlying molecular mechanism of negative regulation of APC/C by MAPK upon stress stimuli, and provides an unappreciated framework for regulation of anaphase entry in eukaryotic cells.

**One-sentence summary:** Inhibitory effect on the activation of anaphase promoting complex/cyclosome (APC/C) by MAPK Pmk1

## INTRODUCTION

The evolutionarily conserved mitogen-activated protein kinase (MAPK) signaling pathways regulate multiple cellular functions in eukaryotic organisms in response to a wide variety of environmental cues (Plotnikov et al., 2011). However, different MAPK pathways have been evolved in an organism to integrate diverse signals and fulfill different regulations on various effectors (Cansado et al., 2021; Ronkina and Gaestel, 2022). This makes specifying the MAPKs and substrates involved in a specific physiological process rather complex and challenging.

The fission yeast *Schizosaccharomyces pombe* has three MAPK-signaling cascades: the pheromone signaling pathway (PSP), the stress-activated pathway (SAP) and the cell integrity pathway (CIP), with Spk1, Sty1 or Pmk1 as MAPK respectively (Figure 1A) (Cansado et al., 2021; Perez and Cansado, 2010). So far, the only known cell-cycle control stages linked to MAPKs in fission yeast are at G_2_/M transition and during cytokinesis (Gomez-Gil et al., 2020; Petersen and Hagan, 2005; Petersen and Nurse, 2007). The p38 MAPK family member Sty1 either arrests or promotes mitotic commitment in unperturbed, stressed or resumed cell cycles after stress depending on the downstream kinases it associates with. Sty1 can phosphorylate kinase Srk1 or Polo kinase Plo1 to negatively or positively regulate Cdc25 phosphatase and thus the onset of mitosis, respectively (Lopez-Aviles et al., 2005; Lopez-Aviles et al., 2008; Petersen and Hagan, 2005; Petersen and Nurse, 2007; Smith et al., 2002). Recent studies have also demonstrated that Sty1 and Pmk1 can pose negative effect on assembly of contractile actomyosin ring (CAR) and cytokinesis in response to actin cytoskeleton damage or cell wall stress (e.g. treatment with blankophor, caspofungin and caffeine) (Edreira et al., 2020; Gomez-Gil et al., 2020), though the detailed mechanisms have not been elucidated.

**Figure 1.**
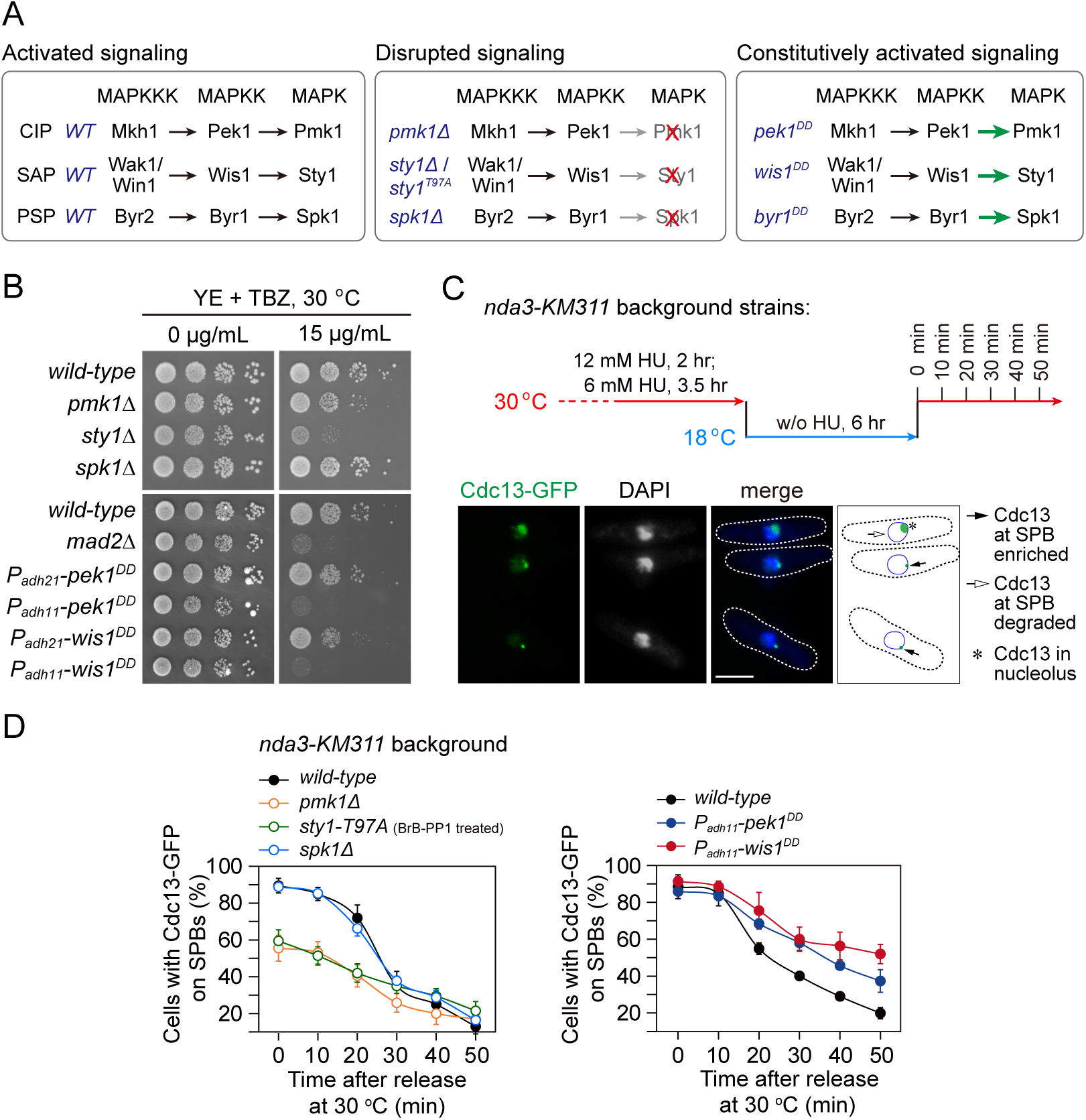
*pmk1*Δ and *sty1-T97A* mutants display spindle checkpoint activation defects, while *pek1^DD^* and *wis1^DD^* mutants are defective in checkpoint silencing. (A) Schematic of core modules of three *Schizosaccharomyces pombe* mitogen-activated protein kinase (MAPK) signaling pathways. Each cascade consists of three core kinases (MAP kinase kinase kinase (MAPKKK), MAP kinase kinase (MAPKK), and MAPK). CIP, the cell integrity pathway; SAP, the stress-activated pathway; PSP, pheromone signaling pathway. These pathways can be disrupted by *pmk1Δ*, *sty1Δ* or ATP analogue-sensitive mutation *sty1-T97A* or *spk1Δ*, and constitutively activated by mutations in MAPKKs Pek1 (*pek1^DD^*, *pek1-S234D;T238D*), Wis1 (*wis1^DD^*, *wis1-S469D;T473D*) or Byr1 (*byr1^DD^*, *byr1-S214D;T218D*), respectively. (B) Serial dilution assay on TBZ sensitivity of all MAPK deletion mutants and *pek1^DD^*- or *wis1^DD^*-overexpressing mutants. *mad2*Δ is a positive control. Note *P_adh11_* is a stronger version of *P_adh21_*promoter. (C) Schematic depiction of the experiment design for time-course analyses on SAC or APC/C activation. *nda3-KM311* cells carrying Cdc13-GFP were grown, synchronized with HU and treated at 18 °C for 6 hours to activate SAC, and finally shifted back to the permissive temperature 30 °C. Samples were collected at 10 min intervals and subjected to microscopy analyses. Example pictures of cells with Cdc13-GFP signals enriched or disappeared at spindle pole bodies (SPBs) are shown. Scale bar, 5 μm. (D) Time-course analyses of SAC activation and inactivation in *nda3-KM311 cdc13-GFP* strains with indicated genotypes. *sty1-T97A* was inactivated by 5μM 3-BrB-PP1. For each time point, ≥300 cells were counted for every sample. The experiment was repeated ≥3 times and error bars indicate mean ± standard deviation.

In late mitosis, the timely polyubiquitylation and subsequent degradation of securin and cyclin B by anaphase-promoting complex/cyclosome (APC/C) play a critical role for anaphase onset and chromosome segregation (Peters, 2006; Sullivan and Morgan, 2007; Yamano, 2019). Given the essential role of APC/C in triggering chromosome segregation, it is not surprising that APC/C is a key molecular target of the spindle assembly checkpoint (SAC), which is an intricate surveillance mechanism that prolongs mitosis until all chromosomes achieve correct bipolar attachments to spindle microtubules (McAinsh and Kops, 2023; Murray, 2011; Musacchio, 2015). Previous studies have revealed that the active MAPK is required for spindle checkpoint activation and APC/C inhibition in *Xenopus* egg extracts or tadpole cells (Chen, 2004; Chung and Chen, 2003; Minshull et al., 1994; Takenaka et al., 1997; Zhao and Chen, 2006). These studies suggested that phosphorylation of several components of SAC, including Cdc20, Mps1 and Bub1, by MAPK serves as regulatory signals to block APC/C activation upon SAC activation (Chen, 2004; Chung and Chen, 2003; Zhao and Chen, 2006). However, all these studies used either the anti-MAPK antibodies or small molecule inhibitor UO126 to inhibit MAPK or MAPK-specific phosphatase MKP-1 to antagonize MAPK activity, these reagents could not distinguish the contributions of different MAPKs with high specificity. Thus, those seemingly defined mechanisms might suffer the ambiguity due to potential specificity issues.

In *S. pombe*, whether MAPKs play key roles in regulation of the APC/C and spindle checkpoint remains largely unexplored. Intriguingly, all genes encoding the major components of three MAPK-signaling cascades in fission yeast are non-essential (Kim et al., 2010), this provides the possibility for investigations on their potential functions in a genetic background with complete deletion of relevant genes. In this work, we set out to directly examine the requirement of the fission yeast MAPKs in mitotic APC/C activation and anaphase entry. Our results uncovered a previously unappreciated mechanism in which Pmk1, the MAPK of CIP, phosphorylates the APC/C activator Slp1 (the Cdc20 homologue in *S. pombe*) upon SAC activation. Phosphorylation of Slp1^Cdc20^ results in its ubiquitylation and degradation and thus subsequently impedes APC/C activation and anaphase entry. This mechanism also operates in response to spindle defects under environmental stress. Therefore, our finding extends cell-cycle control stages linked to MAPKs from previously recognized G_2_/M transition and cytokinesis to anaphase entry and mitotic exit in fission yeast. It will also be interesting to address whether this MAPK-mediated Cdc20 regulation exists as a general mechanism for balancing spindle checkpoint activity in higher eukaryotes.

## RESULTS

### SAP and CIP signaling components are required for spindle checkpoint activity

As the first step to examine the possible requirement of the fission yeast MAPKs in mitotic progression, we tested the sensitivity of deletion mutants of three fission yeast MAPKs to microtubule-destabilizing drug thiabendazole (TBZ), which has been routinely used as an indicator of defective kinetochore or spindle checkpoint (Akera et al., 2015; Saitoh et al., 1997). We noticed that *pmk1*Δ and *sty1*Δ but not *spk1*Δ cells were extremely sensitive to TBZ, and constitutively activated CIP or SAP signaling by overexpressing Pek1^DD^ (Pek1-S234D;T238D) (Sugiura et al., 1999) or Wis1^DD^ (Wis1-S469D;T473D) (Shiozaki et al., 1998) respectively, but not activated PSP signaling by overexpressing Byr1^DD^ (Byr1-S214D;T218D) (Ozoe et al., 2002), also caused yeast cells to be sensitive to TBZ (Figure 1B; Figure 1-figure supplement 1A and Figure 1-figure supplement 2A, B). These observations indicated that Pmk1 and Sty1 pathways might be involved in mitosis-related processes.

To extend our analysis and precisely quantify whether the SAC is properly activated and maintained in MAPK mutants, we adopted one well-established assay using the cold-sensitive β-tubulin mutant *nda3-KM311*, which compromises the kinetochore-spindle microtubule attachment upon cold treatment (Hiraoka et al., 1984), to analyze the ability of cells to arrest in mitosis in the absence of spindle microtubules and enter anaphase when spindle microtubules are reassembled (Bai et al., 2022; May et al., 2017; Vanoosthuyse and Hardwick, 2009). In this assay, the SAC was first robustly activated by the *nda3-KM311* mutant at 18 °C and then inactivated simply by shift mitotically arrested cells back to permissive temperature (30 °C) (Figures 1C). Because Cdc13 (cyclin B in *S. pombe*) localizes to the spindle pole bodies (SPBs) in early mitosis and should be degraded by APC/C to promote metaphase-anaphase transition, accumulation of Cdc13-GFP on SPBs after cold treatment serves as the read-out of the SAC activation, and the disappearance rate of Cdc13-GFP spot under permissive temperature reflects the SAC inactivation efficacy (Figures 1C). It is noteworthy that the ATP analogue-sensitive mutant *sty1-T97A* (i.e. *sty1-as2*) (Zuin et al., 2010), instead of *sty1Δ* deletion mutant, was used in this assay. That was due to the G_2_ delay that has been demonstrated in the *sty1Δ* deletion mutant (Shiozaki and Russell, 1995a), which could interfere mitotic timing for further analysis in this assay. We found that inactivation of CIP or SAP signaling by *pmk1*Δ or *sty1-T97A* compromised full SAC activation, as these two mutants had only less than 60% of cells with Cdc13-GFP on SPBs after 6 hours of cold treatment, whereas the percentage in wild-type and *spk1*Δ cells after the same treatment was roughly 90% (Figure 1D; Figure 1-figure supplement 1C). Consistently, the disappearance of Cdc13-GFP from SPBs in *pek1^DD^*and *wis1^DD^* but not *byr1^DD^* mutant cells released from metaphase-arrest was severely delayed (Figure 1D; Figure 1-figure supplement 1B and Figure 1-figure supplement 2C, D), suggesting delayed SAC inactivation and anaphase onset after release from spindle checkpoint arrest when CIP or SAP signaling is constitutively activated.

To corroborate our above observations of negative effect of CIP or SAP signaling on SAC strength, we took advantage of a different spindle checkpoint activation and metaphase-arresting mechanism involving Mad2 overexpression (May et al., 2017). After *P_nmt1_-mad2^+^* was induced to overexpress for 18 hours, we counted cells with short spindles visualized by GFP-Atb2 as indicative of metaphase-arrest and SAC-activation (Figure 1-figure supplement 3A). We found that the ability of Mad2-overexpressing cells to arrest in mitosis was clearly compromised in *pmk1*Δ and *sty1-T97A* mutants, but not in *spk1*Δ mutant (Figure 1-figure supplement 3B).

All above data suggested that MAPK pathways are evolutionarily conserved in the fission yeast to participate in spindle checkpoint activation and maintenance, of which the CIP and SAP signaling pathways, but not the Spk1 pathway, are actively involved.

### CIP and SAP signaling attenuate protein levels of Slp1^Cdc20^ and facilitate association of MCC with APC/C respectively

The mitotic checkpoint complex (MCC, consisting of Mad3-Mad2-Slp1^Cdc20^ in *S. pombe*) is the most potent inhibitor of the APC/C prior to anaphase (Kapanidou et al., 2017; Primorac and Musacchio, 2013; Sudakin et al., 2001). Previous study has revealed that *Xenopus* MAPK phosphorylates Cdc20 and facilitates its binding by Mad2, BubR1 (Mad3 in *S. pombe*) and Bub3 to form a tight MCC to prevent Cdc20 from activating the APC/C (Chung and Chen, 2003). In fission yeast, the key SAC components Mad2 and Mad3 and one molecule of Slp1^Cdc20^ form MCC which binds to APC/C through another molecule of Slp1^Cdc20^ upon checkpoint arrest, and the recovery from mitotic arrest accompanies the loss of MCC-APC/C binding (May et al., 2017; Sczaniecka et al., 2008; Sewart and Hauf, 2017; Vanoosthuyse and Hardwick, 2009).

Since we found that constitutive activation of CIP or SAP signaling in fission yeast by *pek1^DD^* or *wis1^DD^* mutations caused prolonged SAC activation, we were suspicious that these two MAPK pathways might also execute their functions cooperatively through a similar mechanism to that in *Xenopus*. To test this possibility, we analyzed the MCC occupancy on APC/C by immunoprecipitations of the APC/C subunit Lid1 (Yoon et al., 2002) in *pek1^DD^*and *wis1^DD^* cells arrested by activated checkpoint. Compared to wild-type, more MCC components (Mad2 plus Mad3 and Slp1^Cdc20^) were co-immunoprecipitated in *wis1^DD^* cells, while less of them were co-immunoprecipitated in *pek1^DD^* cells (Figure 2A), suggesting that activation of the SAP but not the CIP pathway enhanced the association of MCC with APC/C.

**Figure 2.**
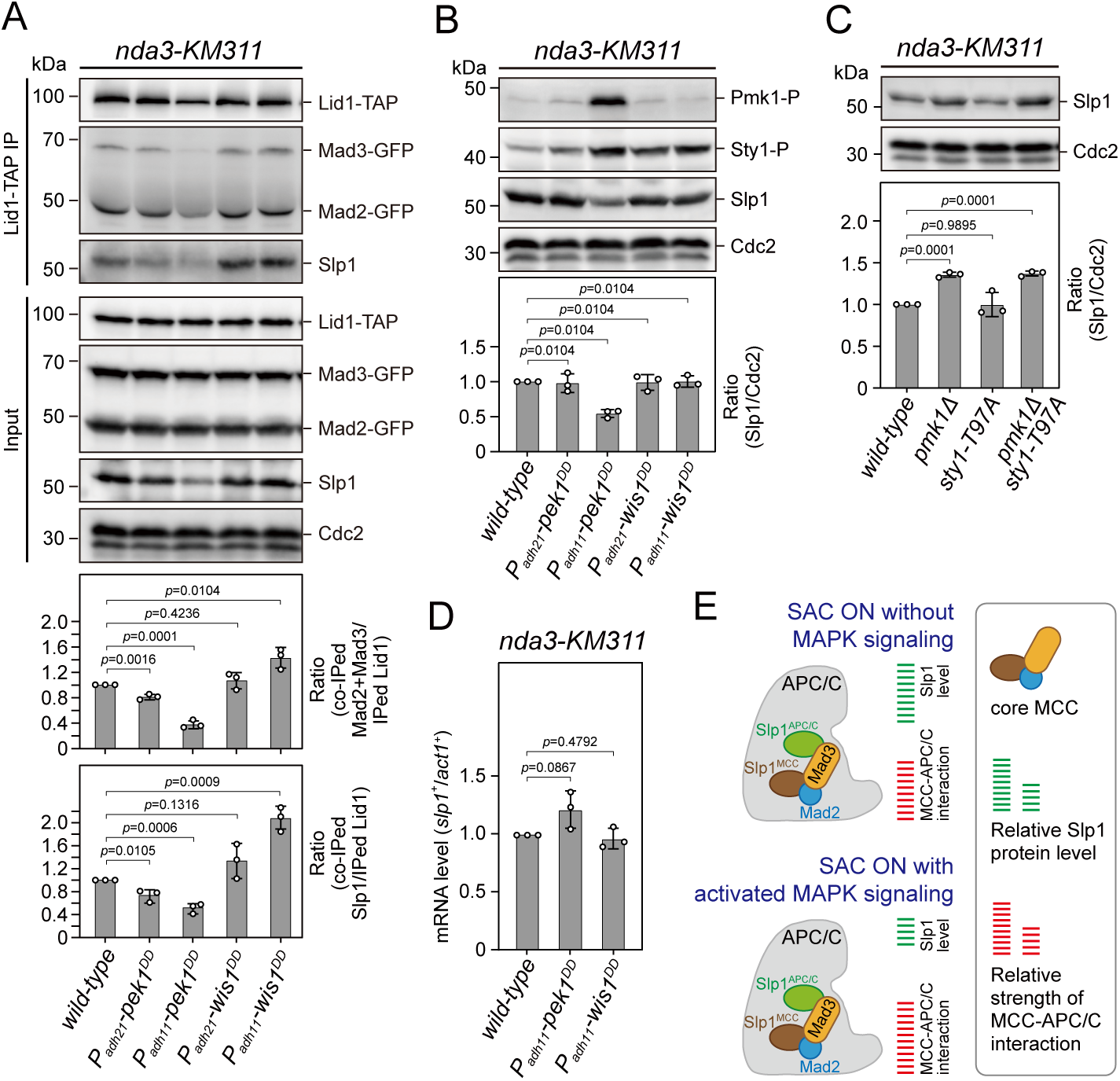
Upon spindle checkpoint activation, Slp1^Cdc20^ levels are reduced in *pek1^DD^* and the MCC-APC/C association is enhanced in *wis1^DD^* cells compared to wild-type cells. (A) Co-immunoprecipitation analysis on MCC-APC/C association. Cells with indicated genotypes were grown at 30 °C to mid-log phase and arrested at 18 °C for 6 hours. Lid1-TAP was immunoprecipitated and associated Mad2, Mad3 and Slp1^Cdc20^ were detected by immunoblotting. The amount of co-immunoprecipitated Mad2, Mad3 and Slp1^Cdc20^ was quantified by being normalized to those of total immunoprecipitated Lid1 in each sample, with the relative ratio between Mad2-GFP plus Mad3-GFP or Slp1^Cdc20^ and Lid1-TAP in wild-type sample set as 1.0. Blots are representative of three independent experiments. *p* values were calculated against wild-type cells. (**B, C**) Immunoblot analysis of Slp1^Cdc20^ abundance in *nda3-KM311* cells treated at 18 °C for 6 hr. Slp1^Cdc20^ levels were quantified with the relative ratio between Slp1^Cdc20^ and Cdc2 in wild-type strain set as 1.0. Phosphorylated Pmk1 (Pmk1-P) or phosphorylated Sty1 (Sty1-P) in (A) were detected using anti-phospho p42/44 and anti-phospho p38 antibodies and represents activated CIP or SAP signaling, respectively. *sty1-T97A* was inactivated by 5μM 3-BrB-PP1. Blots shown are the representative of three independent experiments. *p* values were calculated against wild-type cells. (**D**) RT-qPCR analysis of mRNA levels of *slp1^+^*. Cells with indicated genotypes were grown and treated as in (A-C) before RNA extraction. The relative fold-change (*slp1^+^*/*act1^+^*) in mRNA expression was calculated with that in wild-type cells being normalized to 1.0. Note mRNA level of *slp1^+^* in *pek1^DD^* mutant is not decreased. Error bars indicate mean ± standard deviation of three independent experiments. Two-tailed unpaired *t*-test was used to derive *p* values. (**E**) Schematic summary of the negative effect of activated CIP and SAP signaling on APC/C activation based on primary phenotype characterization of *pmk1Δ*, *sty1-T97A*, *pek1^DD^* and *wis1^DD^* mutants.

In the course of the above immunoprecipitations, we noticed the decrease of co-immunoprecipitated Mad2 and Mad3 in *pek1^DD^* cells, which was largely out of expectation. Further analysis of the inputs of these immunoprecipitations showed that Slp1^Cdc20^ levels were significantly reduced in *pek1^DD^* but not *wis1^DD^* cells (Figure 2A). Our immunoblotting analyses using the total lysate of *pek1^DD^* or *wis1^DD^* cells arrested at metaphase also confirmed these results (Figure 2B). Consistently, the Slp1^Cdc20^ level was significantly increased in *pmk1Δ* and *pmk1Δ sty1-T79A* cells but not in *sty1-T79A* cells (Figure 2C). The elevated Slp1^Cdc20^ level in *pmk1Δ* cells is similar to that we observed in cells harboring two copies of *slp1^+^*, which leads to artificially increased Slp1^Cdc20^ level and also causes a SAC strength defect (Bai et al., 2022). It is noteworthy that attenuated levels of Slp1^Cdc20^ in *pek1^DD^* cells was not due to decreased transcription of *slp1^+^* gene (Figure 2D). These results collectively revealed a negative effect of CIP but not SAP signaling on Slp1^Cdc20^ levels upon SAC activation, which potentially results in the weakened MCC-APC/C interaction detected in CIP mutants (Figure 2A).

Next, we also tested whether activation of the CIP signaling causes downregulation of Slp1^Cdc20^ in normally growing mitotic cells, which were enriched by release of *cdc25-22* background cells from G_2_/M-arrest to 25 °C for 60-90 minutes (Figure 2-figure supplement 1A, B). Intriguingly, we could detect only marginally reduced Slp1^Cdc20^ levels in *pek1^DD^* cells, which was consistent with the slightly retarded anaphase entry in those cells, but we failed to observe any elevated Slp1^Cdc20^ levels in *pmk1Δ* cells (Figure 2-figure supplement 1C). These results suggested that CIP signaling only plays a negligible role in influencing Slp1^Cdc20^ levels when SAC is not activated.

Together, our above data suggested a new mechanism involves division of labor between the two fission yeast MAPK pathways CIP (Pek1-Pmk1) and SAP (Wis1-Sty1) upon SAC activation, which is required in concert to lower Slp1^Cdc20^ levels and enhance MCC affinity for APC/C to strongly inhibit APC/C activity (Figure 2E). In this study, we only focused on characterizing the role of the CIP/Pmk1 in regulating SAC inactivation and anaphase onset hereafter.

### Pmk1 directly binds Slp1^Cdc20^ to attenuate its levels upon SAC activation

To gain further insight into how Slp1^Cdc20^ levels are modulated by Pmk1, we examined the potential interaction of Pmk1 with Slp1^Cdc20^. Through incubating MBP-tagged Slp1^Cdc20^ with metaphase-arrested yeast cell lysates, we found that Pmk1 expressed from yeast cells could be pull-downed by Slp1^Cdc20^ (Figure 3A).

**Figure 3.**
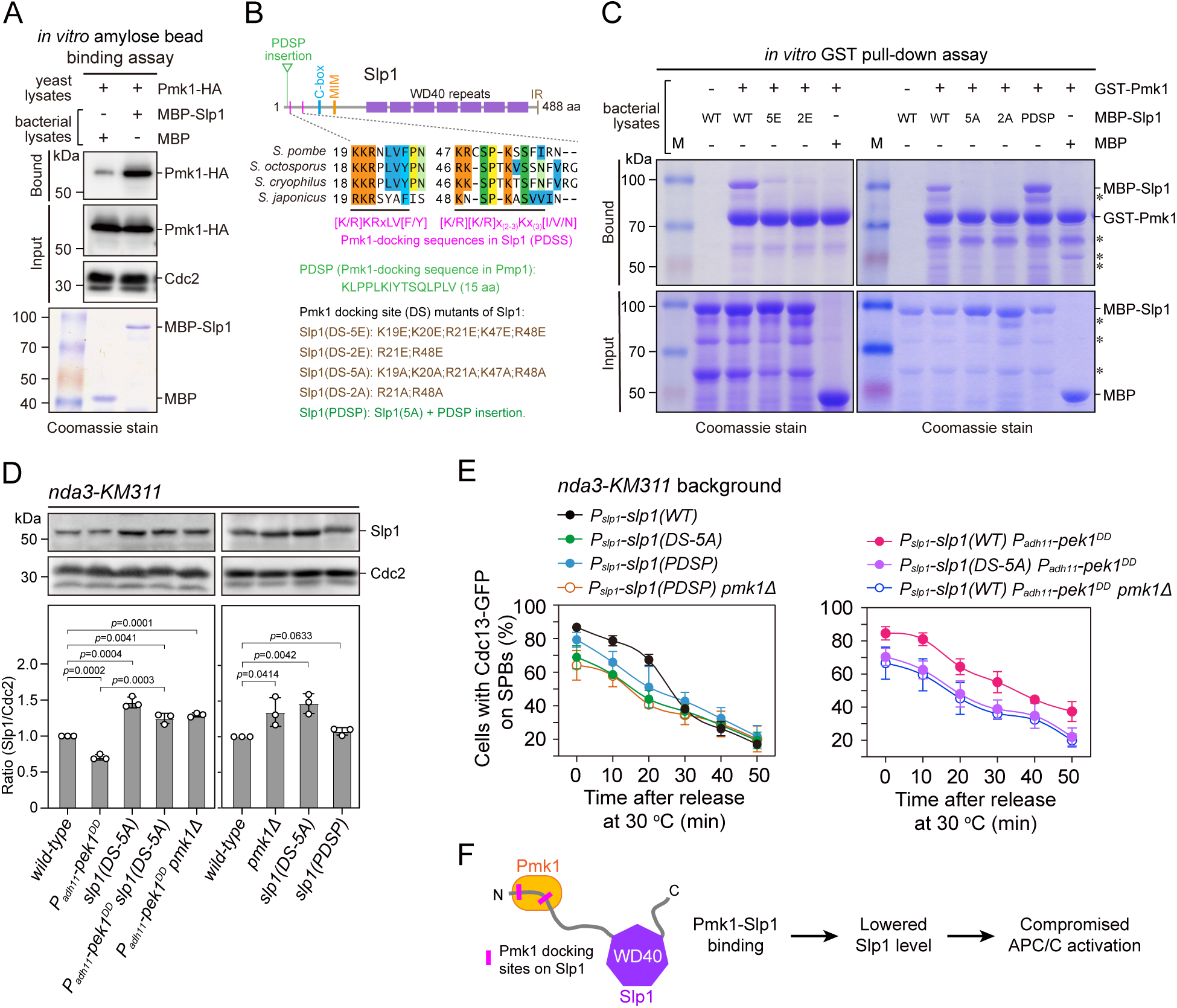
Pmk1 directly binds Slp1^Cdc20^ to attenuate its levels upon spindle checkpoint activation. (**A**) *In vitro* binding assay using bacterially expressed MBP-Slp1^Cdc20^ and yeast lysates prepared from *nda3-KM311 pmk1-HA-6His* cells arrested at 18 °C for 6 hr. Note that weak band detected in MBP sample was due to unspecific background binding to amylose beads. Coomassie blue staining shows inputs for MBP and MBP-Slp1^Cdc20^. (**B**) Schematic depiction of the *S. pombe* Slp1^Cdc20^ protein structure with the positions of two confirmed basic-residue patches mediating Slp1^Cdc20^-Pmk1 association indicated by pink bars. Alignment highlights the conservation of basic-residue patches within 4 *Schizosaccharomyces* species. The deduced Pmk1-docking motifs and different versions of motif mutations are shown. MIM, Mad2-interaction motif; IR, isoleucine-arginine tail. (**C**) *In vitro* GST pull-down assays with bacterially expressed recombinant GST-Pmk1 and MBP-fusions of wild-type Slp1^Cdc20^ or Slp1^Cdc20^ mutants harboring Pmk1-docking motif mutations. An aliquot of the same amount of MBP-Slp1^Cdc20^ as that added in each GST pull-down reactions was immobilized by amylose resin as the input control. Asterisks indicate unspecific or degraded protein bands. (**D**) Immunoblot analysis of Slp1^Cdc20^ abundance in *nda3-KM311* cells with indicated genotypes treated at 18 °C for 6 hr. Slp1^Cdc20^ levels were quantified as in Figure 2B and 2C. The experiment was repeated 3 times. The mean value for each sample was calculated, and *p* values were calculated against wild-type or *pek1^DD^* cells. (**E**) Time-course analyses of SAC activation and inactivation in *nda3-KM311 cdc13-GFP* strains with indicated genotypes. For each time point, ≥300 cells were counted for every sample. The experiment was repeated 3 times and the mean value for each sample was calculated as in Figure 1D. (**F**) Schematic summarizing the negative effect of Pmk1-Slp1^Cdc20^ association on Slp1^Cdc20^ abundance and APC/C activation.

In previous reports, MAPK-docking sites, which are important for interaction with MAPKs, have been identified at the N termini of different MAPKKs from yeast to humans with a notable feature of a cluster of at least two basic residues (mainly Lys/K and Arg/R) separated by a spacer of 2-6 residues from a hydrophobic-X-hydrophobic sequence (Bardwell et al., 2001; Bardwell and Thorner, 1996). Thus, we attempted to identify the potential key residues in Slp1^Cdc20^ that mediate the Pmk1-Slp1^Cdc20^ interaction. By visual scanning of Slp1^Cdc20^ sequence, we noticed five basic-residue patches which are also present in several fungal species and loosely resemble the consensus sequence of the known MAPK-docking sites, and four of them are within N-terminal portion of the protein (Figure 3B and Figure 3-figure supplement 1). However, only mutating either of the two most N-terminal basic-residue patches (19-KKR-21 and 47-KR-48) to glutamates (Glu, E) or alanines (Ala, A) in Slp1^Cdc20^ resulted in decreased amount of Slp1^Cdc20^ pulled down by Pmk1-GST *in vitro* using the bacterially expressed recombinant proteins, and mutating both patches or mutating only two arginine residues (R21 and R48) in those patches was sufficient to completely disrupt their interaction (Figure 3C and Figure 3-figure supplement 2). Importantly, mutations of these Pmk1-docking sequences in Slp1 (PDSS for short) to alanines mimicked *pmk1Δ* mutant in both elevated Slp1^Cdc20^ protein levels (Figure 3D) and defective activation and maintenance of the SAC upon *nda3*-mediated checkpoint arrest (Figure 3E and Figure 3-figure supplement 3). Also, similar to removal of Pek1 downstream MAPK Pmk1, the lowered Slp1^Cdc20^ protein levels and delayed SAC inactivation in *pek1^DD^* mutant could be relieved by *slp1(DS-5A)* mutant, which is deficient in Pmk1-docking (Figure 3D, 3E and Figure 3-figure supplement 3).

Previously, one conserved non-canonical Pmk1-docking sequence which bears positively charged residues and IYT motif has been identified in MAPK phosphatase Pmp1 (Sacristan-Reviriego et al., 2014). Very strikingly, insertion of a 15 amino acid sequence containing the Pmk1-docking sequence in Pmp1 (or PDSP for short) right in front of the first basic-residue patch in Slp1(DS-5A) mutant protein restored its interaction with Pmk1 (Figure 3B and 3C). And this artificially forced Pmk1-Slp1^Cdc20^interaction via PDSP was able to lower Slp1^Cdc20^ protein levels and restore higher percentage of metaphase arrest upon SAC activation in *slp1(DS-5A)* mutant, which is Pmk1-docking-deficient, although the rescuing effect was less efficient than the presence of genuine PDSS (Figure 3D, 3E and Figure 3-figure supplement 3). These data firmly supported the idea that Pmk1 binds Slp1^Cdc20^ to downregulate its levels upon SAC activation.

Collectively, these results demonstrated that Pmk1 directly binds to Slp1^Cdc20^ through two short N-terminal basic patches serving as the major docking sites for Pmk1, and the physical interaction of Pmk1-Slp1^Cdc20^ is involved in attenuation of Slp1^Cdc20^ abundance upon SAC activation (Figure 3F).

### Pmk1 synergizes with Cdk1 to phosphorylate Slp1^Cdc20^

The association of Pmk1 with Slp1^Cdc20^ and the negative effect of constitutively activated CIP signaling on SAC inactivation relies on Pmk1 kinase suggested that Slp1^Cdc20^ may be a direct substrate of Pmk1. To test this possibility, we performed *in vitro* phosphorylation reactions and found that bacterially expressed recombinant GST-Slp1^Cdc20^ could be efficiently phosphorylated in the presence of Pmk1-HA-His purified from yeast cells (Figure 4A).

**Figure 4.**
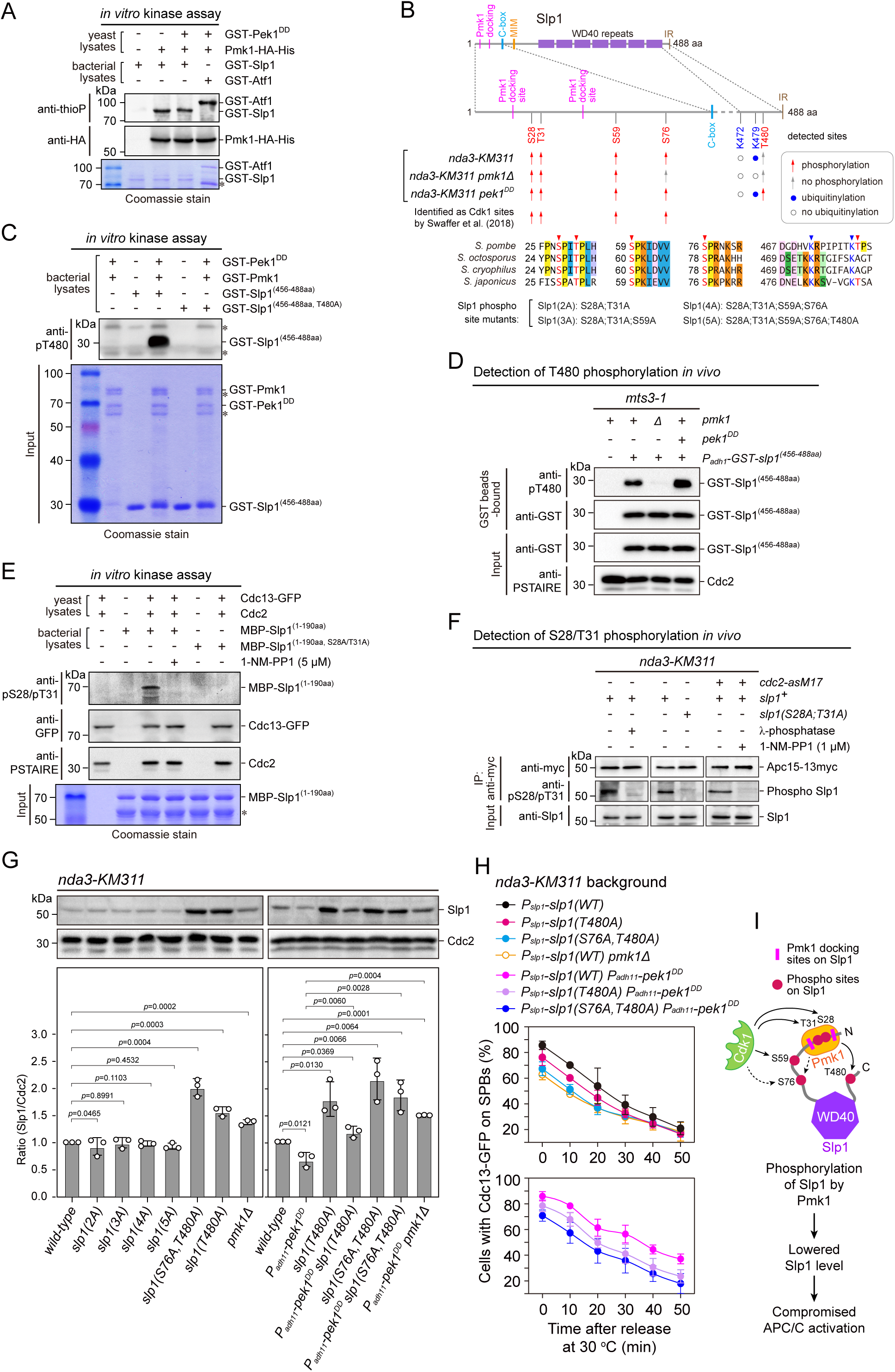
Pmk1 synergizes with Cdk1 to phosphorylate and reduce Slp1^Cdc20^ abundance. (**A**) Non-radioactive *in vitro* phosphorylation assays with bacterially expressed recombinant GST-Slp1^Cdc20^ and Pmk1-HA-His purified from yeast cells. The incorporation of the thiophosphate group was determined using anti-thiophosphate ester antibodies (anti-thioP) as indicative of phosphorylation. Note that the presence or absence of GST-Pek1^DD^ does not affect Slp1^Cdc20^ phosphorylation efficiency. A known Pmk1 substrate Atf1 was used as a positive control (Atf1-P). Asterisks indicate bands corresponding to unspecific or likely degraded proteins. (**B**) Summary of mass spectrometry data on Slp1^Cdc20^ phosphorylation and ubiquitylation *in vivo* in *nda3-KM311*-arrested cells. Red arrows and filled blue circles denote all detected phosphorylated or ubiquitylated sites respectively, while gray arrows and unfilled circles indicate the absence of phosphorylation or ubiquitylation in some of these sites respectively. Alignment highlights the conservation of most of the detected phosphorylation and ubiquitylation sites within 4 *Schizosaccharomyces* species. (**C**) *In vitro* phosphorylation assays with bacterially expressed recombinant GST-fusion of Slp1^Cdc20^ fragment (456-488aa), GST-Pmk1 and GST-Pek1^DD^. The reactions were blotted with pT480 antibodies. Asterisks indicate bands corresponding to unspecific or likely degraded proteins. (**D**) Immunoblot detection of Slp1^Cdc20^ phosphorylation at T480 *in vivo*. GST-slp1(456-488aa) was purified from *mts3-1* cells with indicated genotypes arrested at 36 [for 3.5 hours, and detected with anti-GST and anti-pThr480 antibodies. Note that pThr480 is absent in *pmk1*Δ cells, and enhanced in *pek1^DD^* cells relative to that in wild type cells. (**E**) *In vitro* phosphorylation assays with bacterially expressed recombinant MBP-fusion of Slp1^Cdc20^ fragment (1-190aa) and Cdc13 (cyclin B)-containing Cdk1 complexes purified from metaphase-arrested *nda3-KM311* yeast cells. 1-NM-PP1 was added as inhibitor for analogue-sensitive Cdc2-as. The reactions were blotted with pS28/pT31 antibodies. Asterisks indicate bands corresponding to unspecific or likely degraded proteins. (**F**) Immunoblot detection of Slp1^Cdc20^ phosphorylation at S28/T31 *in vivo*. Apc15-13myc was immunoprecipitated from *nda3-KM311* cells treated at 18 °C for 6 hr and the samples were blotted with pS28/pT31 antibodies. One IP sample from wild type background was treated with λ-phosphatase. For *cdc2-asM17* cells, 1-NM-PP1 was added to inactivate Cdc2 during culturing. (**G**) Immunoblot analysis of Slp1^Cdc20^ abundance in *nda3-KM311* cells treated at 18 °C for 6 hr. Slp1^Cdc20^ levels were quantified as in Figure 2B and 2C. The experiment was repeated 3 times. The mean value for each sample was calculated, and *p* values were calculated against wild-type or *pek1^DD^* cells. (**H**) Time-course analyses of SAC activation and inactivation in *nda3-KM311 cdc13-GFP* strains with indicated genotypes. For each time point, ≥300 cells were counted for every sample. The experiment was repeated 3 times and the mean value and *p* value for each sample were calculated as in Figure 1D. (**I**) Schematic summarizing the negative effect of Slp1^Cdc20^ phosphorylation by Pmk1 on its abundance and APC/C activation.

To identify potential phosphorylation sites *in vivo*, we purified Apc15-GFP from wild type, *pmk1Δ* and *pek1^DD^* cells arrested by activated checkpoint with intention that the Slp1^Cdc20^-containing APC/C complexes could be captured (Figure 4-figure supplement 1A and 1B). Purifications were performed in the presence of phosphatase inhibitors to enrich for phosphopeptides. Mass spectrometry analysis revealed that the phosphorylation of Thr480 residue at the most C-terminus of Slp1^Cdc20^ was dependent on the presence of Pmk1, and this modification was present in *pek1^DD^* but not in wild type cells (Figure 4B, Figure 4-figure supplement 1C-E and Figure 4-figure supplement 2), indicating that phosphorylation of Thr480 is very likely triggered by CIP signaling. We raised phospho-specific antibodies against pT480, it could recognize Slp1^Cdc20^ fragment phosphorylated by Pmk1 *in vitro* and mutation of Thr480 to alanine (T480A) abolished site-specific phosphorylation (Figure 4C). We also confirmed that endogenous Slp1^Cdc20^ was indeed phosphorylated at Thr480 in metaphase-arrested *mts3-1* cells, which was enhanced in *pek1^DD^* mutant and diminished in *pmk1Δ* mutant (Figure 4D). Furthermore, we examined how *in vivo* Slp1^Cdc20^ phosphorylation at Thr480 is affected by forced targeting of Slp1^Cdc20^ to Pmk1. By taking advantage of the high binding affinity of GFP-binding protein (GBP) to GFP and its variants (Chen et al., 2017; Rothbauer et al., 2006), we constructed strains ectopically expressing Slp1^(1-60aa)^-mEGFP-2xNLS-GST-Slp1^(456-488aa)^ and Pmk1-GBP-mCherry, which could bring about artificial tethering of Slp1^Cdc20^ fragments to Pmk1 inside nuclei (Figure 4-figure supplement 3A, B). Immunoblot detection of Slp1^Cdc20^ phosphorylation at T480 *in vivo* demonstrated that indeed the presence of Pmk1-GBP-mCherry could increase pThr480 levels of mEGFP-fusion of Slp1^Cdc20^ fragments, and simultaneously overexpressing Pek1^DD^ from *P_adh11_-6xHA-pek1^DD^* further enhanced the effect (Figure 4-figure supplement 3C). Thus, our results established Thr480 as a major Pmk1 phosphorylation site *in vivo* upon SAC activation.

In addition to pT480, our mass spectrometry analysis also identified Ser28, Thr31 and Ser59 as Pmk1-independent and Ser76 as Pmk1-dependent phosphorylation sites (Figure 4B, Figure 4-figure supplement 1C-E and Figure 4-figure supplement 2). All these four residues are located within the N-terminal unstructured region, fit with the Cdk1 core motif (SP or TP), and more importantly, they are conserved only in four *Schizosaccharomyces* species and several other fungal species but not in higher eukaryotes (Figure 4B, Figure 3-figure supplement 1B and Figure 4-figure supplement 4). Interestingly, they were also proposed as Cdk1 phosphorylation sites based on quantitative phosphoproteomic analysis in a recent study (Swaffer et al., 2018) (Figure 4B). By using the phospho-specific antibodies against both pS28 and pT31, we could detect the phosphorylation of Ser28 and Thr31 *in vitro* by Cdc13 (cyclin B)-containing Cdk1 complexes purified from metaphase-arrested yeast cells, which was diminished when the kinase activity of ATP analogue-sensitive Cdk1 [Cdc2-as1(F84G)] was inhibited by 1-NM-PP1 or when Slp1^(S28A;T31A)^ was added as the substrate in *in vitro* kinase reactions (Figure 4E). Next, we examined whether Ser28 and Thr31 were phosphorylated *in vivo* using immunoblotting with phospho-specific antibodies to directly detect Slp1^Cdc20^, which was co-immunoprecipitated with Apc15 from *nda3*-arrested yeast cells. We observed phosphorylation of Ser28/Thr31 in wild type cells, but it disappeared following phosphatase treatment of wild type samples after immunoprecipitation, 1-NM-PP1 treatment of *cdc2-as1* cells before being collected for immunoprecipitation, or when cells expressing Slp1^(S28A;T31A)^ were used (Figure 4F). Thus, Slp1^Cdc20^ is also phosphorylated by Cdk1 at least at Ser28 and Thr31 in fission yeast mitotic cells.

We wondered whether Slp1^Cdc20^ phosphorylation by Pmk1 affects its stability and APC/C activation. Indeed, phospho-deficient Slp1^T480A^ mutant showed elevated Slp1^Cdc20^ levels in both metaphase-arrested and asynchronously grown cells (Figure 4G and Figure 4-figure supplement 5) and compromised SAC activation compared to wild type cells, likely due to more efficient APC/C activation (Figure 4H and Figure 4-figure supplement 6). Furthermore, Slp1^T480A^ mutant also reversed the inhibitory effect of *pek1^DD^* on Slp1^Cdc20^ stability and APC/C activation after release from SAC-mediated arrest (Figure 4G and 4H).

Ser76 was identified as a Pmk1-dependent phosphorylation residue in our mass spectrometry analysis, but it was considered as a Cdk1 targeting site in an earlier study (Swaffer et al., 2018). It is fairly possible that phosphorylation of Ser76 is promoted by both Cdk1 and Pmk1, which is conceivable as CDKs and MAPKs recognize the same core motif (SP/TP) (Pinna and Ruzzene, 1996), and crosstalk between CDK and MAPK has recently been reported for pheromone signaling in budding yeast (Repetto et al., 2018). We tested how the double phosphorylation sites (S76A;T480A) mutant affects Slp1^Cdc20^ levels or APC/C activity, and observed that the protein levels of Slp1^S76A;T480A^ were further elevated and SAC activation efficiency in this mutant was further lowered compared to Slp1^T480A^ mutant (Figure 4G and 4H). Of note, Slp1^T480A^ and Slp1^S76A;T480A^ mutant proteins became easily detected in interphase cells (Figure 4-figure supplement 5), this is very different from wild-type Slp1 which normally accumulates exclusively in mitosis (Yamada et al., 2000).

We further tested whether replacement of other identified phospho-serine/threonine residues with alanines affects Slp1^Cdc20^ protein levels and APC/C activity. We constructed strains carrying combinations of mutations of S28A, T31A and S59A with S76A, and also with confirmed Pmk1 phosphorylation-deficient T480A, and found that mutations of S28A, T31A and S59A did not affect Slp1^Cdc20^ protein levels, though they could also profoundly compromise SAC activation efficiency similarly to S76A;T480A mutant (Figure 4G, 4H, Figure 4-figure supplement 5, and Figure 4-figure supplement 6). We noticed that the presence of mutations of S28A, T31A and S59A also abolished the elevated Slp1^Cdc20^ protein levels conferred by S76A;T480A mutations (Figure 4G and Figure 4-figure supplement 5). Thus, phosphorylation of S28, T31 and S59 targeted by Cdk1 most likely influences APC/C activation through distinct mechanism from Pmk1-mediated S76/T480 phosphorylation.

Taken together, our data suggested that activated Pmk1 phosphorylates Slp1^Cdc20^ *in vivo*, which leads to lowered Slp1^Cdc20^ levels, delayed SAC inactivation and APC/C activation as we observed in *pek1^DD^*cells. Also, Pmk1 can join Cdk1 to enhance the multi-site phosphorylation of their shared substrate Slp1^Cdc20^ and collaboratively dampen APC/C activity.

### Pmk1- but not Cdk1-mediated Slp1^Cdc20^ phosphorylation promotes its ubiquitylation

Our above data support the notion that the direct binding of Slp1^Cdc20^ by Pmk1 mediates its phosphorylation, which possibly results its subsequent rapid turnover. It has been shown in both human and fission yeast cells that Apc15, one of the conserved components of APC/C, promotes Cdc20/Slp1^Cdc20^ autoubiquitylation and its turnover by APC/C (Mansfeld et al., 2011; May et al., 2017; Sewart and Hauf, 2017; Uzunova et al., 2012). The increased Slp1^Cdc20^ levels in *pmk1Δ* cells (Figure 2C) is reminiscent of the previous observations in *apc15Δ* mutant, which also shows elevated levels of Slp1^Cdc20^ (May et al., 2017; Sewart and Hauf, 2017). We reasoned that the absence of Apc15 may reverse the negative effect of *pek1^DD^* on Slp1^Cdc20^ levels and spindle checkpoint inactivation and APC/C activation. As expected, we indeed observed that deletion of *apc15* recovered Slp1^Cdc20^ levels in *pek1^DD^* cells (Figure 5A), and also lowered the percentage of cells with Cdc13-GFP retained on SPBs in *nda3-KM311 pek1^DD^* cells after cold treatment (Figure 5B and Figure 5-figure supplement 1). These observations are consistent with previous reports showing the relative abundance between checkpoint proteins and the checkpoint target Slp1^Cdc20^ is an important determinant of checkpoint robustness (Heinrich et al., 2013) and *apc15Δ* mutant is spindle checkpoint-defective (May et al., 2017).

**Figure 5.**
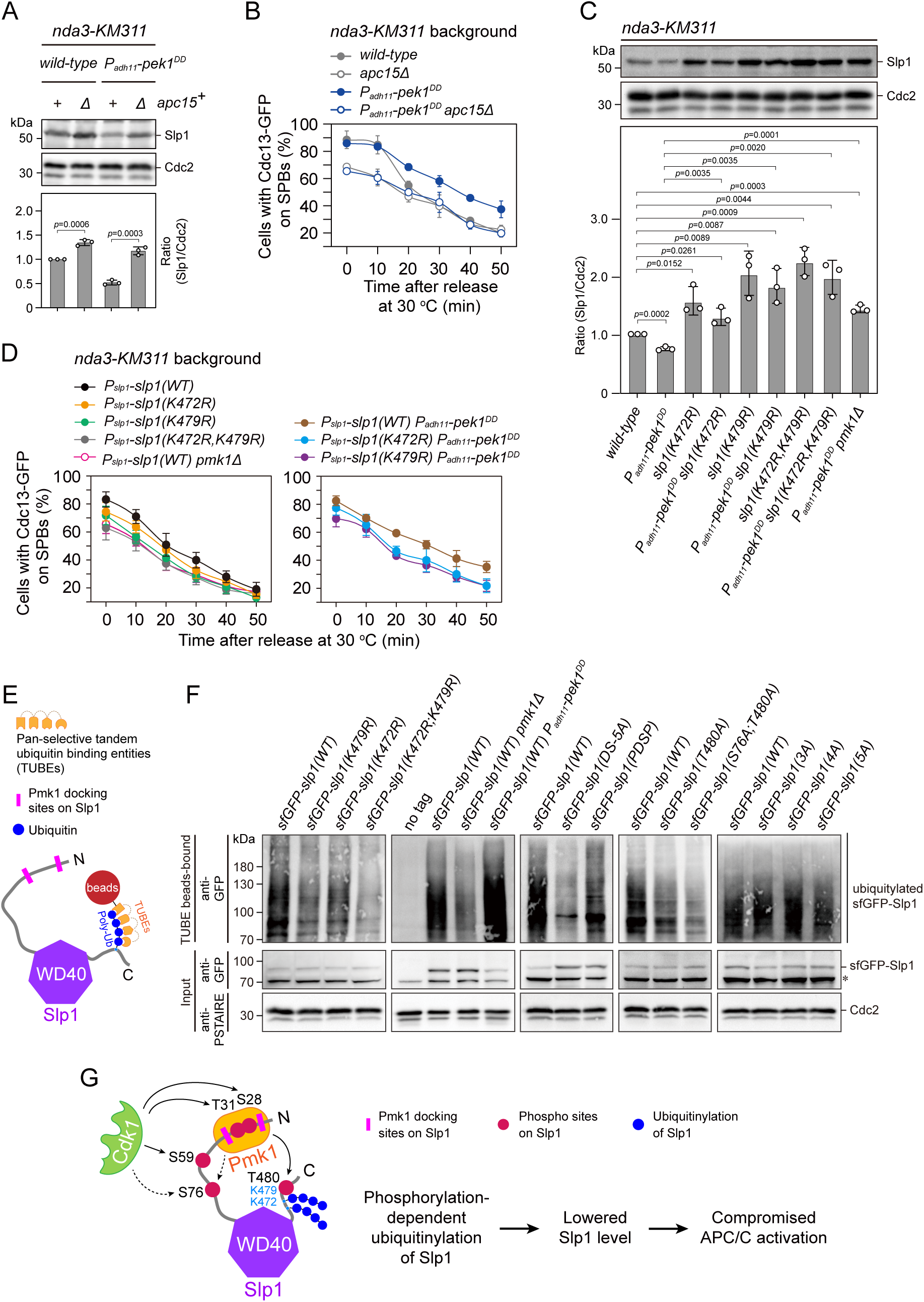
Pmk1- but not Cdk1-mediated Slp1^Cdc20^ phosphorylation promotes its ubiquitylation. (**A**) Immunoblot analysis of Slp1^Cdc20^ abundance in *apc15^+^* or *apc15*Δ background cells with indicated genotypes after being treated at 18 °C for 6 hr. Slp1^Cdc20^ levels were quantified as in Figure 2B. The experiment was repeated 3 times. The mean value for each sample was calculated, and *p* values were calculated against wild-type or *pek1^DD^* cells. (**B**) Time-course analyses of SAC activation and inactivation in *nda3-KM311 cdc13-GFP* strains with indicated genotypes. The experiments were performed and analyzed as in Figure 1D. (**C**) Immunoblot analysis of Slp1^Cdc20^ abundance in K472R, K479R or K472R/K479R mutants. The experiment was repeated 3 times. The mean value for each sample was calculated, and *p* values were calculated against wild-type or *pek1^DD^*cells. (**D**) Time-course analyses of SAC activation and inactivation in *nda3-KM311 cdc13-GFP* strains with K472R, K479R or K472R/K479R mutations. The experiments were performed as in (B). (**E**) Schematic depiction of affinity pull-down assays using TUBE (Tandem Ubiquitin Binding Entity) agarose beads to detect Slp1^Cdc20^ ubiquitylation. (**F**) TUBE pull-down assays in *mts3-1* strains carrying sfGFP-tagged wild type or mutants of Slp1^Cdc20^ or *pmk1Δ* or *pek1^DD^* mutations. The TUBE beads-bound samples were blotted with anti-GFP antibodies. Asterisk indicates unspecific bands recognized by anti-GFP antibodies. (**G**) Schematic summarizing the negative effect of the coupling of Pmk1 phosphorylation and K472/K479-mediated ubiquitylation on Slp1^Cdc20^ abundance and APC/C activation.

During the course of our mass spectrometry analysis on Slp1^Cdc20^ phosphorylation, we also detected Lys 479 (K479) as an ubiquitin-modifying site (Figure 4B, Figure 4-figure supplement 1 and Figure 4-figure supplement 2). We wondered whether K479 is involved in Slp1^Cdc20^ autoubiquitylation in fission yeast. In agreement with the mass spectrometry analysis, mutation of K479 to arginine indeed conferred stability of Slp1^Cdc20^ (Figure 5C), even in asynchronously growing cells (Figure 5-figure supplement 2), which is reminiscent of the stabilized Slp1^Cdc20^ in *apc15Δ* mutant cells at interphase (Sewart and Hauf, 2017). These results suggested that K479 is likely Slp1^Cdc20^ ubiquitylation-relevant. Furthermore, *nda3-KM311 slp1^K479R^* cells also demonstrated lowered metaphase arrest rate after cold treatment (Figure 5D and Figure 5-figure supplement 1), indicating defective SAC activation. Previous studies have identified two neighbouring lysine residues close to the carboxy terminus of human Cdc20, Lys485 and Lys490, as ubiquitylation sites in prometaphase cells, and mutating these two lysines to arginine completely prevented Cdc20 ubiquitylation (Danielsen et al., 2011; Mansfeld et al., 2011). Very interestingly, a second lysine residue, K472, is also present in fission yeast Slp1^Cdc20^ adjacent to K479 (Figure 4B and Figure 5-figure supplement 3). Although K472 was not identified as an ubiquitin-modifying residue in our mass spectrometry analysis, replacement of K472 with arginine also efficiently stabilized Slp1^Cdc20^, similar to K479R mutant (Figure 5C and Figure 5-figure supplement 2). More strikingly, simultaneously mutating both K472 and K479 caused additive effects in elevating Slp1^Cdc20^ levels and lowering SAC activation rate in wild-type or *pek1^DD^* strain background (Figure 5C, 5D, Figure 5-figure supplement 1 and Figure 5-figure supplement 2), suggesting that likely both these two lysine residues at the C-terminal tail of Slp1^Cdc20^ contribute to its turnover via ubiquitylation, which is very similar to the mechanism in human Cdc20. Consistently, using TUBEs (tandem ubiquitin-binding entities) (Hjerpe et al., 2009) to enrich for ubiquitinated proteins in metaphase-arrested *mts3-1* cells expressing sfGFP-Slp1, we were able to demonstrate that mutating both K472 and K479 largely removed higher molecular weight bands detected by immunoblotting, which were corresponding to ubiquitylated species of Slp1^Cdc20^ (Figure 5E, F). We also noticed that Slp1^K472R; K479R^ rendered more severely compromised viability than either single mutants (Figure 5-figure supplement 4), indicating blocking Slp1^Cdc20^ ubiquitylation could bring about detrimental consequence to the cells.

Our observation that Slp1^Cdc20^ levels were lowered in *pek1^DD^*mutant (Figure 2B) suggested that attenuation of Slp1^Cdc20^ levels upon constitutive activation of CIP signaling is very likely through facilitating its autoubiquitylation. To test this possibility, we examined and compared the ubiquitylation levels of sfGFP-Slp1 in wild type, *pmk1*Δ and *pek1^DD^* cells by affinity pull-down assays using TUBEs (Figure 5E). Indeed, we observed a much weaker or stronger discrete laddering pattern of sfGFP-Slp1 in *pmk1*Δ and *pek1^DD^* cells respectively than that of wild type cells, indicating abated or enhanced ubiquitylation (Figure 5F). Furthermore, mutations of T480A, S76A/T480A and Pmk1-docking site, but not S28A/T31A- or S28A/T31A/S59A-containing mutants, also reduced polyubiquitylation of Slp1^Cdc20^ in both *pek1^+^* cells (Figure 5F) and *pek1^DD^*-overexpressing (i.e. *P_adh11_*-*pek1^DD^*) cells (Figure 5-figure supplement 5). All these results confirmed that Pmk1- but not Cdk1-mediated Slp1^Cdc20^ phosphorylation promotes its ubiquitylation.

Together, these findings lent further support to the idea that the CIP signaling pathway restrain APC/C activity through a cascade mechanism, in which Pmk1 directly binds and phosphorylates Slp1^Cdc20^ to facilitate its autoubiquitylation, and eventually lowers its levels (Figure 5G).

### Osmotic stress and cell wall damage trigger rapid Slp1^Cdc20^ downregulation and mitotic exit delay

It has been well established that two MAPKs Sty1 and Pmk1 can be activated by environmental perturbations including osmotic stress, cell wall damage, thermal stress, oxidative stress, and glucose limitation, among others (Cansado et al., 2021; Madrid et al., 2006; Shiozaki and Russell, 1995b). We wondered whether the phosphorylation of Slp1^Cdc20^ by Pmk1 could tune the strength of SAC activation and APC/C activity in response to extracellular stress. To investigate this possibility, we performed arrest-and-release experiments with *nda3-KM311* mutants and added 0.6 M KCl or 2 µg/mL of caspofungin 1 hr before release from metaphase arrest to elicit strong osmotic saline stress (KCl) or cell wall damage stress (caspofungin) response (Madrid et al., 2006; Madrid et al., 2016), respectively (Figure 6A). As previously reported, treatment with KCl or caspofungin resulted in full activation of both Pmk1 and Sty1 (for KCl treatment) or only Pmk1 (for caspofungin treatment) as determined using anti-phospho p42/44 or anti-phospho p38 antibodies, which serve as indicative of respective Pmk1 and Sty1 activation (Figure 6B). Intriguingly, treatment with KCl or caspofungin also correlated with a marked and rapid decrease in Slp1^Cdc20^ levels (Figure 6B). It is of note that SAC activation *per se* did not trigger activation of CIP pathway, as we did not observe any elevated Pmk1 phosphorylation in metaphase-arrested cells either by *nda3*-*KM311* mutation (see “control” samples in Figure 6B) or by Mad2-overexpression (Figure 6-figure supplement 1).

**Figure 6.**
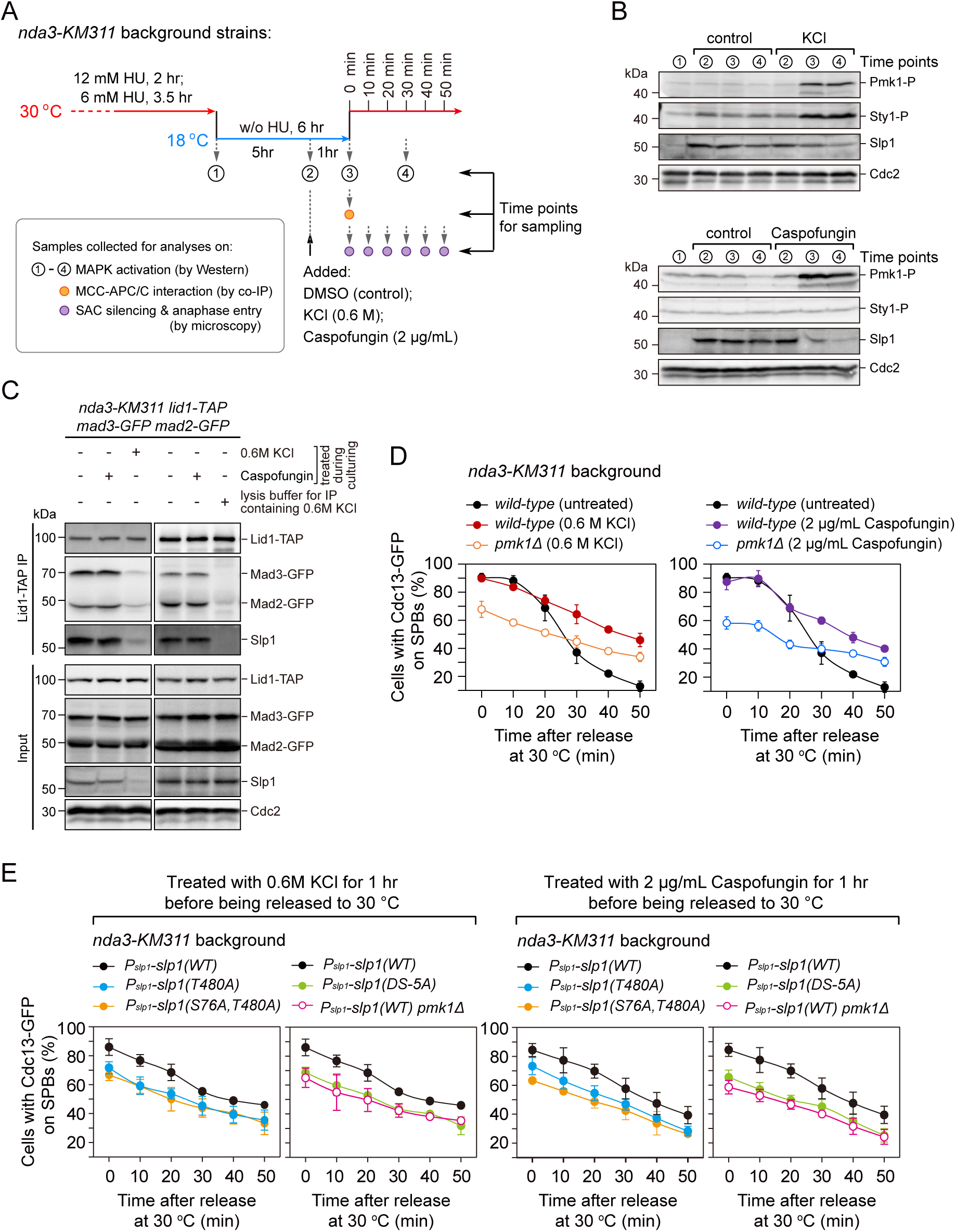
Osmotic stress and cell wall damage trigger rapid Pmk1 phosphorylation-dependent Slp1^Cdc20^ downregulation and mitotic exit delay. (A) Schematic depiction of the experimental design for treatment with KCl or caspofungin during *nda3*-mediated spindle checkpoint activation to activate MAPKs. Samples were collected at indicated time points for subsequent analyses including immunoblotting, co-IP and time-course analysis on SAC or APC/C activation. (**B**) Immunoblot analysis of activation of MAPKs and Slp1^Cdc20^ protein levels. Samples with or without indicated treatments were blotted with anti-phospho p42/44 and anti-phospho p38 antibodies as indicative of phosphorylated Pmk1 (Pmk1-P) or phosphorylated Sty1 (Sty1-P), respectively. Slp1^Cdc20^ levels were detected with anti-Slp1 antibodies and anti-Cdc2 was used as loading control. (**C**) Co-immunoprecipitation analysis of APC/C-MCC association upon environmental stress. Lid1-TAP was immunoprecipitated from *nda3-KM311*-arrested cells and associated Mad2-GFP, Mad3-GFP and Slp1^Cdc20^ were detected as in Figure 2A. Note that APC/C-MCC association was disrupted when 0.6 M KCl was present during cell culturing or during immunoprecipitation procedures. (**D**) Time-course analyses of SAC activation and inactivation in *nda3-KM311 cdc13-GFP* strains with indicated genotypes after arrest at 18 °C and KCl or caspofungin treatments. The experiment was repeated 3 times and the mean value and *p* value for each sample were calculated as in Figure 1D. (b) Time-course analyses of SAC activation and inactivation efficiency in Pmk1-docking- and phosphorylation-deficient *slp1* mutants under environmental stresses elicited by 0.6 M KCl or 2 μg/mL caspofungin.

Next, we investigated whether environmental perturbations affect APC/C-MCC interaction. Surprisingly, we did not see any alteration of APC/C-MCC interaction accompanying Pmk1 activation when cells were treated by caspofungin (Figure 6C), which was different from what we observed in *P_adh11_-pek1^DD^* cells (Figure 2A). Also, we failed to detect enhanced APC/C-MCC association like we observed in *P_adh11_-wis1^DD^* cells when cells were treated by KCl, where Sty1 was activated (Figure 6C). The latter observation was most likely due to the disruptive effect of high concentration of KCl on protein-protein interactions, because we observed similar effect when we immunoprecipitated APC/C in the presence of 0.6 M KCl in the lysis buffer (Figure 6C, right panel). With this data, we could not exclude the possibility that activated Sty1 transiently enhanced APC/C-MCC association in yeast cells, which was soon counteracted by the disruptive effect of saline. Nevertheless, the high SAC activation efficiency in mitotis-arrested cells exposed to osmotic stress (KCl) or cell wall damage stress (caspofungin) was largely lowered by *pmk1* deletion (Figure 6D and Figure 6-figure supplement 2), indicating that at least activated Pmk1 exerts regulation of the spindle checkpoint and APC/C activation under environmental stimuli. Further, Pmk1-docking site mutant *slp1(DS-5A*) and Pmk1 phosphorylation-deficient mutants *slp1^T480A^* and *slp1^S76A;T480A^* could partially suppress the sustained SAC activation responding to osmotic and cell wall damage stresses (Figure 6E and Figure 6-figure supplement 2). And accordingly, protein levels of Pmk1 phosphorylation-deficient mutants Slp1^T480A^ and Slp1^S76A;T480A^ under above stresses were restored almost to the wild-type Slp1 levels in untreated metaphase-arrested cells (Figure 6-figure supplement 3). We also noticed that protein levels of ubiquitylation-deficient mutants, including Slp1^K472R^, Slp1^K479R^ and Slp1^K472R;K479R^, were more efficiently maintained than Slp1^T480A^ and therefore more resistant to osmotic and cell wall damage stresses (Figure 6-figure supplement 3).

### *pmk1***Δ** cells are defective in faithful chromosome segregation upon environmental stress

Our above data demonstrated that, upon SAC activation, Pmk1 is involved in controlling the stability of Slp1^Cdc20^ and thus the activation of APC/C and anaphase entry under environmental stresses. We wondered whether the absence of *pmk1^+^* would affect fidelity of chromosome segregation, particularly when cells are exposed to environmental stimuli. To answer this question, we treated G_2_-arrested and -released *cdc25-22* cells with 0.6 M KCl or 2 μg/mL caspofungin to activate MAPKs (Figure 7A), and then examined possible chromosome segregation defects in *pmk1*Δ cells (Figure 7B). We found that stressed *pmk1*Δ cells displayed greatly increased frequency of lagging chromosomes and chromosome mis-segregation at mitotic anaphase compared to similarly treated wild-type cells and also untreated *pmk1*Δ cells (Figure 7C). This data further validated the importance of Pmk1 activation under adverse stress in delaying anaphase onset and maintaining genome integrity.

**Figure 7.**
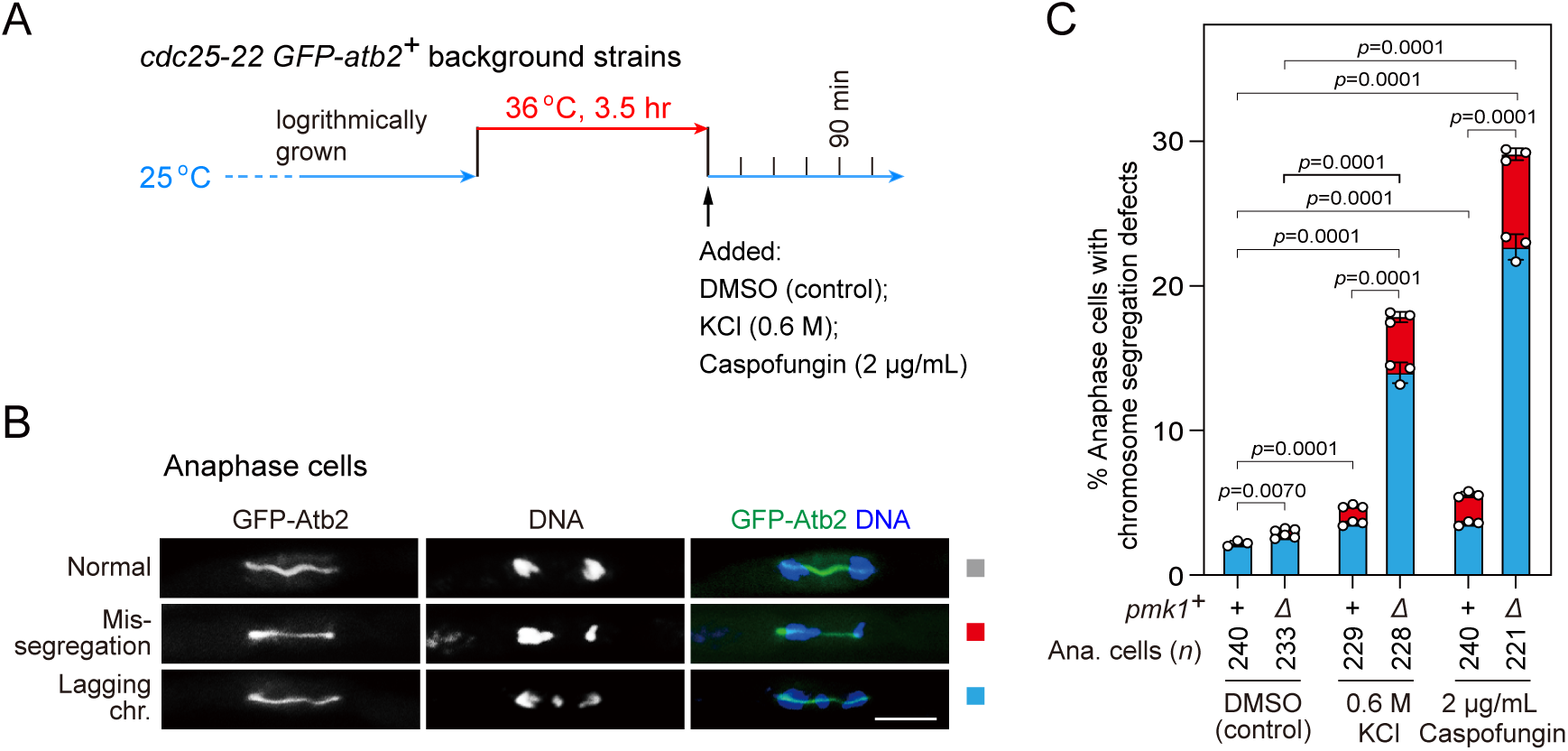
*pmk1*Δ cells are defective in faithful chromosome segregation upon environmental stress. **(A)** Schematic depiction of the experiment design for arrest-and-release of *cdc25-22 GFP-atb2*^+^ strains. Environmental stresses were imposed by 0.6 M KCl or 2 μg/mL caspofungin. Samples were taken at 90 min after being shifted back from 36 °C to 25 °C to enrich anaphase cells **(B and C)** Analyses of chromosome segregation in anaphase cells treated without or with KCl or caspofungin and after being fixed and stained with DAPI. Example pictures of anaphase cells with long spindles judged by GFP-Atb2 signals are shown (B). Anaphase cells with two categories of chromosome segregation defects (i.e. unequal chromosome segregation (mis-segregation) and lagging chromosomes) were quantified (C). >200 cells were counted for every sample. The mean values for each category were calculated, error bars indicate mean ± standard deviation of three independent experiments. *p* values were calculated with pooled data of two categories for each sample. *n*, numbers of anaphase cells analyzed. Scale bar, 5 μm.

## DISCUSSION

### MAPKs are involved in regulation of mitotic anaphase onset in fission yeast

APC/C-mediated proteolysis of cyclin B and securin is key for anaphase entry by inactivating Cdk1 and permitting chromosome segregation, respectively. In response to unattached or tensionless kinetochores, the SAC induces MCC generation. As a critical APC/C inhibitor, MCC fulfills its function through the physical binding of BubR1/Mad3 and Mad2 to two molecules of Cdc20/Slp1^Cdc20^ (*i.e.* Cdc20^MCC^ and Cdc20^APC/C^), and this mechanism has been confirmed at least in both humans and fission yeast (Alfieri et al., 2016; Chao et al., 2012; Izawa and Pines, 2015; Sewart and Hauf, 2017; Yamaguchi et al., 2016). Thus, Cdc20 is unique amongst the MCC proteins in that it is simultaneously a coactivator and inhibitor of the APC/C.

In the current study, we have uncovered a dual mechanism, in which two fission yeast MAPKs Pmk1 and Sty1 restrain APC/C activity through distinct manners, either by phosphorylating Slp1^Cdc20^ to lower its levels or by phosphorylating unknown substrate(s) to promote MCC-APC/C association and thus SAC signaling strength, respectively (Figure 8). Therefore, our study has established another critical APC/C-inhibitory mechanism in the spindle checkpoint, though it will be important for future studies to identify the phosphorylation target(s) posed by SAP signaling pathway. We should mention that most of our experimental setups in this study were based on cold-sensitive *nda3-KM311* mutant, which causes loss of both kinetochore-microtubule attachments and intra-kinetochore tension when being grown at 18 °C. However, we were unable to distinguish whether MAPK involvement in SAC dynamics is relevant to one perturbation or another or both. Nevertheless, our current study together with earlier report (Edreira et al., 2020) have implicated fission yeast Pmk1 in both the mitotic and cytokinesis surveillance mechanisms in response to unfavorable environmental stimuli.

**Figure 8.**
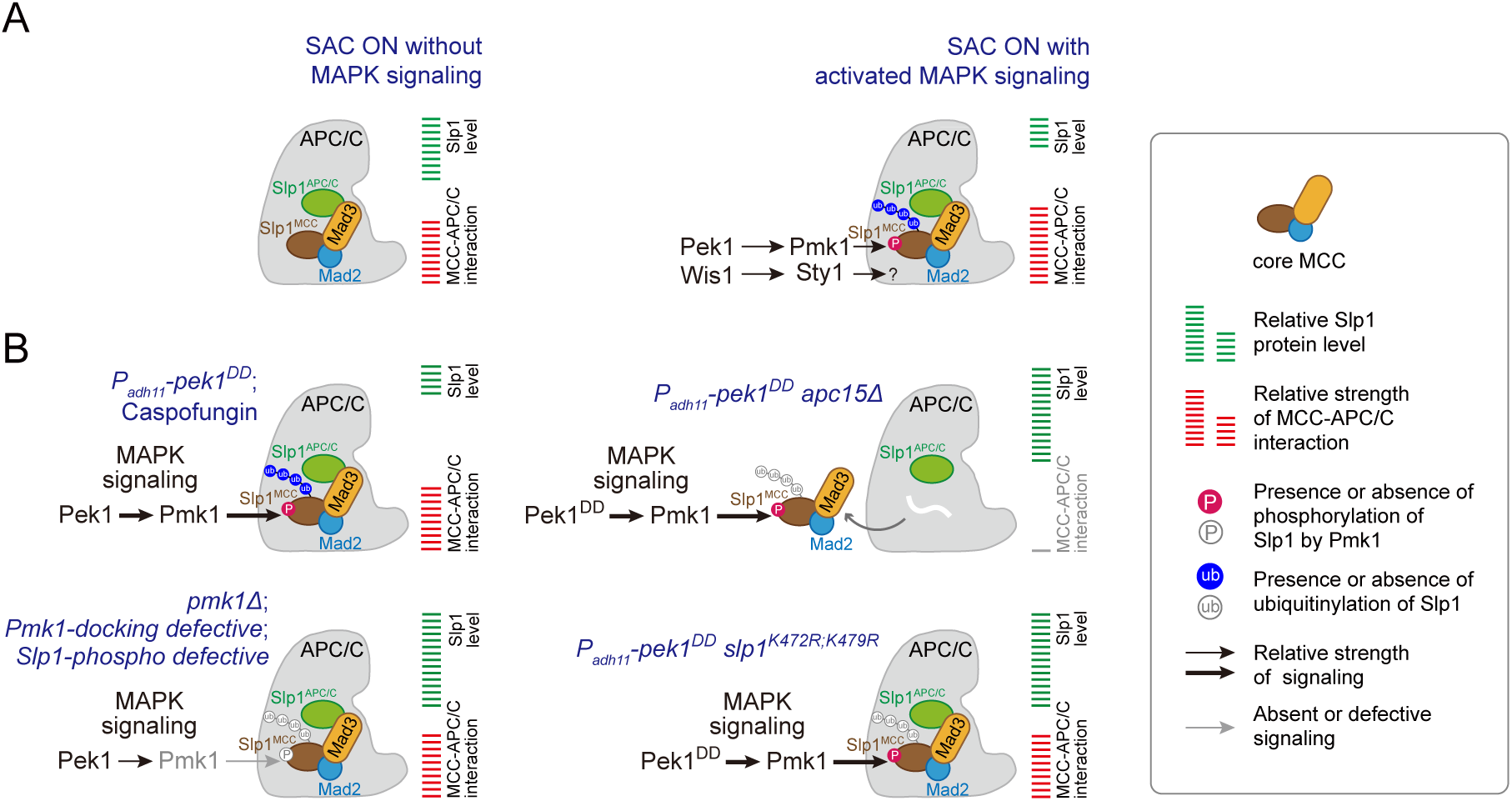
Summary of the mechanisms that how MAPKs negatively regulate APC/C activity in fission yeast. (**A**) Schematic depiction of a possible dual mechanism that how activated MAPK signaling pathways are involved in delaying APC/C activation and SAC inactivation. Upon SAC and MAPK signaling activation, division of labor between the CIP and the SAP pathways enables the phosphorylation of Slp1^Cdc20^ and unidentified substrate(s) by Pmk1 and Sty1 respectively, which leads to lowered Slp1^Cdc20^ level and enhanced MCC affinity for APC/C. (**B**) Summary of the alteration of Slp1^Cdc20^ protein levels and MCC-APC/C association strength under various conditions examined in this study. White wavy line indicates the absence of Apc15.

The regulation of mitotic entry by MAPK in fission yeast is triggered by many environmental insults, including high osmolarity, oxidative stress, heat shock, centrifugation, nutrient starvation and rapamycin treatment (Hartmuth and Petersen, 2009; Petersen and Hagan, 2005; Petersen and Nurse, 2007; Shiozaki and Russell, 1995a; Shiozaki et al., 1998). We found that treatment of fission yeast cells with KCl or caspofungin, which provokes a strong osmotic stress or cell wall damage response and quick Pmk1 activation, correlated with an attenuated Slp1^Cdc20^ protein levels and delayed APC/C activation. Thus, our finding implicates MAPK Pmk1 as an important regulator for anaphase entry in response to adverse extracellular conditions. Intriguingly, one previous study showed that treatment of human cells with the carcinogen cadmium activates MAPK p38 (fission yeast Sty1 homologue) and triggers accelerated ubiquitination and proteolysis of Cdc20, which is essential for prometaphase arrest under SAC (Yen and Yang, 2010). Therefore, although different MAPKs might be employed to fulfil the task in yeast and higher eukaryotes, it is still possible that phosphorylation and regulation of Cdc20 levels by MAPKs is an evolutionarily conserved mechanism to adjust mitotic exit to extracellular or environmental stimuli.

### Negative phospho-regulation of Cdc20 by MAPK as a spindle assembly checkpoint-dependent APC/C-inhibitory mechanism

APC/C activity can be regulated at multiple levels, Cdk1-dependent APC/C phosphorylation is one of the major prerequisites for APC/C activation and anaphase onset, in which Apc3 and Apc1 are key subunits for positive phospho-regulation (Fujimitsu et al., 2016; Kraft et al., 2003; Qiao et al., 2016; Steen et al., 2008; Zhang et al., 2016). However, it has been shown that Cdk1-dependent phosphorylation of APC/C co-activator Cdc20 lessens its interaction with APC/C, thus serves as a negative phospho-regulation (Labit et al., 2012). In addition to its phosphorylation by Cdk1, phosphorylation of Cdc20 by multiple other kinases, such as Bub1, PKA, MAPK and Plk1, has also been demonstrated or suggested in *Xenopus*, humans or budding yeast to be inhibitory to APC/C activation and improper anaphase onset upon SAC activation ((Jia et al., 2016) and reviewed in (Yu, 2007)). Therefore, Cdc20 generally serves as an integrator of multiple intracellular signaling cascades that regulate progression through mitosis.

Although multiple kinases have been proposed to be directly involved in phosphorylation of Cdc20 proteins (Yu, 2007), precisely identifying the *in vivo* phosphorylation sites in Cdc20 has turned out to be challenging, largely due to its protein scarcity and instability. Based on known Cdk1 core consensus (SP/TP), 9 residues in human Cdc20 were originally suggested as putative Cdk1 sites, but among them only Ser41 has been proven to be phosphorylated *in vivo* by mass spectrometry analysis (Kramer et al., 2000; Yudkovsky et al., 2000). However, subsequent mass spectrometry analyses identified Ser41 as one of the Bub1 phosphorylation sites both *in vivo* and *in vitro* and another putative Cdk1 site Thr106 as one of the Plk1 sites (Jia et al., 2016; Tang et al., 2004). It is very likely that there are not as many genuine Cdk1 sites in Cdc20 as originally assumed. One supporting evidence is that mutation of only one (Thr32) of the three putative Cdk sites to alanine in N terminus of *C. elegans* CDC-20 is sufficient to phenocopy 3A mutant for accelerated APC/C activation (Kim et al., 2017). Interestingly, Thr32 of *C. elegans* CDC-20 is conserved as Thr 59 and Thr68 in human and *Xenopus* Cdc20, respectively (see Figure 4-figure supplement 4). Based on two-dimensional tryptic phosphopeptide analysis, Thr68 in *Xenopus* Cdc20 has also been regarded as a MAPK target site (Chung and Chen, 2003). These studies in diverse organisms raised an issue that whether multiple kinases share the same phosphorylation sites in Cdc20? It would not be so surprising if it turns out to be the case eventually, at least for Cdk1 and MAPKs, because CDKs and MAPKs recognize the same core motif (SP/TP) (Pinna and Ruzzene, 1996).

By purifying Apc15-aasociated fission yeast Slp1^Cdc20^ proteins from checkpoint-arrested *pek1^DD^* and *pmk1*Δ cells, we were able to identify Thr480 and Ser76 as *in vivo* Pmk1-relevant phosphorylation sites and three Pmk1-independent phosphorylation sites (S28, T31, S59) based on mass spectrometry analyses. All our identified sites except Thr480 are located in the N terminal disordered portion of Slp1^Cdc20^ sequence, this is similar to the phosphorylation residues identified by biochemical and functional analyses in other organisms, in which the major phosphorylation sites all fall within the N-terminus of Cdc20 homologues. We further provided evidence that S28/T31 and T480 are phosphorylated by Cdk1 and Pmk1, respectively, both *in vitro* and *in vivo*. We noticed that there is a tendency for reported phosphorylation residues to be concentrated in the N terminus of Cdc20 proteins (Jia et al., 2016; Tang et al., 2004; Yu, 2007). However, due to the lack of clear sequence homology in their N-terminal and C-terminal tails between Slp1^Cdc20^ and its Cdc20 homologues in higher eukaryotes (see Figure 4-figure supplement 4), it is impossible to certainly assign any identified sites from different organisms as homologous sites. But it is possible that multiple kinases modify shared sites in Cdc20, this collaborative effect may be especially helpful under adverse environmental stress. Nevertheless, all reported Cdc20 phosphorylation events constitute an APC/C-inhibitory mechanism, no matter what kinases are responsible for the modifications (Chung and Chen, 2003; D’Angiolella et al., 2003; Jia et al., 2016; Kim et al., 2017; Kramer et al., 2000; Labit et al., 2012; Tang et al., 2004; Yu, 2007; Yudkovsky et al., 2000).

### Coupling of phosphorylation and autoubiquitylation of Slp1^Cdc20^

The current model of inhibitory signaling from Cdc20 posits that its phosphorylation either promotes the formation of MCC or inhibits APC/C^Cdc20^ catalytically, both are required for proper spindle checkpoint signaling (Jia et al., 2016; Labit et al., 2012; Yamano, 2019). Interestingly, our current study suggests Slp1^Cdc20^ phosphorylation by fission yeast MAPK Pmk1 may represent another critical APC/C-inhibitory mechanism in the spindle checkpoint through reducing the protein levels of Slp1^Cdc20^, which is achieved via autoubiquitylation-mediated degradation (Figure 8). This mechanism is distinct from that reported in *Xenopus* egg extracts and tadpole cells, in which phosphorylation of Cdc20 by MAPK is instead responsible for facilitated Cdc20 binding by MCC to prevent it from activating the APC/C (Chung and Chen, 2003). Actually, previous studies in fission and budding yeast have underlined the importance of accurate relative abundance between checkpoint proteins and Cdc20/Slp1^Cdc20^, which sets an important determinant of checkpoint robustness (Heinrich et al., 2013; Pan and Chen, 2004). A benefit of allowing the association of MCC-bound Cdc20 with APC/C is to provide an opportunity for Cdc20 autoubiquitylation by APC/C, a process aided by APC/C subunit Apc15 (Mansfeld et al., 2011; May et al., 2017; Sewart and Hauf, 2017; Uzunova et al., 2012). Notably, the depletion of *apc15^+^*restores Slp1^Cdc20^ levels in *pek1^DD^* cells (Figure 5A), consistent with the idea that Pmk1-mediated Slp1^Cdc20^ phosphorylation very likely facilitates its autoubiquitylation. Indeed, our biochemical and functional analyses further establish K479 and K472 in Slp1^Cdc20^ as two key residues directly involved in its ubiquitylation, which likely follows its phosphorylation, particularly at adjacent Thr480. Despite K485 and K490 are similarly located at C terminus of human Cdc20 and can be ubiquitylated and responsible for Cdc20 degradation (Danielsen et al., 2011; Mansfeld et al., 2011), there has been no reports on phosphorylation-modified residues in this region. However, although the sequence homology within N- and C-terminal regions between Slp1^Cdc20^ and Cdc20 homologues in higher eukaryotes is extremely low, we do notice that there are several lysine-flanking serine or threonine residues within Cdc20 C-terminal tails of higher eukaryotes (see Figure 5-figure supplement 3), this raises a possibility that this region may be involved in phosphorylation-coupled ubiquitylation. Currently, we do not know how T480 phosphorylation accelerate ubiquitylation of K479 and K472 in Slp1^Cdc20^. It is possible that phosphorylation of Slp1^Cdc20^ might alter the mode of Cdc20 binding to APC/C which becomes conducive to catalyze Slp1^Cdc20^ ubiquitylation.

In summary, our studies revealed a distinct dual mechanism of MAPK-dependent anaphase onset delay imposed directly on APC/C co-activator Slp1^Cdc20^ and an unidentified substrate, which may emerge as an important regulatory layer to fine tune APC/C and MCC activity upon SAC activation. Importantly, although this mechanism of APC/C regulation by two MAPKs has not been examined in other organisms, it is very likely widespread across eukaryotes including humans, given the involvement of MAPKs in general response to wide range of adverse environmental situations and similar regulations of APC/C and SAC in eukaryotes characterized so far. It is also plausible to assume that the MAPK-mediated APC/C inhibition pathway may also contribute to the pathogenesis of multiple cancers. Indeed, Cdc20 upregulation has been demonstrated in several types of cancers, including breast and colon cancer, and it contributes to aggressive tumor progression and poor prognosis in gastric cancer and primary non-small cell lung cancer (Chi et al., 2019). One very recent study conducted in human pancreatic ductal adenocarcinoma (PDAC) cell lines revealed that inhibition of extracellular signal-regulated kinase (ERK) led to dephosphorylation of Cdc20 and concomitant loss of APC/C target proteins securin and cyclin B (CCNB1 and CCNB2) (Klomp et al., 2024). Because human ERKs are homologs of fission yeast Pmk1, the finding in this study supports a notion that very likely human MAPKs also negatively regulates APC/C through Cdc20 phosphorylation and the altered levels of Cdc20 in cancers are MAPK relevant. Nevertheless, inhibition of APC/C^Cdc20^ activity or induction of Cdc20 degradation have already been listed as alternative therapeutic strategies to control cancer (Greil et al., 2022; Jeong et al., 2022). Thus, although the detailed molecular mechanisms may vary, further studies on MAPK-mediated APC/C^Cdc20^ inhibition mechanism in human would lead to better understanding of its oncogenic role in tumor progression.

## Materials and Methods

### Fission yeast strains, media and genetic methods

Fission yeast cells were grown in either YE (yeast extract) rich medium or EMM (Edinburgh minimal medium) containing the necessary supplements. G418 disulfate (Sigma-Aldrich; A1720), hygromycin B (Sangon Biotech; A600230) or nourseothiricin (clonNAT; Werner BioAgents; CAS#96736-11-7) was used at a final concentration of 100 μg/ml and thiabendazole (TBZ) (Sigma-Aldrich; T8904) at 5-15 μg/ml in YE media where appropriate. For serial dilution spot assays, 10-fold dilutions of a mid-log-phase culture were plated on the indicated media and grown for 3 to 5 days at indicated temperatures.

To create strain with an ectopic copy of *slp1^+^* at *lys1^+^* locus on chromosome 1, the open reading frame of *slp1^+^*(1467 bp) and its upstream 1504 bp sequence was first cloned into the vector pUC119-*P_adh21_*-MCS-*hphMX6-lys1*\* with *adh21* promoter (*P_adh21_*) removed using the ‘T-type’ enzyme-free cloning method (Chen et al., 2017). The sequence of *hphMX6* in above plasmid was replaced by fragment corresponding to nourseothiricin resistance gene sequence by *Bgl*II-*EcoR*I cut and re-ligation, generating pUC119-*P_slp1_*-*slp1^+^-T_adh1_::natMX6-lys1*\*. The resultant plasmids were linearized by *Apa*I and integrated into the *lys1^+^*locus, generating the strains *lys1*Δ*::P_slp1_-slp1^+^-T_adh1_::hphMX6* and *lys1*Δ*::P_slp1_-slp1^+^-T_adh1_::nat^R^*.

To generate the mutant strains harboring lysine/arginine (K/R) or serine/threonine to glutamic acid (E) or alanine (A) mutations, or lysine (K) to arginine (R) mutations within Slp1, the desired mutations were introduced into pUC119-*P_slp1_-slp1^+^-T_adh1_::hphMX6* using standard methods of site-directed mutagenesis. To generate the strains carrying superfolder GFP (sfGFP)-tagged wild type or mutant Slp1, a fragment corresponding to the sequence of *sfGFP* was inserted in front of *slp1* in pUC119-*P_slp1_-slp1-T_adh1_::hphMX6-lys1*\* using the ‘T-type’ enzyme-free cloning method. The resulting plasmids were linearized and integrated into the *lys1^+^* locus of chromosome 1 using the *hyg^r^* marker. Then endogenous *slp1^+^* was deleted and replaced with *ura4^+^*.

To generate the strains expressing *ade6::P_adh1_-GST-slp1^(456-488aa)^::nat^R^*, C-terminal sequence of *slp1^+^* corresponding to 456-488aa was first cloned into the vector pUC119-*P_adh1_*-MCS-*natMX6-lys1*\* and the sequence of *lys1*\* was replaced with *ade6^+^* from pAde6-NotI-hphMX (a kind gift of Li-lin Du), and then the sequence of GST from pGEX-4T-1 was inserted in front of *slp1*. The resulting plasmid pUC119-*P_adh1_-GST-slp1^(456-488aa)^*-*natMX6-ade6* was linearized and integrated into the *ade6^+^*locus of chromosome III using the *nat^r^* marker.

To generate the strains expressing *ade6::P_adh1_-slp1^(1-60aa)^-mEGFP-2xNLS-GST-slp1^(456-488aa)^::nat^R^*, sequences corresponding to *slp1^(1-60aa)^*, *mEGFP* and two tandem SV40 NLS (CCT AAG AAA AAA CGA AAA GTT GAG GAT CCT AAA AAG AAA CGA AAA GTT GAT) were cloned into the plasmid pUC119-*P_adh1_-GST-slp1^(456-488aa)^*-*natMX6-ade6* using the ‘T-type’ enzyme-free cloning method. The resultant plasmid was linearized and integrated into the *ade6^+^*locus as describes above.

To generate the strains expressing *lys1*Δ*::P_adh11_-pmk1-GBP-mCherry::hyg^R^*, the coding sequence of *pmk1^+^* was first cloned into the vector pUC119-*P_adh11_-GBP-mCherry-hphMX6-lys1*\* (Chen et al., 2017), then the plasmid was linearized and integrated into the *lys1^+^* locus in the genome.

To generate the constitutively active MAPKK strains *pek1^DD^*or *wis1^DD^*, the Ser234 and Thr238 of *pek1^+^* in pUC119-*P_adh21_*-*6HA-pek1^+^*-*T_adh1_*-*hphMX6-lys1*\*, pUC119-*P_adh11_*-*6HA-pek1^+^*-*T_adh1_*-*hphMX6-lys1*\* and pUC119-*P_adh11_*-*6HA-pek1^+^*-*T_adh1_*-*kanMX6-ura4^+^* and the Ser469 and Thr473 of *wis1^+^*in pUC119-*P_adh21_*-*6HA-wis1^+^*-*T_adh1_*-*hphMX6-lys1*\*, pUC119-*P_adh11_*-*6HA-wis1^+^*-*T_adh1_*-*hphMX6-lys1*\* and pUC119-*P_adh11_*-*6HA-wis1^+^*-*T_adh1_*-*kanMX6-ura4^+^*were changed to aspartic acids (D) by standard methods of site-directed mutagenesis. The resulting plasmids were linearized and integrated into the *lys1^+^* locus of chromosome I or the *ura4^+^*locus of chromosome III using the *kan^r^* marker. A list of the yeast strains used is in Supplementary file 1.

### Fission yeast cell synchronization methods

For *nda3-KM311* strains, cells were grown at the permissive temperature for *nda3-KM311* (30 °C) to mid-log phase, synchronized at S phase by adding HU (Sangon Biotech; A600528) to a final concentration of 12 mM for 2 hours followed by a second dose of HU (6 mM final concentration) for 3.5 hours. HU was then washed out and cells were released at specific temperatures as required by subsequent experiments.

For *cdc25-22* background strains, cells were first grown at 25 °C and then arrested at the G_2_/M transition by shifting to 36 °C for 3.5 hours. To allow cells to progress into mitosis, the cultures were released by shifting back to 25 °C. Progression through cell cycle was monitored by DAPI staining after being fixed and counting binucleate cells.

### Spindle checkpoint activation and silencing assays in fission yeast

For checkpoint silencing assay in the absence of microtubules, mid-log *cdc13-GFP nda3-KM311* cells were first synchronized with HU at 30 °C and then arrested in early mitosis by shifting to 18 °C for 6 hours, followed by incubation at 30 °C to allow spindle reformation and therefore spindle checkpoint inactivation. For osmotic stress or cell wall damage stress experiments, 0.6 M KCl or 2 μg/mL caspofungin was added into cultures during 18 °C incubation 1 hr before being shifted to 30 °C. Cells were withdrawn at certain time intervals and fixed with cold methanol and stained with DAPI. 200-300 cells were analyzed for each time point. Each experiment was repeated at least three times.

To achieve spindle checkpoint activation and metaphase arrest through a different mechanism, Mad2 was overexpressed from the *nmt1* promotor (i.e. *P_nmt1_*). To induce Mad2 overexpression, strains carrying *P_nmt1_-mad2::leu1^+^*were pre-cultured in minimal medium with supplements (EMM5S) and 15 µM thiamine, cells were washed 5 times in EMM5S to remove thiamine before being grown in EMM5S at 30 °C for 18 hours. Mitotic arrest was determined by DAPI staining and GFP-Atb2-labeled short spindles.

### Immunoblotting and immunoprecipitation

For routine Western blot and immunoprecipitation experiments, yeast cells were collected from *nda3-KM311* arrested and released cultures, followed by lysing with glass bead disruption using Bioprep-24 homogenizer (ALLSHENG Instruments, Hangzhou, China) in NP40 lysis buffer (6 mM Na_2_HPO_4_, 4 mM NaH_2_PO4, 1% NP-40, 150 mM NaCl, 2 mM EDTA, 50 mM NaF, 0.1 mM Na_3_VO_4_) plus protease inhibitors as previously described (Wang et al., 2012). Proteins were immunoprecipitated by IgG Sepharose beads (GE Healthcare; 17-0969-01) (for Lid1-TAP) or anti-GFP nanobody agarose beads (AlpalifeBio, Shenzhen, China; KTSM1301) (for Mad3-GFP). Immunoblot analysis of cell lysates and immunoprecipitates was performed using appropriate antibodies at 1:500 - 1:5000 dilutions and was read out using chemiluminescence.

### Detection of *in vivo* phosphorylation of Slp1^Cdc20^ at Thr480 and Ser28/Thr31

For detection of phosphorylation of Slp1^Cdc20^ at Thr480 *in vivo*, *mts3-1* mutant strains expressing *ade6::P_adh1_-GST-slp1(456-488aa)::natR* in wild type, *pmk1*Δ and *lys1*Δ*::P_adh11_-6xHA-pek1^DD^*background were arrested at 36 [for 3.5 hours, followed by pull-down of GST-Slp1(456-488aa) by glutathione Sepharose 4B (GE Healthcare). Purified GST-Slp1(456-488aa) was detected with either goat anti-GST HRP-conjugated antibodies (GE Healthcare) or rabbit polyclonal anti-pThr480 antibodies.

For detection of phosphorylation of Slp1^Cdc20^ at Ser28/Thr31 *in vivo*, *nda3-KM311 apc15-13myc::kanR* strains expressing *slp1^+^* or *slp1(S28A;T31A)* in wild type or *cdc2-asM17* background were first synchronized with HU at 30 °C and then arrested in early mitosis by shifting to 18 °C for 6 hours. For *cdc2-asM17* strains, 1 μM 1-NM-PP1 (Toronto Research Chemicals; A603003) was added into one of the cultures to inactivate Cdc2 (Cdk1) 20 min before being collected. Apc15-13myc was immunoprecipitated by anti-c-Myc magnetic beads (MedChemExpress (MCE); HY-K0206). To remove Slp1^Cdc20^ phosphorylation, one of the immunoprecipitated samples was treated with 400 units of Lambda protein phosphatase (New England Biolabs, P0753S) at 30 °C for 30 min. Co-immunoprecipitated Slp1^Cdc20^ was detected with rabbit polyclonal anti-Slp1 antibodies, and the phosphorylated form of Slp1^Cdc20^ was detected with rabbit polyclonal anti-pSer28/pThr31 antibodies respectively.

### Purification of APC/C and analyses by mass spectrometry

To prepare APC/C for mass spectrometry analyses, Apc15-GFP was purified from 6-liter cultures of *nda3-KM311* arrested cells with desired genetic background. Cells were disrupted and cell lysates were prepared as described above for routine immunoprecipitation experiments.

For mass spectrometry analyses, purified samples were first run on PAGE gels, after staining of gels with Coomassie blue, excised gel segments were subjected to in-gel trypsin (Promega, V5111) digestion and dried. Samples were then analyzed on a nanoElute (plug-in V1.1.0.27; Bruker, Bremen, Germany) coupled to a timsTOF Pro (Bruker, Bremen, Germany) equipped with a CaptiveSpray source. Peptides were separated on a 15cm X 75μm analytical column, 1.6 μm C18 beads with a packed emitter tip (IonOpticks, Australia). The column temperature was maintained at 55 °C using an integrated column oven (Sonation GmbH, Germany). The column was equilibrated using 4 column volumes before loading sample in 100% buffer A (99.9% MilliQ water, 0.1% FA) (Both steps performed at 980 bar). Samples were separated at 400 nL/min using a linear gradient from 2% to 25% buffer B (99.9% ACN, 0.1% FA) over 90 min before ramping to 37% buffer B (10min), ramp to 80% buffer B (10min) and sustained for 10 min (total separation method time 120 min). The timsTOF Pro (Bruker, Bremen, Germany) was operated in PASEF mode using Compass Hystar 6.0. Mass Range 100 to 1700m/z, 1/K0 Start 0.6 V⋅s/cm^2^ End 1.6 V⋅s/cm^2^, Ramp time 110.1ms, Lock Duty Cycle to 100%, Capillary Voltage 1600V, Dry Gas 3 L/min, Dry Temp 180°C, PASEF settings: 10 MS/MS scans (total cycle time 1.27sec), charge range 0-5, active exclusion for 0.4 min, Scheduling Target intensity 10000, Intensity threshold 2500, CID collision energy 42eV. All raw files were analyzed by PEAKS Studio Xpro software (Bioinformatics Solutions Inc., Waterloo, ON, Canada). Data was searched against the *S. pombe* proteome sequence database (Uniprot database with 5117 entries of protein sequences at https://www.uniprot.org/proteomes/UP000002485). *De novo* sequencing of peptides, database search and characterizing specific PTMs were used to analyze the raw data; false discovery rate (FDR) was set to ≤1%, and [-10*log(*p*)] was calculated accordingly where *p* is the probability that an observed match is a random event. The PEAKS used the following parameters: (i) precursor ion mass tolerance, 20 ppm; (ii) fragment ion mass tolerance, 0.05 Da (the error tolerance); (iii) tryptic enzyme specificity with two missed cleavages allowed; (iv) monoisotopic precursor mass and fragment ion mass; (v) a fixed modification of cysteine carbamidomethylation; and (vi) variable modifications including N-acetylation of proteins and oxidation of Met.

### Ubiquitin affinity pull-down assays

For detection of *in vivo* ubiquitylation of Slp1^Cdc20^, strains all carried *mts3-1* mutation were used. sfGFP-tagged wild type Slp1 or Slp1 mutants were expressed in the genomically integrated form of *P_slp1_-sfGFP-slp1(WT or mutants)::hyg^R^* in wild type, *pmk1*Δ or *P_adh11_-6xHA-pek1^DD^*background. Cultures were first grown at 25 °C to mid-log phase and then shifted to 36 [for 3.5 hours to block cells in mitosis prior to harvesting and cell lysis. Ubiquitinated proteins were pulled down from yeast lysates using Tandem Ubiquitin Binding Entities (TUBEs) (LifeSensors, UM402) according to the manufacturer’s instructions. The bound proteins were eluted from washed Agarose-TUBE2 beads and analyzed by western blotting using a mouse monoclonal anti-GFP antibody (Beijing Ray Antibody Biotech; RM1008). Cell extracts were also immunoblotted for GFP and Cdc2 as loading controls (inputs).

### Expression and purification of recombinant proteins

GST or MBP fusion constructs of fission yeast full-length or truncated Slp1^Cdc20^, Pmk1, Pek1 and Atf1 were generated by PCR of the corresponding gene fragments from yeast genomic DNA or cDNA and cloned in frame into expression vectors pGEX-4T-1 (GE Healthcare) or pMAL-c2x (New England Biolabs), respectively. Site-directed mutagenesis was done by standard methods to generate vectors carrying Pek1-DD(S234D,[T238D) or Slp1^Cdc20^ harboring K(Lys)/R(Arg) to E(Glu) or A(Ala) mutations at individual or combined sites including K19, K20, R21, K47, R48, K140, K141, K161, R162, R163, K472, R473. Integrity of cloned DNA was verified by sequencing analysis. All recombinant GST- or MBP-fusion proteins were expressed in *Escherichia coli* BL21 (DE3) cells. The recombinant proteins were purified by glutathione Sepharose 4B (GE Healthcare) or Amylose Resin High Flow (New England Biolabs) respectively according to the manufacturers’ instructions.

### *In vitro* pulldown assay

The recombinant MBP-Slp1^Cdc20^ and MBP, or GST-Pmk1 and GST proteins were produced in bacteria and purified as described above. To detect the interaction between bacterially expressed Slp1^Cdc20^ and Pmk1-HA expressed in yeast, about 1 μg of MBP-Slp1^Cdc20^ or MBP immobilized on amylose resin was incubated with cleared cell lysates prepared from yeast strains *nda3-KM311 pmk1-6His-HA*, which were HU synchronized and arrested at prometaphase by being grown at 18 °C for 6 hours. To detect the direct interaction between bacterially expressed Slp1 and Pmk1, about 1 μg of GST-Pmk1 or GST immobilized on glutathione Sepharose 4B resin was incubated with cleared bacterial lysates expressing MBP-Slp1^Cdc20^. Samples were incubated for 1-3 hours at 4 °C. Resins were washed 3 times with lysis buffer (6 mM Na_2_HPO_4_, 4 mM NaH_2_PO_4_, 1% NP-40, 150 mM NaCl, 2 mM EDTA, 50 mM NaF, 0.1 mM Na_3_VO_4_, plus protease inhibitors), suspended in SDS sample buffer, and then subject to SDS-PAGE electrophoresis and Coomassie brilliant blue (CBB) staining.

### Non-radioactive *in vitro* kinase assay

GST-Slp1^Cdc20^, GST-Atf1, GST-Pmk1, and GST-Pek1^DD^ fusions were expressed and purified from *Escherichia coli* with glutathione Sepharose 4B as described above. Pmk1-HA-His and Cdc13-GFP were purified from yeast lysates by immuprecipitation with anti-HA or anti-GFP antibodies respectively.

For detection of phosphorylation of Slp1^Cdc20^ and Atf1 by yeast derived Pmk1, glutathione Sepharose 4B beads bound with GST-Slp1^Cdc20^ or GST-Atf1 and agarose beads bound with Pmk1-HA-His were mixed and washed 3 times with kinase buffer (50 mM Tris-HCl, pH 7.5, 10 mM MgCl_2_, 1 mM EGTA), then the reactions were incubated in kinase buffer with 20 μM ATPγS (Sigma-Aldrich; A1388) at 30°C for 45[min. The kinase reactions were stopped by adding 20 mM EDTA, and the reaction mixture was alkylated after incubation at room temperature with 2.5[mM *p*-nitrobenzyl mesylate (PNBM) (Abcam; ab138910) for 1 hr. The alkylation reaction was stopped by boiling in SDS-PAGE loading buffer. Phosphorylated proteins were detected with rabbit monoclonal anti-Thiophosphate ester antibody (Abcam, ab239919).

For detection of phosphorylation of Slp1^Cdc20^ at Thr480 by bacterially expressed Pmk1, beads bound with GST-Pmk1, GST-Pek1(DD) and fused substrates GST-Slp1^(456-488aa)^ or GST-Slp1^(456-488aa, T480A)^ were washed and incubated in kinase buffer (50 mM Tris-HCl, pH 7.5, 10 mM MgCl_2_, 1 mM EGTA) with 10 mM ATP (Sangon Biotech; A600020-0005) at 30°C for 45 min. The reaction was stopped by boiling in SDS-PAGE loading buffer. Phosphorylated proteins were detected with rabbit polyclonal anti-pThr480 antibodies.

For detection of phosphorylation of Slp1^Cdc20^ at Ser28/Thr31 by Cdk1, beads bound with bacterially expressed MBP-Slp1^(1-190aa)^ or MBP-Slp1^(1-190aa,S28A;T31A)^ and agarose beads bound with Cdc13-GFP/Cdc2 from yeast lysate were mixed and washed 3 times with kinase buffer (10 mM Tris-HCl, pH 7.4, 10 mM MgCl_2_, 1 mM DTT), then the reactions were incubated in kinase buffer with 10 mM ATP (Sangon Biotech; A600020-0005) at 30°C for 45 min. To inactivate Cdc2-as1, 5 μM 1-NM-PP1 (Toronto Research Chemicals; A603003) was added into the reaction. The reaction was stopped by boiling in SDS-PAGE loading buffer. Phosphorylated proteins were detected with rabbit polyclonal anti-pSer28/pThr31 antibodies.

### Antibodies for immunoblotting

The phospho-site specific antibodies were made in an in-house facility (for anti-Slp1-pS28/pT31) or by ABclonal Technology Co., Ltd. (Wuhan, China) (for anti-Slp1-pT480) by immunizing rabbits with pS28/pT31- or pT480-containing peptides (LVFPN(S-p)PI(T-p)PLHQQ or PITK(T-p)PSSS, respectively) coupled to haemocyanin (Sigma). The purified antibodies were concentrated to 0.2 mg/mL and used at 1:100 dilution. The following antibodies used for immunoblot analyses were purchased from the indicated commercial sources and were used at the indicated dilution: peroxidase-anti-peroxidase (PAP) soluble complex (Sigma-Aldrich; P1291; 1:10,000); rabbit polyclonal anti-Myc (GeneScript; A00172-40; 1:2,000); mouse monoclonal anti-GFP (Beijing Ray Antibody Biotech; RM1008; 1:2,000); rat monoclonal anti-HA (Roche, Cat. No. 11 867 423 001; 1:2,000); rabbit polyclonal anti-Slp1 (Kim et al., 1998) (1:500); rabbit polyclonal anti-phospho-p44/42 (detecting activated Pmk1) (Cell Signaling Technology; #9101; 1:1,000); rabbit monoclonal anti-phospho-p38 (Cell Signaling Technology; #4511; 1:1,000); rabbit polyclonal anti-PSTAIRE (detecting Cdc2) (Santa Cruz Biotechnology; sc-53; 1:1,000); rabbit monoclonal anti-Thiophosphate ester antibody (Abcam; ab239919; 1:5,000); goat anti-GST HRP-conjugated antibody (RRID:AB_771429; GE Healthcare; 1:10,000). Secondary antibodies used were goat anti-mouse or goat anti-rabbit polyclonal IgG (H+L) HRP conjugates (Thermo Fisher Scientific; #31430 or #32460; 1:10,000).

### RT-qPCR

For RT-qPCR of *slp1^+^* mRNA, *nda3-KM311* strains were grown, synchronized by HU and arrested at metaphase with SAC being activated as described above. Cells were collected and total cellular RNA was isolated by using TRIZOL method. cDNA was prepared using oligo(dT) and PrimeScript reverse transcriptase (Takara, D2680A). Real-time PCR was performed in the presence of SYBR Green using an Applied Biosystems 7900HT light cycler. Relative RNA levels were calculated using the Δ*CT* method and normalized to *act1^+^* levels.

### Fluorescence microscopy

Cdc13-GFP proteins were observed in cells after fixation with cold methanol. For DAPI staining of nuclei, cells were fixed with cold methanol, washed in PBS and resuspended in PBS plus 1 μg/ml DAPI. Photomicrographs of cells were obtained using a Nikon 80i fluorescence microscope coupled to a cooled CCD camera (Hamamatsu, ORCA-ER) or a Perkin Elmer spinning-disk confocal microscope (UltraVIEW^®^ VoX) with a 100x NA 1.49 TIRF oil immersion objective (Nikon) coupled to a cooled CCD camera (9100-50 EMCCD; Hamamatsu Photonics) and spinning disk head (CSU-X1, Yokogawa). Image processing and analysis were carried out using Element software (Nikon), ImageJ software (National Institutes of Health) and Adobe Photoshop.

### Statistical analyses and reproducibility

No statistical methods were used to predetermine sample size. Sample size was as large as practicable for all experiments. The experiments were randomized and the investigators were blinded to group allocation during experiments but not blinded to outcome assessment and data analysis.

All experiments were independently repeated two to more than three times with similar results obtained. For quantitative analyses of each time course experiment, ≥300 cells were counted for each time point or sample. The same sample was not measured or counted repeatedly. No data were excluded from our studies. Data collection and statistical analyses were performed using Microsoft Office Excel or GraphPad Prism 6 softwares. Data are expressed as mean values with error bars (±[standard deviation/s.d.) and were compared using two-tailed Student’s *t*-tests unless indicated otherwise.

## Acknowledgments

We thank Kathy Gould, Li-lin Du, Silke Hauf, Jonathan Millar, Takashi Toda, Yoshinori Watanabe, Elena Hidalgo and National BioResource Project (NBRP), Japan (http://yeast.nig.ac.jp/yeast/) for fission yeast strains or plasmids; Rafael Daga for communication on unpublished results; Jose Cansado for advice on active MAPK detection. We also thank Yaying Wu, Zheni Xu and Chang-chuan Xie for mass spectrometry experiments and data analyses; Yu-ting He for contributing a few time course experimental data; and Shiyu Huang and Jun-ming Zhuang for help with construction of some plasmids or yeast strains.

## Author Contributions Statement

W.S., J.C., and C.J. and R.W. performed all microscopy experiments with the help from Z.L.. L.S., X.C., C.S., W.S., L.L., S.B. and S.W. performed all immunoblotting, co-immunoprecipitation experiments, *in vitro* binding assays and *in vitro* kinase aassays. S.W. and X.W. performed large-scale protein purifications. X.C. and S.W. carried out genetic analyses. Q.J. and Y.W. conceived the study. Q.J. acquired funding. Y.W. and Q.J. designed the experiments and supervised the research. Y.W. and Q.J. wrote the original draft with the inputs from Z.L.. R.W., Y.W. and Q.J. reviewed and edited the revised manuscript.

## Data availability statement

The authors confirm that all data supporting the findings of this study are available within the manuscript main figures, supplementary figures, supplementary tables and source data files. The mass spectrometry proteomics data have been deposited to the ProteomeXchange Consortium via the PRIDE partner repository with the dataset identifier PXD048800.

## Supplemental Figure legends

**Figure 1-figure supplement 1.**
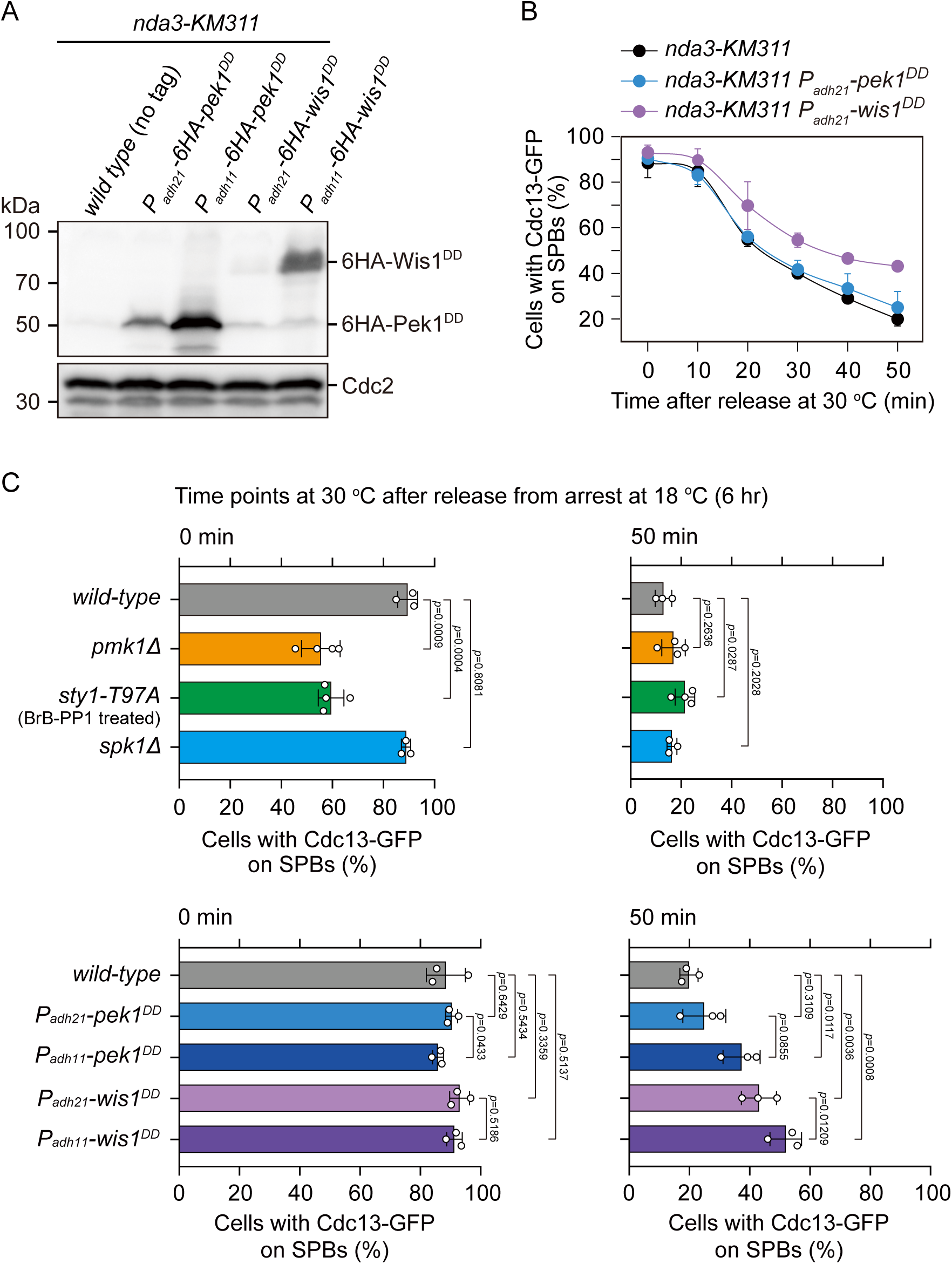
Examination of protein levels of overexpressed Pek1^DD^ and Wis1^DD^ and analyses of spindle checkpoint inactivation efficiency in *P_adh21_*-*pek1^DD^*and *P_adh21_*-*wis1^DD^* mutants. (**A**) Immunoblot analysis of the constitutively overexpressed Pek1^DD^ and Wis1^DD^. Cells with indicated genotypes were grown at the permissive temperature for *nda3-KM311* (30 °C) to mid-log phase and then arrested at 18 °C for 6 hours. Protein samples were prepared and immunoblotting was performed with anti-HA and anti-Cdc2 antibodies to detect total 6HA-Pek1^DD^, 6HA-Wis1^DD^ and Cdc2, respectively. Note *P_adh11_* is a stronger version of *P_adh21_* promoter. (**B**) Time-course analyses of SAC activation and inactivation in *nda3-KM311 cdc13-GFP* strains with indicated genotypes. Compared with *P_adh11_*-*pek1^DD^*or *P_adh11_*-*wis1^DD^* mutants, *P_adh21_*-*pek1^DD^* or *P_adh21_*-*wis1^DD^* mutants showed weaker defect in spindle checkpoint inactivation efficiency respectively. n ≥ 3. For each time point, ≥300 cells were counted for every sample. (**C**) Quantification of spindle checkpoint inactivation rate in MAPK-deficient or constitutively active mutants at 0 and 50 min after release at 30 °C from *nda3*-mediated arrest. Cells of indicated strains bearing Cdc13-GFP were grown at 30 °C to mid-log phase and arrested at 18 °C for 6 hours, and then released at 30 °C. For each time point, ≥300 cells were counted for every sample. Data from time points of 0 and 50 min after release were subjected to statistical analysis. Error bars indicate mean ± standard deviation of three independent experiments. *p* values were calculated against wild-type cells.

**Figure 1-figure supplement 2.**
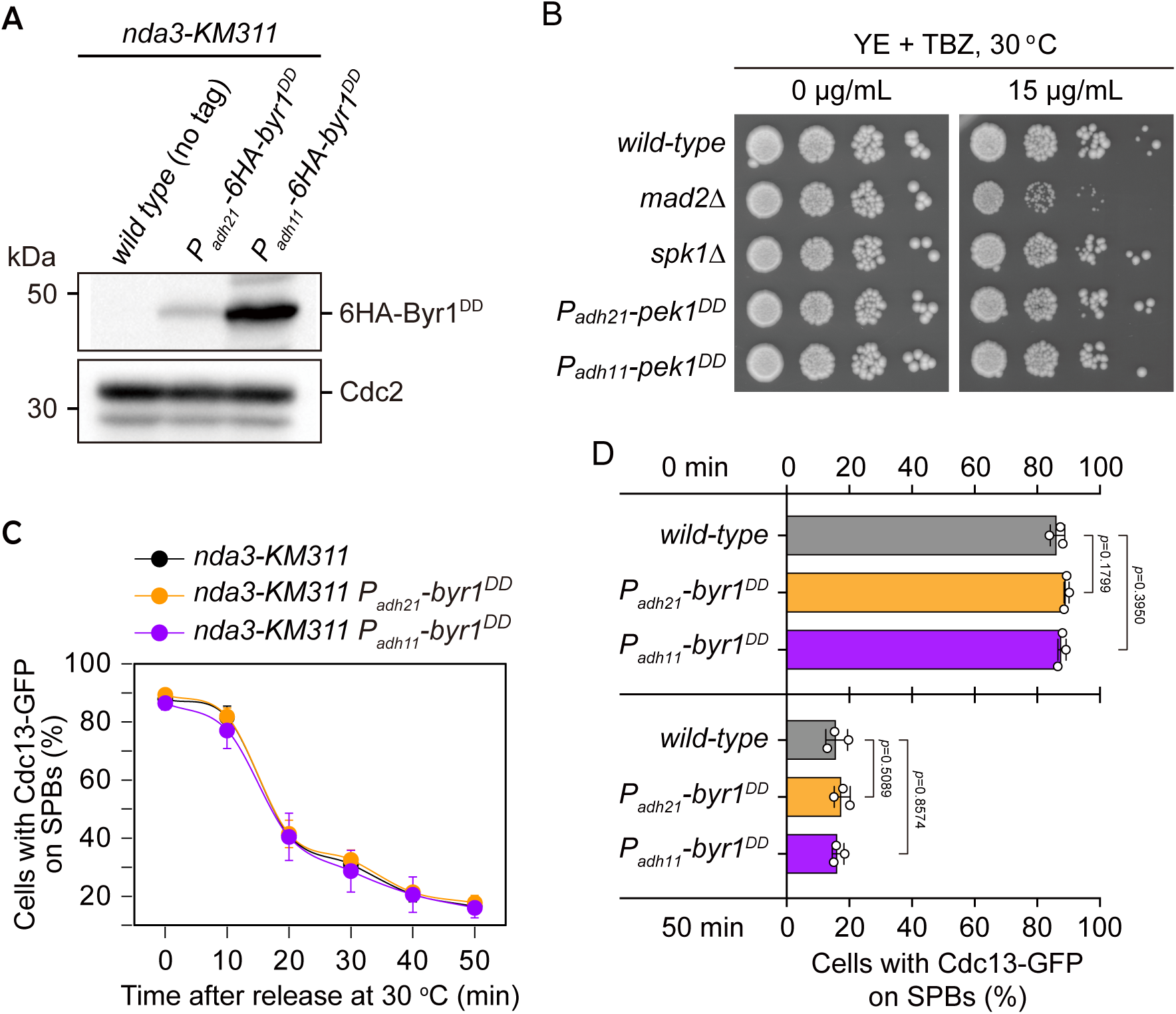
Characterization of possible effect of overexpressed Byr1^DD^ on spindle checkpoint activation and inactivation. **(A)** Immunoblot analysis of the constitutively overexpressed Byr1^DD^. *nda3-KM311* strains with indicated genotypes were first grown at 30 °C to mid-log phase and then arrested at 18 °C for 6 hours. Protein samples were prepared and immunoblotting was performed with anti-HA and anti-Cdc2 antibodies to detect total 6HA-Byr1^DD^ and Cdc2, respectively. Note *P_adh11_* is a stronger version of *P_adh21_* promoter. **(B)** Serial dilution assay on TBZ sensitivity of Byr1^DD^-overexpressing mutants. *mad2*Δ served as a positive control. Note that neither *P_adh21_*-*6HA*-*byr1^DD^* nor *P_adh11_*-*6HA*-*byr1^DD^*mutants were sensitive to TBZ. **(C)** Time-course analyses of SAC activation and inactivation in *nda3-KM311 cdc13-GFP* strains with indicated genotypes. Cells of indicated strains were grown at 30 °C to mid-log phase and arrested at 18 °C for 6 hours, and then released at 30 °C. For each time point, ≥300 cells were counted for every sample. The experiment was repeated 3 times and error bars indicate mean ± standard deviation. **(D)** Quantification of spindle checkpoint inactivation rate in *byr1^DD^* mutants at 0 and 50 min after release at 30 °C from *nda3*-mediated arrest. Data from time points of 0 and 50 min collected in (C) were subjected to statistical analysis. Error bars indicate mean ± standard deviation of three independent experiments. *p* values were calculated against wild-type cells.

**Figure 1-figure supplement 3.**
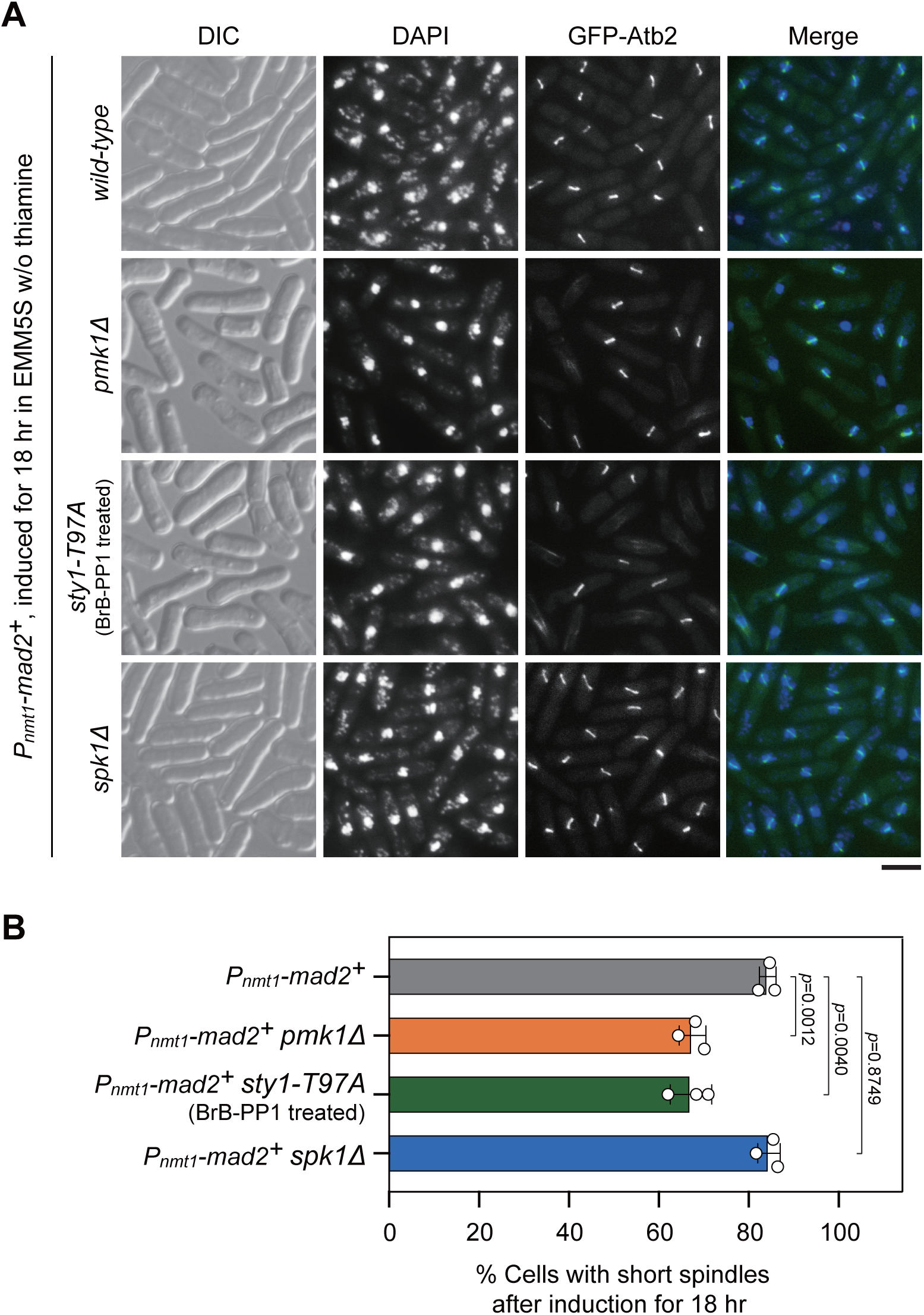
Analyses of metaphase arrest efficiency in *pmk1*Δ, *sty1-T97A* and *spk1*Δ mutants upon Mad2 overexpression. Cultures of *P_nmt1_-mad2^+^ GFP-atb2^+^* strains with indicated genotypes were induced in EMM5S liquid media in the absence of thiamine for 18 hr, and cells were collected, fixed and DAPI stained, and cells with short spindles were scored. For *sty1-T97A* strain, 5μM 3-BrB-PP1 was added at 16 hr after induction. **(A)** Representative images of cells from indicated strains with GFP-Atb2-labeled spindles. DIC, differential interference contrast microscopy. Scale bar, 5 μm. **(B)** Quantitative analyses of cells with short spindles in response to Mad2 overexpression. The experiment was repeated 3 times with >280 cells counted for each strain, and the data are plotted as mean ± standard deviation. Two-tailed unpaired *t*-test was used to derive *p* values.

**Figure 2-figure supplement 1.**
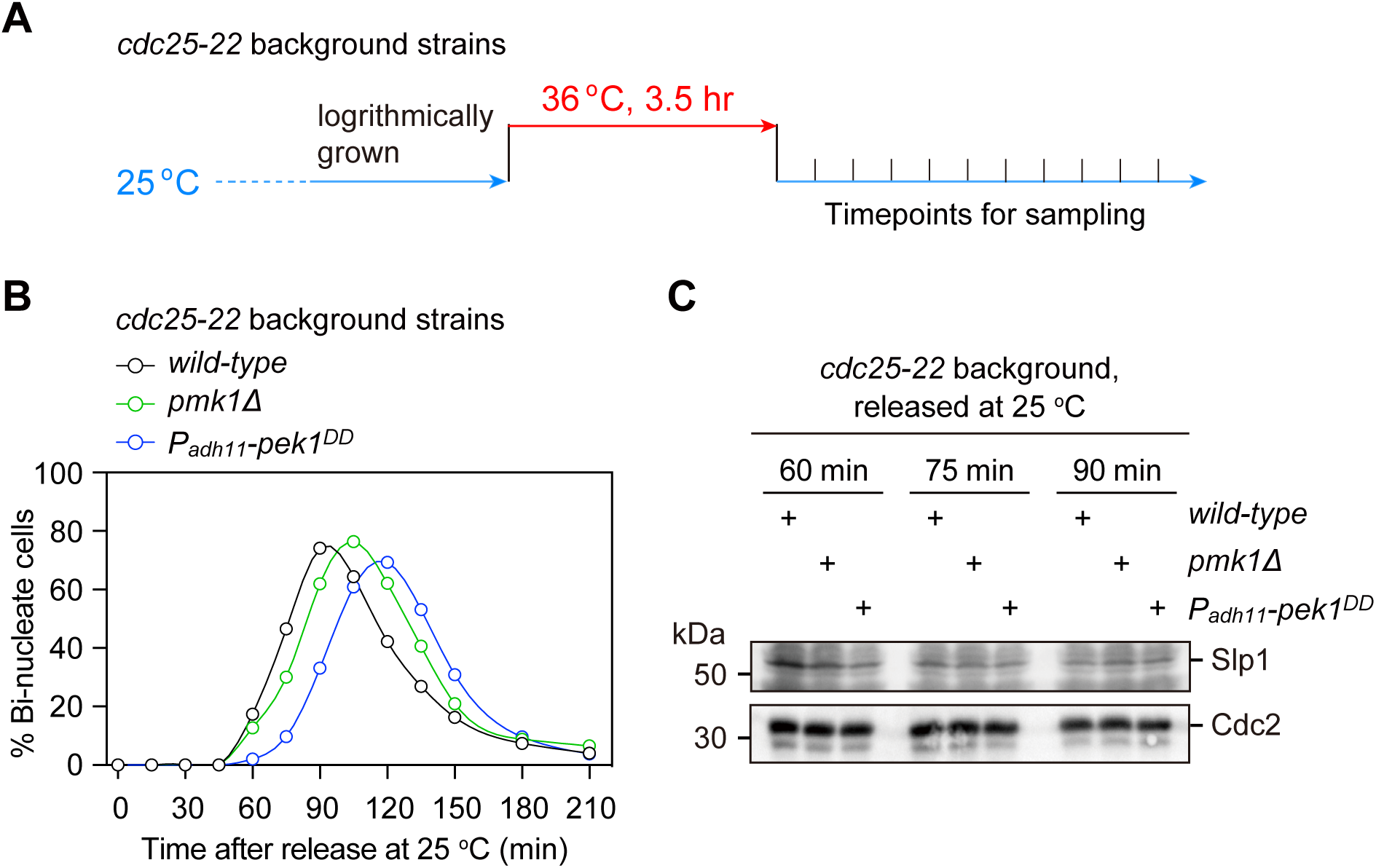
Examination of protein levels of Slp1^Cdc20^ in normally growing mitotic *pmk1Δ* or *pek1^DD^* cells. **(A)** Schematic depiction of the arrest-and-release experiment design. For *cdc25-22* background strains, cells were grown at 25 °C and then arrested at the G_2_/M transition by shifting to 36 °C for 3.5 hours. The cultures were released by shifting back to 25 °C, and aliquots were taken at different time intervals and were fixed and stained with DAPI to monitor cell cycle progression. **(B)** The percentages of binucleated cells were counted based on DAPI staining for each time point after release at 25 °C. **(C)** Samples of indicated strains taken at 60, 75 and 90 min after being shifted back from 36 °C to 25 °C were subjected to immunoblotting with anti-Slp1 and anti-PSTAIR antibodies to detect total Slp1 and Cdc2, respectively. Blots shown are representative of two independent biological replicates.

**Figure 3-figure supplement 1.**
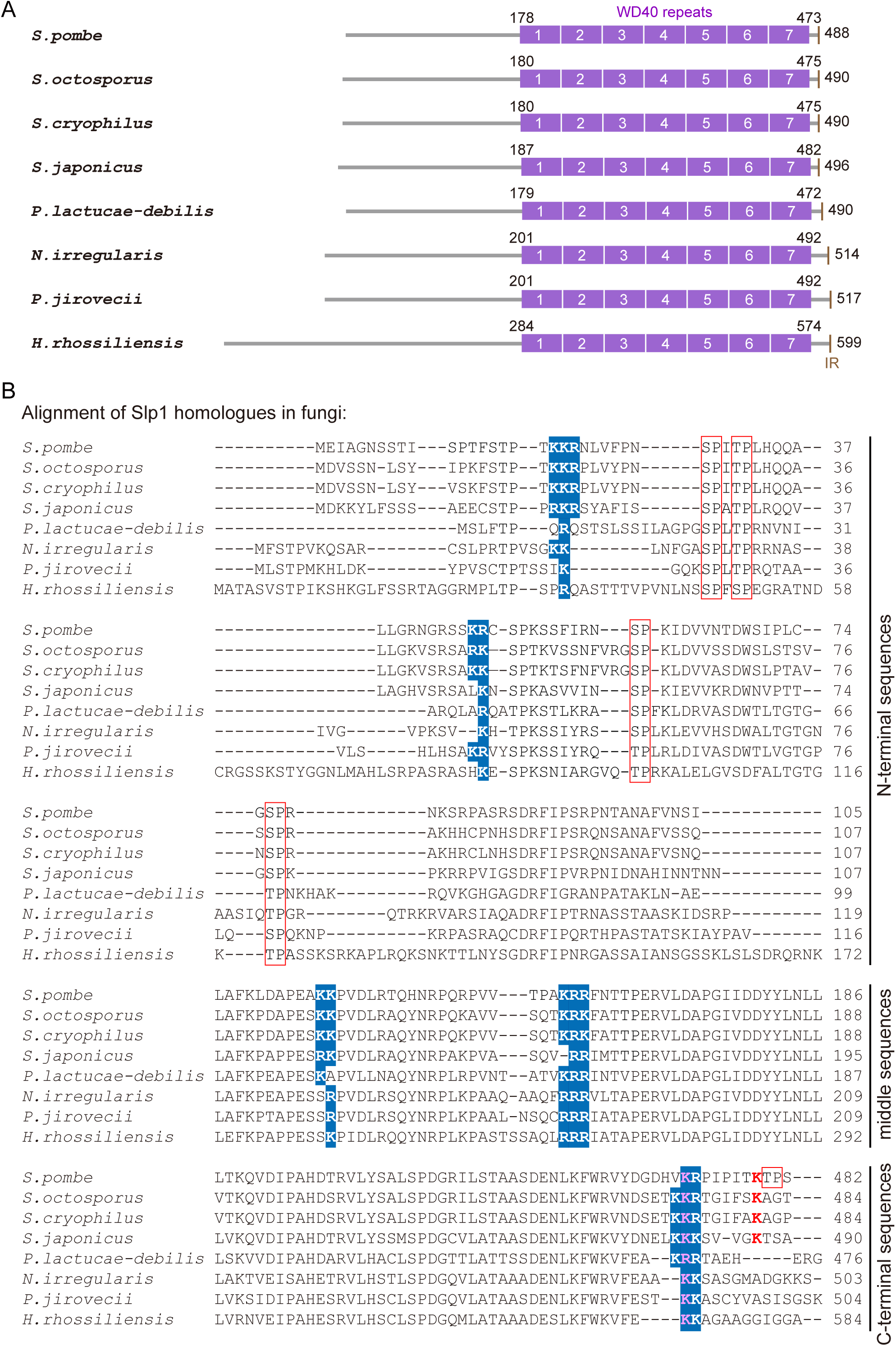
Conservation of basic-residue patches and phosphorylation or ubiquitylation sites within Slp1 homologues in fungi species. (**A**) Schematic depiction of the *S. pombe* (*Schizosaccharomyces pombe*) Slp1 protein with its homologues in 7 other fungi species: *Schizosaccharomyces octosporus*, *Schizosaccharomyces cryophilus*, *Schizosaccharomyces japonicas*, *Protomyces lactucae-debilis*, *Neolecta irregularis*, *Pneumocystis jirovecii* and *Hirsutella rhossiliensis.* Positions of amino acids corresponding to seven WD40 repeats and IR (isoleucine-arginine) motif are indicated. (**B**) Local sequence alignment performed with *S. pombe* Slp1 and its homologue sequences from 7 other fungi species. 5 potential basic-residue patches within Slp1 and their conserved positions in other species are highlighted in blue. Phosphorylated or ubiquitylated residues detected by mass spectrometry in *S. pombe* Slp1 together with their corresponding residues in homologues from 7 other fungi species are labeled with red frames or red letters, respectively. Lysine 472 (K472) in *S. pombe* Slp1 and its corresponding residues in homologues from 7 other fungi species are also indicated in pink. Note that only aligned local sequences (N-terminal, middle and C-terminal portions) of Slp1 homologues harboring the basic-residue patches are shown.

**Figure 3-figure supplement 2.**
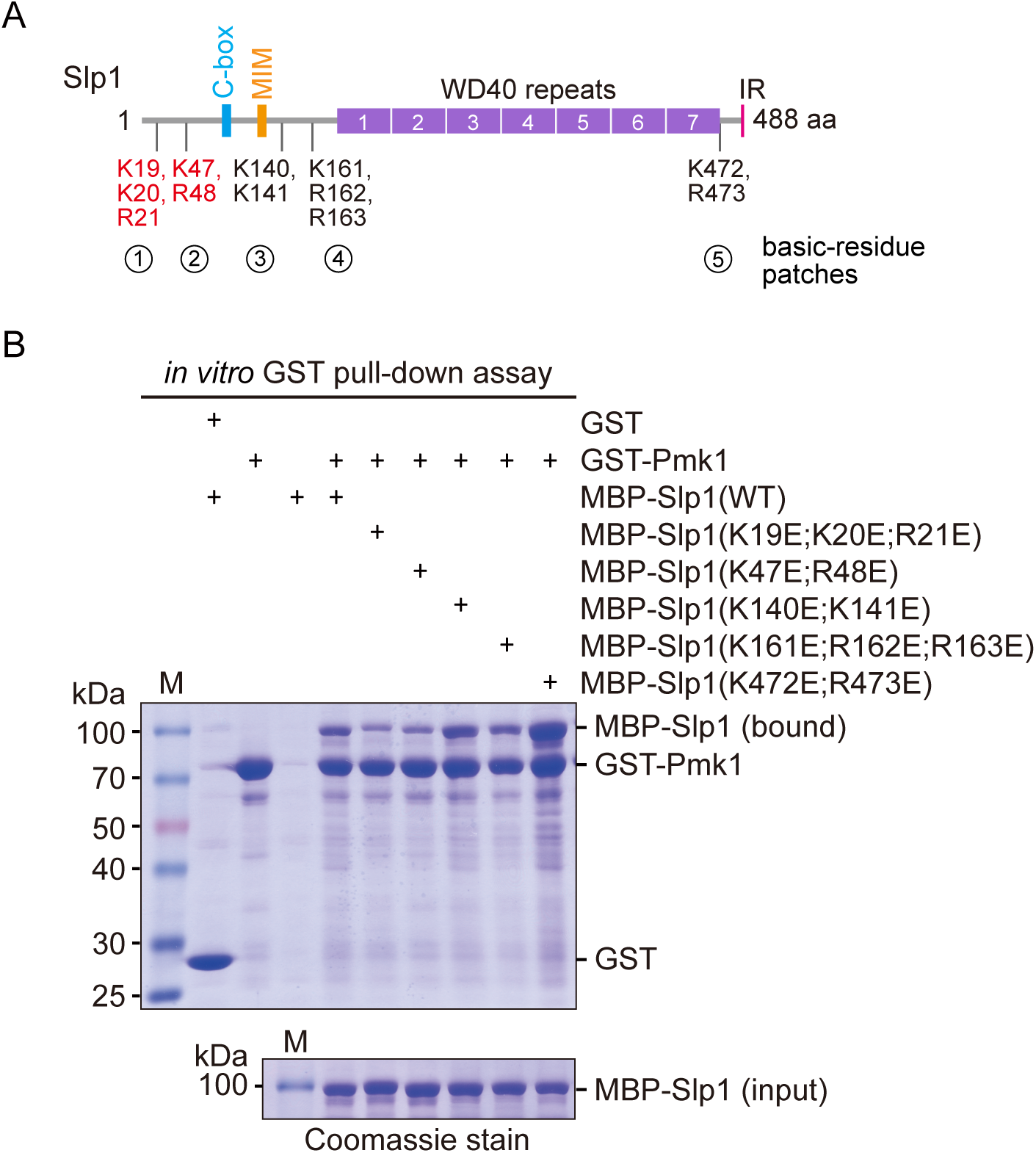
Screen for basic-residue patches in Slp1 mediating its direct interaction with Pmk1 by *in vitro* binding assay. (**A**) Schematic depiction of the *S. pombe* Slp1 protein with five basic-residue patches potentially required for Slp1-Pmk1 association indicated, and two confirmed interaction-mediating patches highlighted in red. MIM, Mad2-interaction motif; IR, isoleucine-arginine tail. (**B**) *In vitro* GST pull-down assays were performed with bacterially expressed GST-Pmk1 and MBP-Slp1 with wild-type Slp1 or mutants harboring lysine/arginine (K/R) to glutamic acid (E) mutations of basic-residue patches within Slp1. Proteins bound on GST beads were separated on SDS-PAGE and visualized by Coomassie blue staining. Note that only Slp1 harboring clustered mutations K19E, K20E and R21E, or K47E and R48E within two most N-terminal basic-residue patches strongly compromised interaction between Slp1 and Pmk1.

**Figure 3-figure supplement 3.**
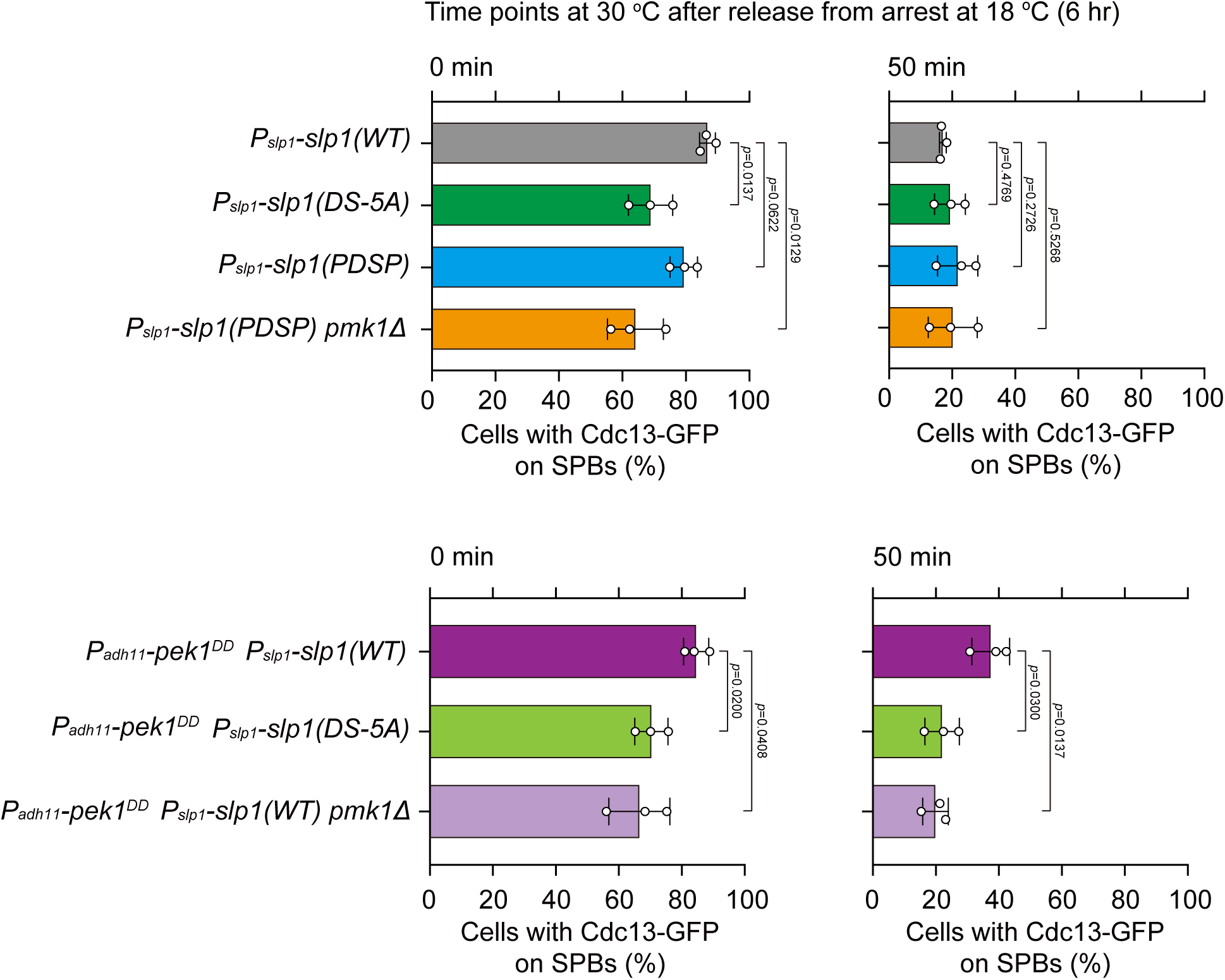
Quantification of spindle checkpoint inactivation rate in *slp1* mutants with Pmk1-docking site mutations at 0 and 50 min after release at 30 °C from *nda3*-mediated arrest. Cells of indicated strains bearing Cdc13-GFP were grown at 30 °C to mid-log phase and arrested at 18 °C for 6 hours, and then released at 30 °C. For each time point, ≥300 cells were counted for every sample. Data from time points of 0 and 50 min after release in Figure 3E were subjected to statistical analysis. Error bars indicate mean ± standard deviation of three independent experiments. *p* values were calculated against wild-type cells.

**Figure 4-figure supplement 1.**
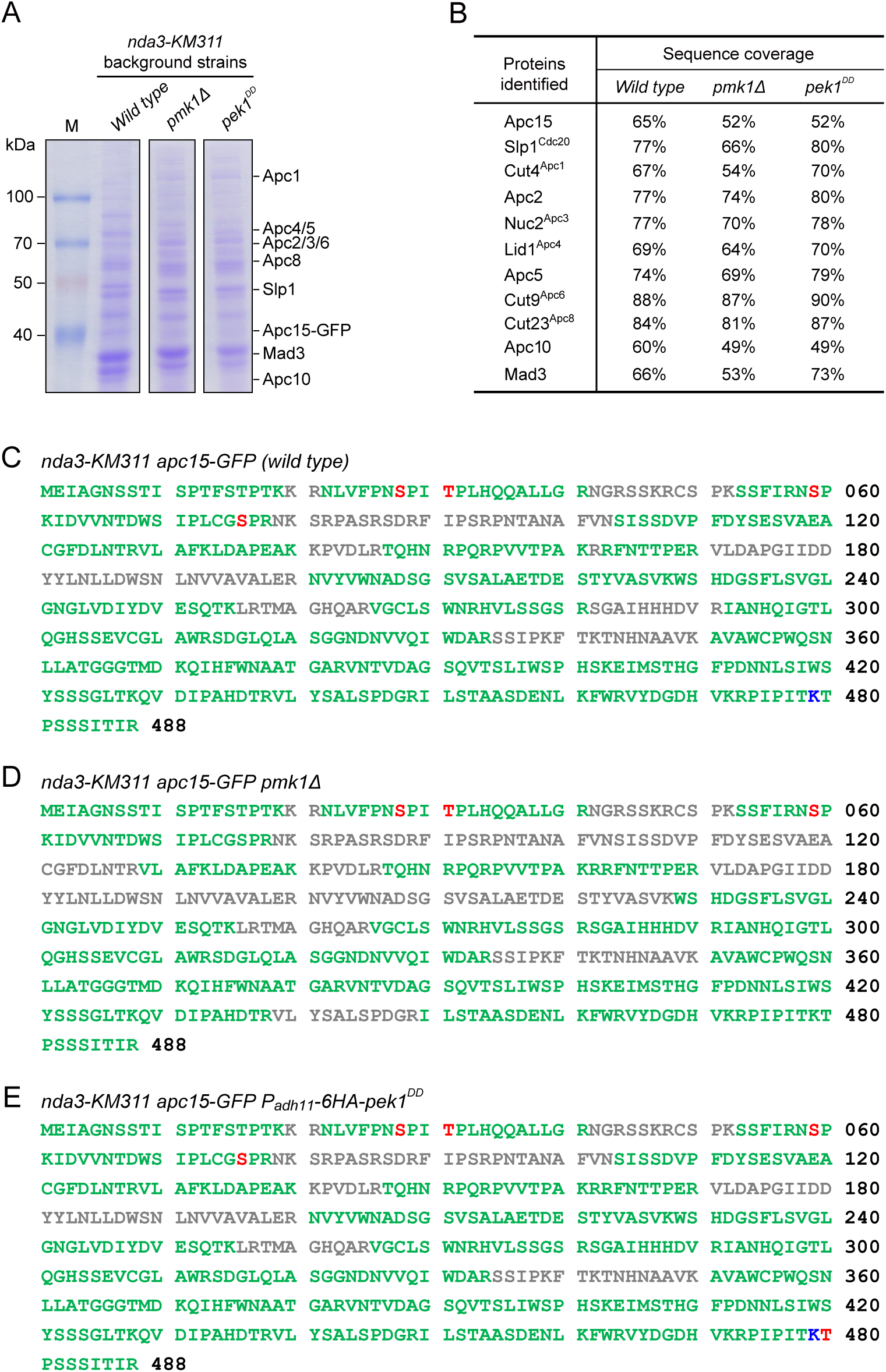
Identification of *in vivo* phosphorylated or ubiquitylated residues in Slp1. (**A, B**) APC/C subunits were isolated by immunoprecipitation of Apc15-GFP from *nda3-KM311* cells arrested at metaphase by being treated at 18 °C for 6 hours. Purified proteins were analyzed by SDS-PAGE and mass spectrometry. Coomassie blue-stained protein gels after SDS-PAGE are shown (A). APC/C subunits or related proteins identified by Apc15-GFP purifications followed by mass spectrometry are listed, and percentages of peptide sequence coverage for each protein are indicated (B). (**C-E**) Slp1 sequences retrieved from 3 purifications in indicated strains with peptide sequence coverage (green), phosphorylated serine or threonine (red), and ubiquitylated lysine (blue). Sequences not covered after mass spectrometry analysis are in gray.

**Figure 4-figure supplement 2.**
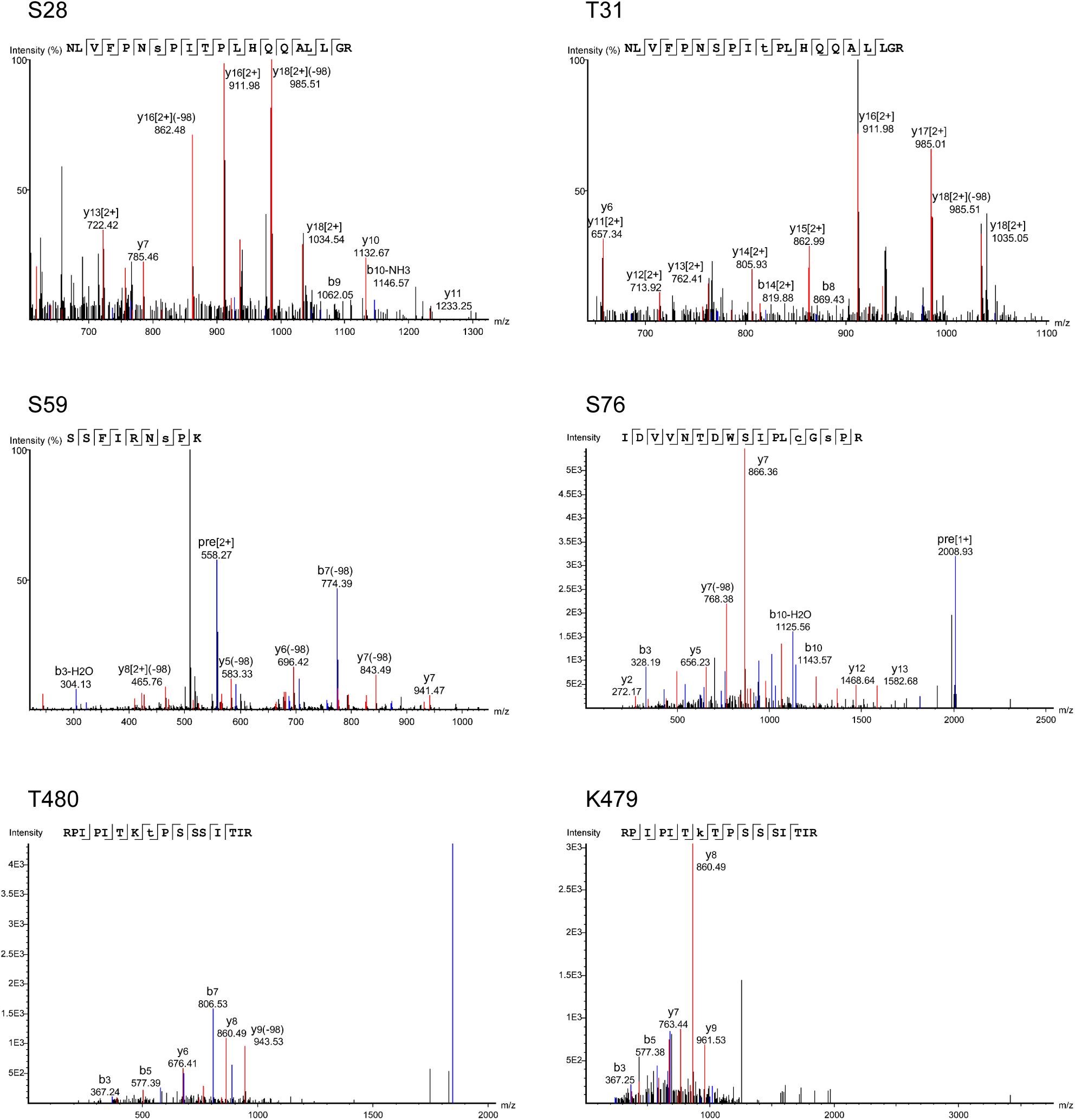
MS spectra from mass spectrometric analyses of Slp1. Examples of spectra for 5 phosphorylation sites (S28, T31, S59, S76, T480) and one ubiquitylation site (K479) identified in Apc15-GFP-assocaited Slp1 purified from metaphase-arrested *P_adh11_-pek1^DD^* cells.

**Figure 4-figure supplement 3.**
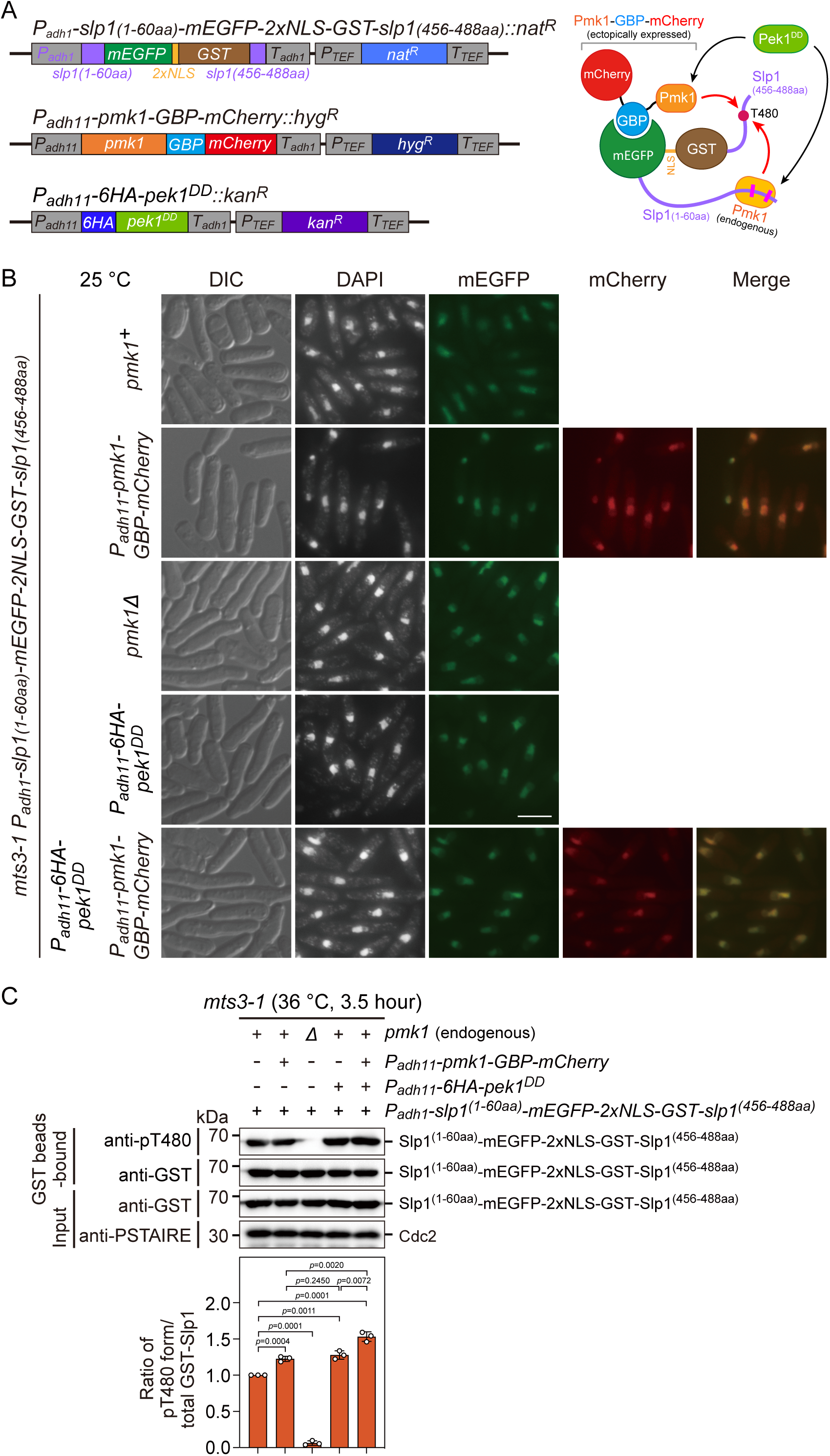
Analysis of Slp1^Cdc20^ phosphorylation at T480 *in vivo* upon forced tethering of Slp1^Cdc20^ to Pmk1. **(A)** Schematic depiction of the experiment design for artificially forced tethering of Slp1^Cdc20^ to Pmk1 using GBP-GFP system. (*Left*) Structures of the genomically integrated *P_adh1_-slp1^(1-60aa)^-mEGFP-2xNLS-GST-slp1^(456-488aa)^::nat^R^*at *ade6^+^* locus, *P_adh11_-pmk1-GBP-mCherry::hyg^R^*at *lys1^+^* locus, and *P_adh11_-6xHA-pek1^DD^::kan^R^*at *ura4^+^* locus. (*Right*) Cartoon depicting the manipulated targeting of mEGFP-2xNLS-GST fusion with Slp1 fragments to GBP-mCherry fusion of Pmk1 mediated by GBP-GFP binding. Note that Thr480 in Slp1 C-terminus could be phosphorylated by either ectopically expressed Pmk1-GBP-mCherry fusion or endogenous Pmk1 activated by Pek1^DD^. **(B)** Representative images of *mts3-1* cells expressing mEGFP-2xNLS-GST, GBP-mCherry or 6HA fusion proteins driven by promoters *P_adh1_* or *P_adh11_*. Cells were grown to early log phase in liquid YE at 25 [, and then collected, fixed, DAPI-stained and visualized by using fluorescence microscopy. DIC, differential interference contrast microscopy. Scale bar, 5 μm. **(C)** Slp1^(1-60aa)^-mEGFP-2xNLS-GST-Slp1^(456-488aa)^ was purified using GST pull-down from *mts3-1* cells with indicated genotypes arrested at 36 [for 3.5 hours, and detected with anti-GST and anti-pThr480 antibodies. Note that pThr480 is absent in *pmk1*Δ cells, while its phosphorylation level was further enhanced in cells when both Pek1^DD^ and Pmk1-GBP-mCherry were present compared to cells only expressing either fusions. Blots shown are the representative of three independent experiments. pThr480 levels were normalized with ratio between anti-pThr480-recognized and GST bead-bound Slp1 fragment fusion in cells only expressing Slp1^(1-60aa)^-mEGFP-2xNLS-GST-Slp1^(456-488aa)^ set as 1.0. Error bars indicate mean ± standard deviation of three independent experiments. Two-tailed unpaired *t*-test was used to derive *p* values against wild-type cells.

**Figure 4-figure supplement 4.**
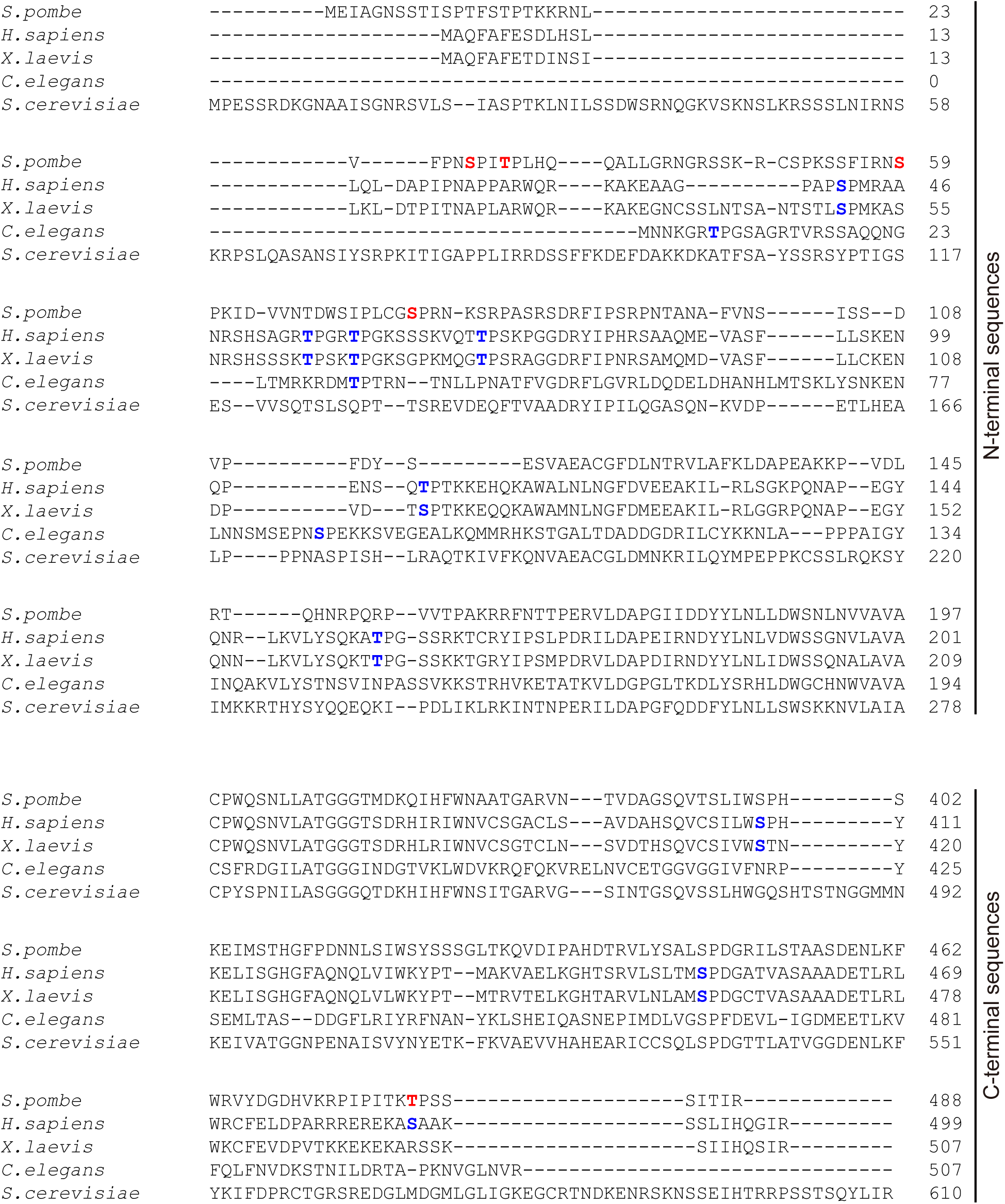
Sequence alignment performed with N-terminal and C-terminal tails of *S. pombe* Slp1 and its homologue sequences from human (*H. sapiens*), frog (*X. laevis*), worm (*C. elegans*) and budding yeast (*S. cerevisiae*). Five phosphorylation sites (Ser28, Thr31, Ser59, Ser76 and Thr480) in Slp1 identified in this study are indicated in red. Putative Cdk1 or MAPK phosphorylation sites in Slp1 homologues (e.g. Ser41, Thr55, Thr59, Thr70, Thr106, Thr157, Ser408, Ser452 and Ser487 in human Cdc20; Ser50, Thr64, Thr68 and Thr79 in frog Cdc20; and Thr7, Thr32 and Ser87 in worm Cdc20) suggested by previous studies are indicated in blue.

**Figure 4-figure supplement 5.**
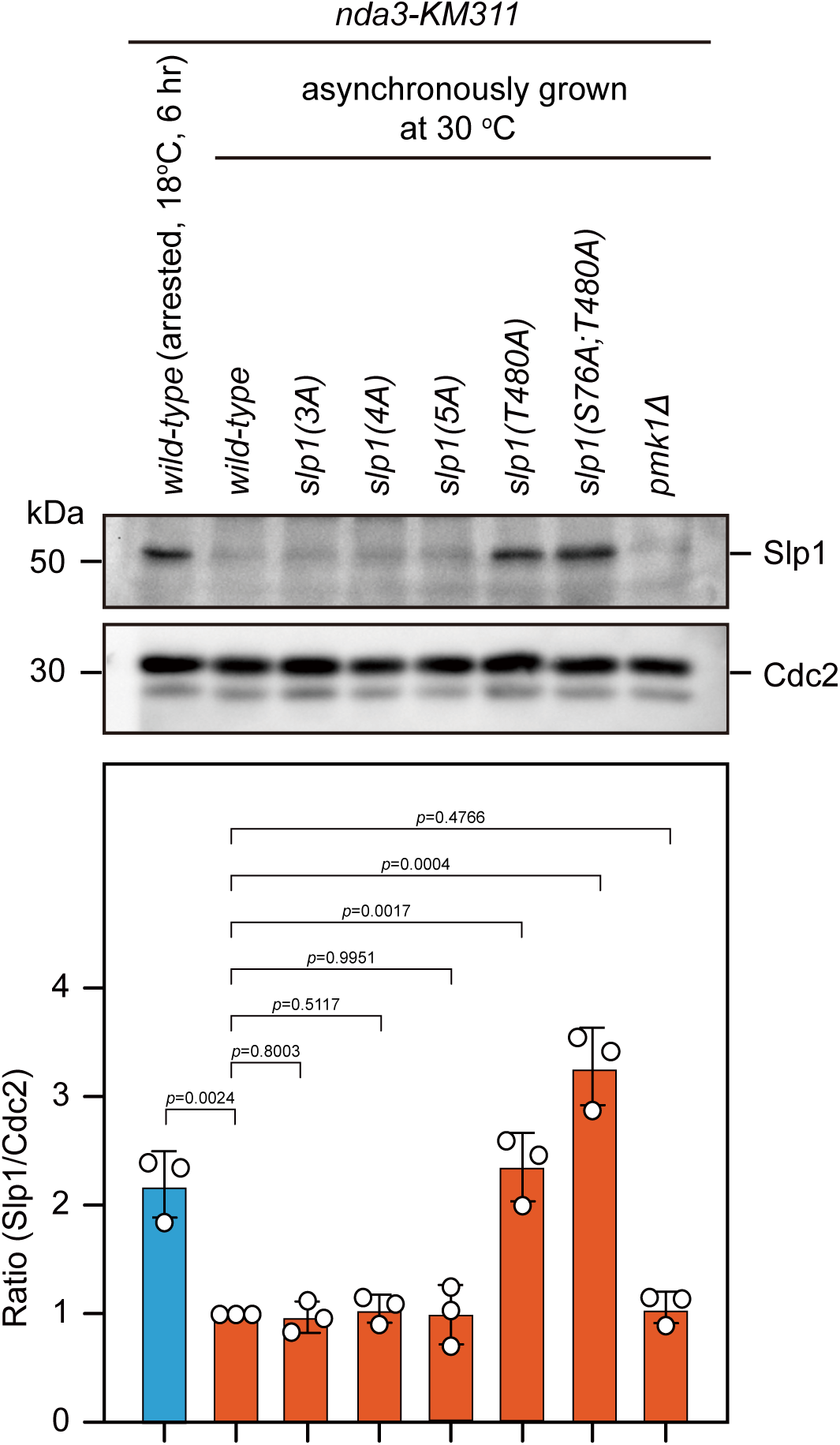
Mutant proteins of Slp1(T480A) and Slp1(S76A;T480A) are also stabilized in asynchronously growing cells. Immunoblotting of extracts from asynchronously growing *nda3-KM311* cells at 30 °C expressing wild type *slp1^+^* or *slp1* with the indicated mutations. An extract from *nda3-KM311 slp1^+^* cells synchronized with HU and arrested at metaphase by treatment at 18 °C for 6 hours was loaded as a control. Blots shown are the representative of three independent experiments. Slp1^Cdc20^ levels were quantified with the relative ratio between Slp1^Cdc20^ and Cdc2 in wild-type strain asynchronously grown 30 °C set as 1.0. Error bars indicate mean ± standard deviation of three independent experiments. Two-tailed unpaired *t*-test was used to derive *p* values against wild-type cells.

**Figure 4-figure supplement 6.**
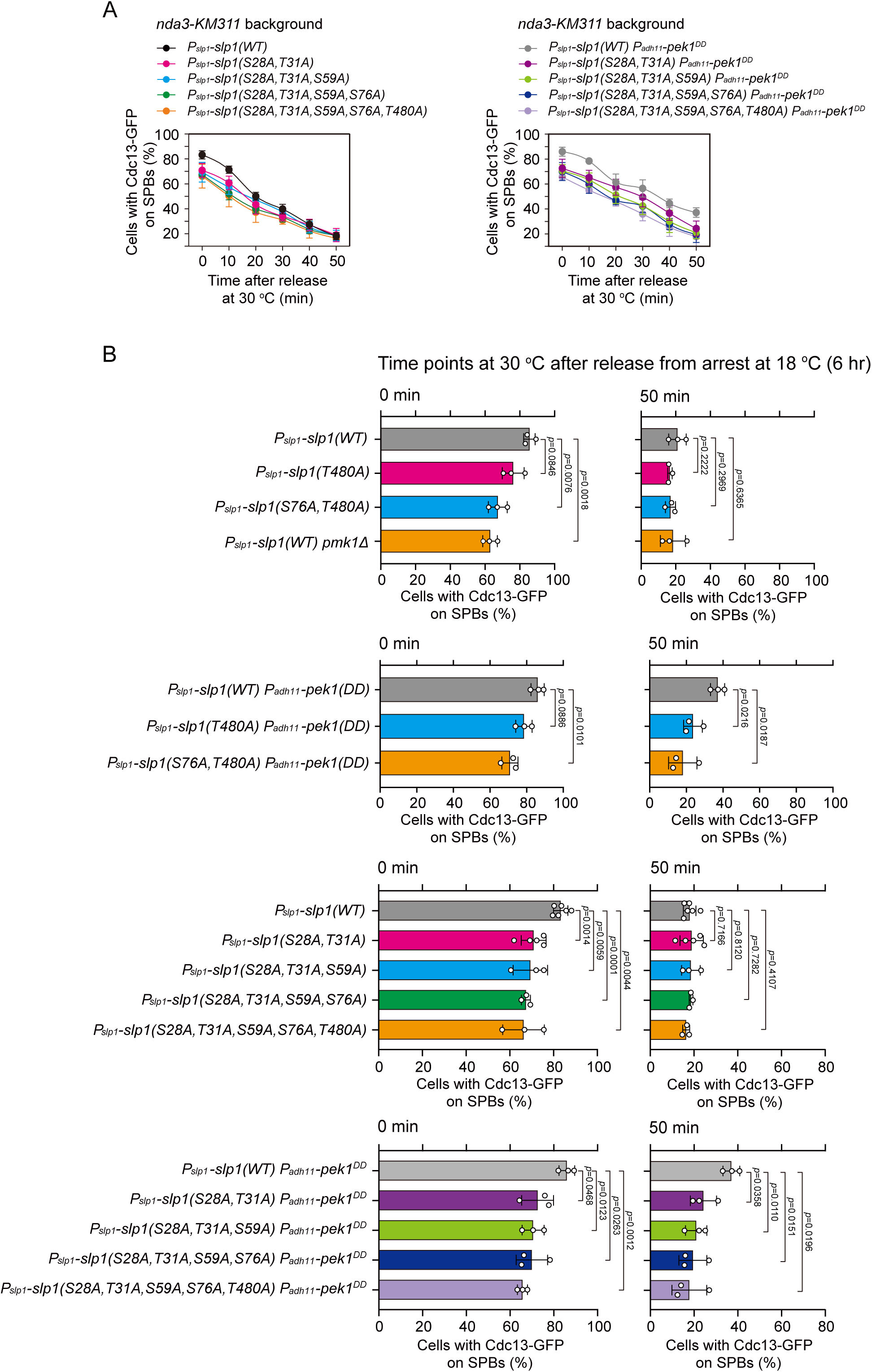
Time-course analyses and quantification of spindle checkpoint inactivation rate in phospho-deficient *slp1* mutants at 0 and 50 min after release at 30 °C from *nda3*-mediated arrest. Cells of indicated strains bearing Cdc13-GFP were grown at 30 °C to mid-log phase and arrested at 18 °C for 6 hours, and then released at 30 °C. For each time point, ≥300 cells were counted for every sample. **(A)** Time-course analyses of SAC activation and inactivation in *nda3-KM311 cdc13-GFP* strains with indicated genotypes. Error bars indicate mean ± standard deviation of three independent experiments. **(B)** Data from time points of 0 and 50 min after release in (A) and Figure 4H were subjected to statistical analysis. Error bars indicate mean ± standard deviation of three independent experiments. *p* values were calculated against wild-type cells.

**Figure 5-figure supplement 1.**
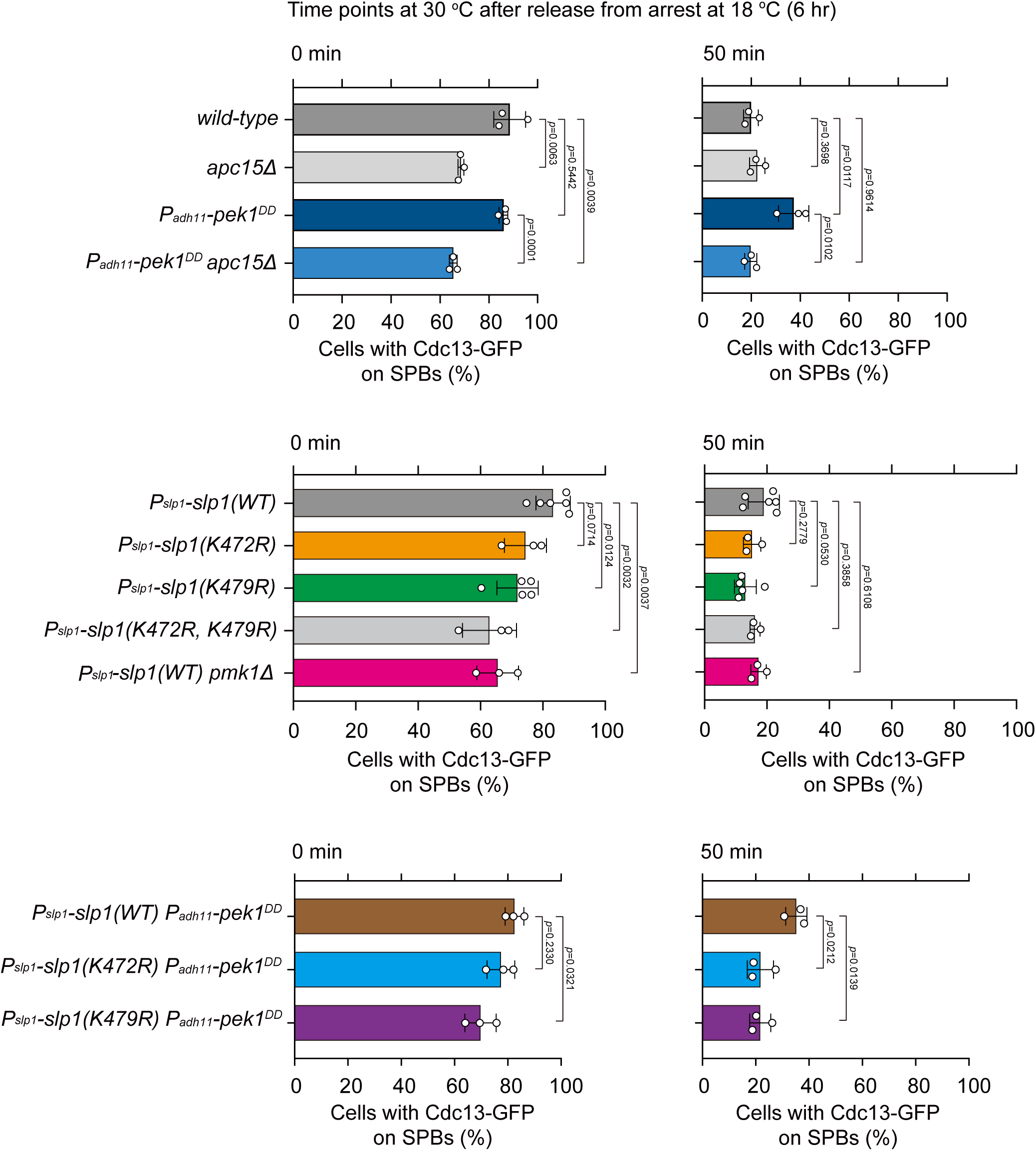
Quantification of spindle checkpoint inactivation rate in ubiquitylation-relevant *slp1* mutants at 0 and 50 min after release at 30 °C from *nda3*-mediated arrest. Cells of indicated strains bearing Cdc13-GFP were grown at 30 °C to mid-log phase and arrested at 18 °C for 6 hours, and then released at 30 °C. For each time point, ≥300 cells were counted for every sample. Data from time points of 0 and 50 min after release in Figure 5B and 5D were subjected to statistical analysis. Error bars indicate mean ± standard deviation of three independent experiments. *p* values were calculated against wild-type cells.

**Figure 5-figure supplement 2.**
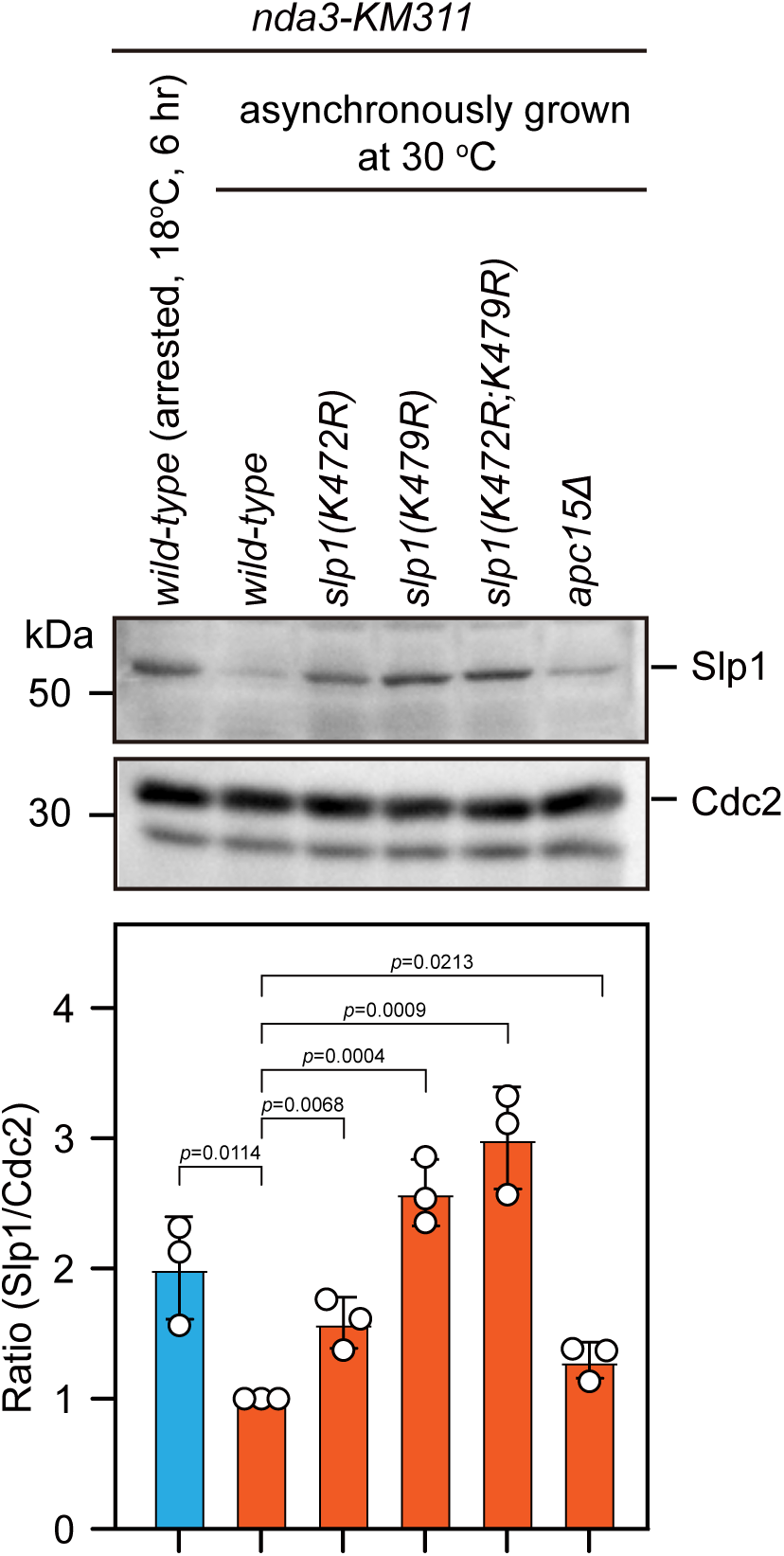
Slp1 proteins with mutations of K479R or K472R are also stabilized in asynchronously growing cells. Immunoblotting of extracts from asynchronously growing *nda3-KM311* cells at 30 °C expressing wild type *slp1^+^* or *slp1* with the indicated mutations. A strain with *apc15* deletion (*apc15Δ*) was included as a control for comparison, which has been previously shown to have stabilized Slp1 in interphase cells (Sewart and Hauf, 2017). An extract from *nda3-KM311 slp1^+^*cells synchronized with HU and arrested at metaphase by treatment at 18 °C for 6 hours was also loaded as a control. Blots shown are the representative of three independent experiments. Slp1^Cdc20^ levels were quantified with the relative ratio between Slp1^Cdc20^ and Cdc2 in wild-type strain asynchronously grown at 30 °C set as 1.0. Error bars indicate mean ± standard deviation of three independent experiments. Two-tailed unpaired *t*-test was used to derive *p* values against wild-type cells.

**Figure 5-figure supplement 3.**
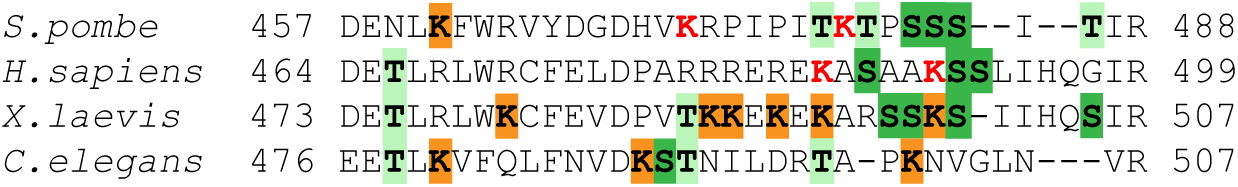
Sequence alignment of C-terminal tails of Cdc20 homologues. Sequence alignment performed with C-terminal tails of *S. pombe* Slp1 and its Cdc20 homologue sequences from human (*H. sapiens*), frog (*X. laevis*), and worm (*C. elegans*). Lysines residues in fission yeast Slp1 (K472 and K479) and human Cdc20 (K485 and K490), which have been confirmed in this study and previous studies (Danielsen et al., 2011; Mansfeld et al., 2011) respectively to be responsible for their ubiquitylation and degradation, are indicated in red. Other lysine residues and flanking threonine or serine residues in these sequences are also highlighted in orange or green respectively.

**Figure 5-figure supplement 4.**
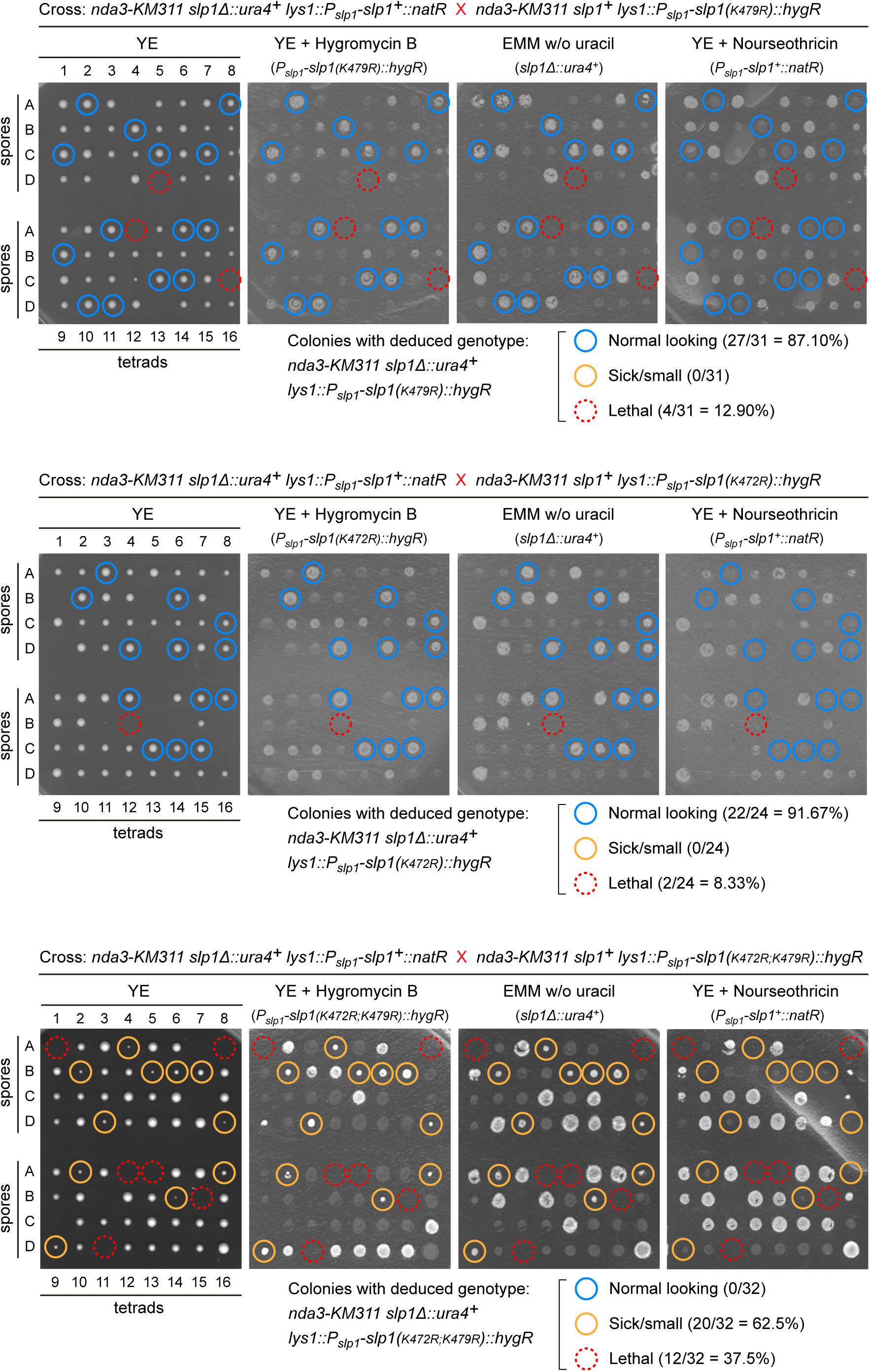
Viability of *slp1(K472R;K479R)* cells is compromised. Normal-looking four-spore asci obtained after crosses between *nda3-KM311* background strains carrying *slp1^+^ lys1::P_slp1_-slp1^(K472R)^::hyg^R^*, *slp1^+^ lys1::P_slp1_-slp1^(K479R)^::hyg^R^*or *slp1^+^ lys1::P_slp1_-slp1^(K472R;K479R)^::hyg^R^*and *slp1*Δ*::ura4^+^ lys1::P_slp1_-slp1^+^::nat^R^* strain were dissected using a micromanipulator. The genotypes of colonies formed from germinated spores were deduced after being replicated on selective plates. Mutant candidates of *nda3-KM311 slp1*Δ*::ura4^+^ lys1::P_slp1_-slp1^(K472R)/(K479R)/(K472R;K479R)^::hyg^R^*are indicated by light blue circles (normal looking colonies), yellow circles (small colonies) or dashed red circles (spores failing to germinate) on representative plates. Quantitative analyses of synthetic lethality of desired mutants based on dissected four-spore asci showed that simultaneous mutation of K472 and K479 in Slp1 to arginine (i.e. *K472R;K479R* double mutant) causes strong synthetic growth defects.

**Figure 5-figure supplement 5.**
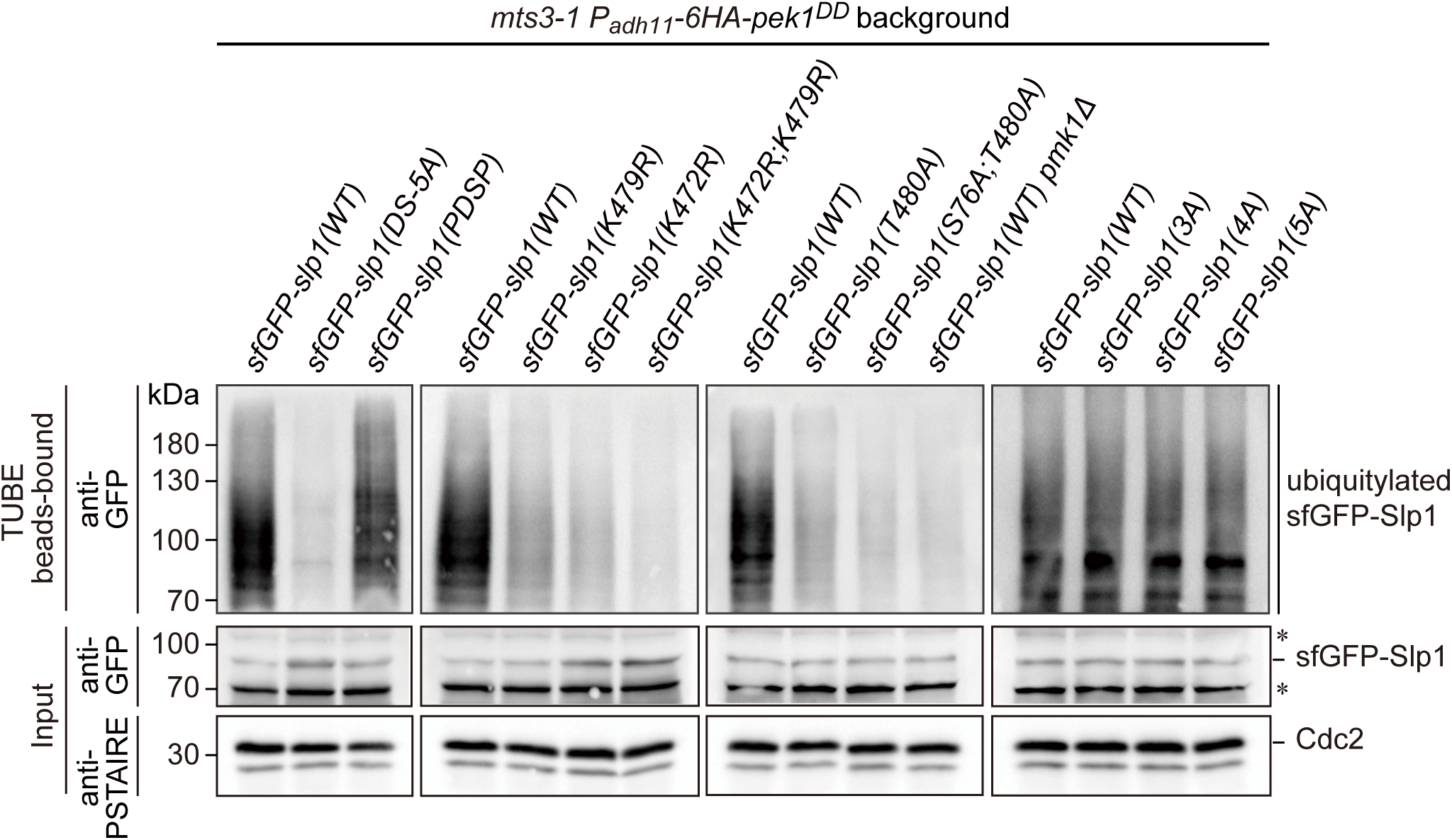
Enhanced Slp1^Cdc20^ ubiquitylation in *pek1^DD^* cells can be removed by Pmk1-docking-, Pmk1-phosphorylation- or ubiquitylation-deficient mutations in Slp1^Cdc20^. Cultures of *mts3-1 his5*::*P_adh11_*-*pek1^DD^::nat^R^* strains carrying sfGFP-tagged wild type or mutants of Slp1^Cdc20^ with mutations in Pmk1-docking motives (DS-5A, PDSP), ubiquitylation sites (K472R; K479R or K472R/K479R), Pmk1 phosphorylation sites (T480A; S76A/T480A), or combined Pmk1 and Cdk1 phosphorylation sites (3A; 4A or 5A) were first grown at 25 °C to mid-log phase and then shifted to 36 [for 3.5 hours to block cells in mitosis prior to harvesting and cell lysis. The strain *mts3-1 his5*::*P_adh11_*-*pek1^DD^::nat^R^ pmk1Δ* served as a negative control. Ubiquitinated proteins were pulled down from yeast lysates using Tandem Ubiquitin Binding Entities (TUBEs). The TUBE beads-bound samples were blotted with anti-GFP antibodies. Asterisks indicate unspecific bands recognized by anti-GFP antibodies.

**Figure 6-figure supplement 1.**
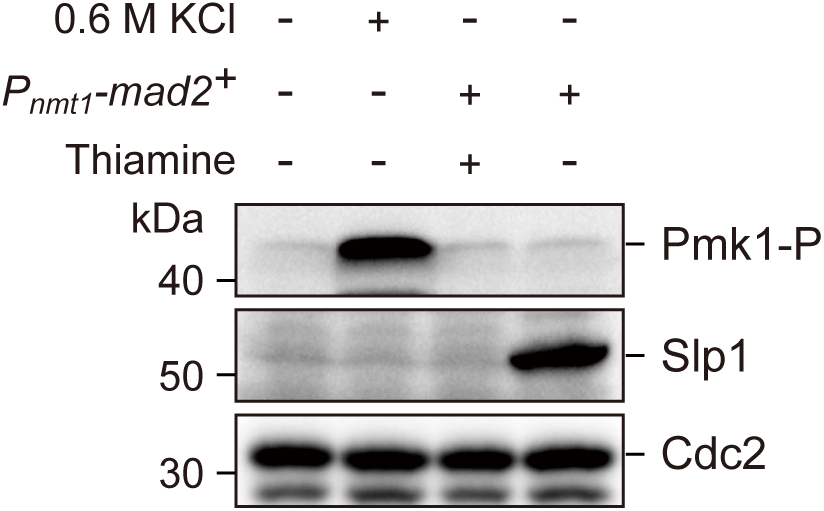
Activation of SAC by Mad2 overexpression does not trigger Pmk1 activation. Yeast strain carrying *P_nmt1_-mad2::leu1^+^* was pre-cultured in minimal medium with supplements (EMM5S) and 15 µM thiamine. Cells were then washed in EMM5S to remove thiamine before being grown in EMM5S at 30 °C for 18 hours to induce Mad2 overexpression. Wild-type cells grown in YE were treated with 0.6 M KCl for 60 min as a positive control for Pmk1 activation. Protein samples after SDS-PAGE were blotted with anti-phospho p42/44 antibodies as indicative of phosphorylated Pmk1 (Pmk1-P). Slp1^Cdc20^ levels were detected with anti-Slp1 antibodies and anti-PSTAIRE (detecting Cdc2) was used as loading control. Blots shown are the representative of three independent experiments.

**Figure 6-figure supplement 2.**
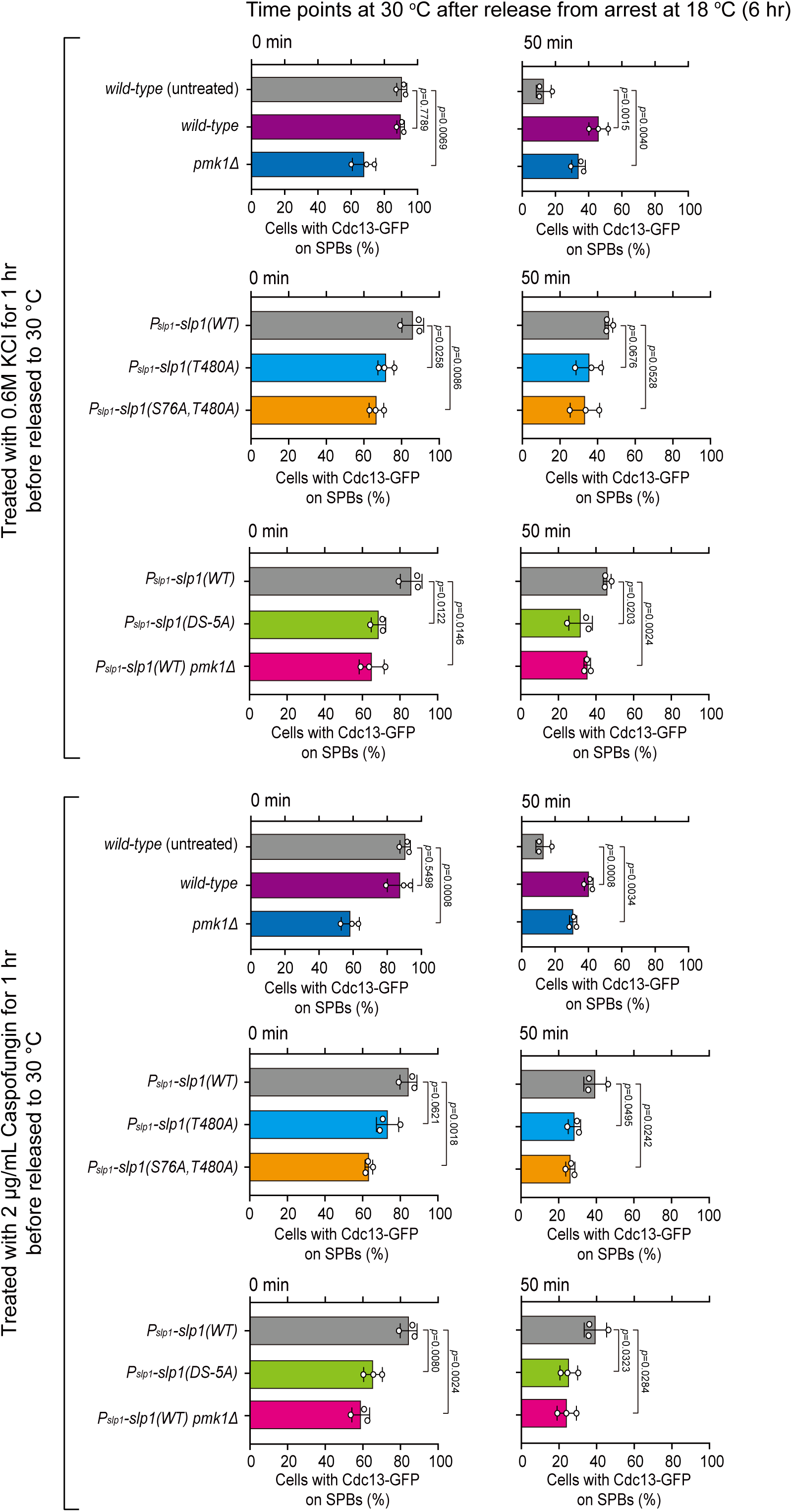
Quantification of spindle checkpoint inactivation rate in the presence of KCl or caspofungin treatment at 0 and 50 min after release at 30 °C from *nda3*-mediated arrest. Cells of indicated strains bearing Cdc13-GFP were grown at 30 °C to mid-log phase and arrested at 18 °C for 6 hours, and then released at 30 °C. For each time point, ≥300 cells were counted for every sample. Data from time points of 0 and 50 min after release in Figure 6D and 6E were subjected to statistical analysis. Error bars indicate mean ± standard deviation of three independent experiments. *p* values were calculated against wild-type cells.

**Figure 6-figure supplement 3.**
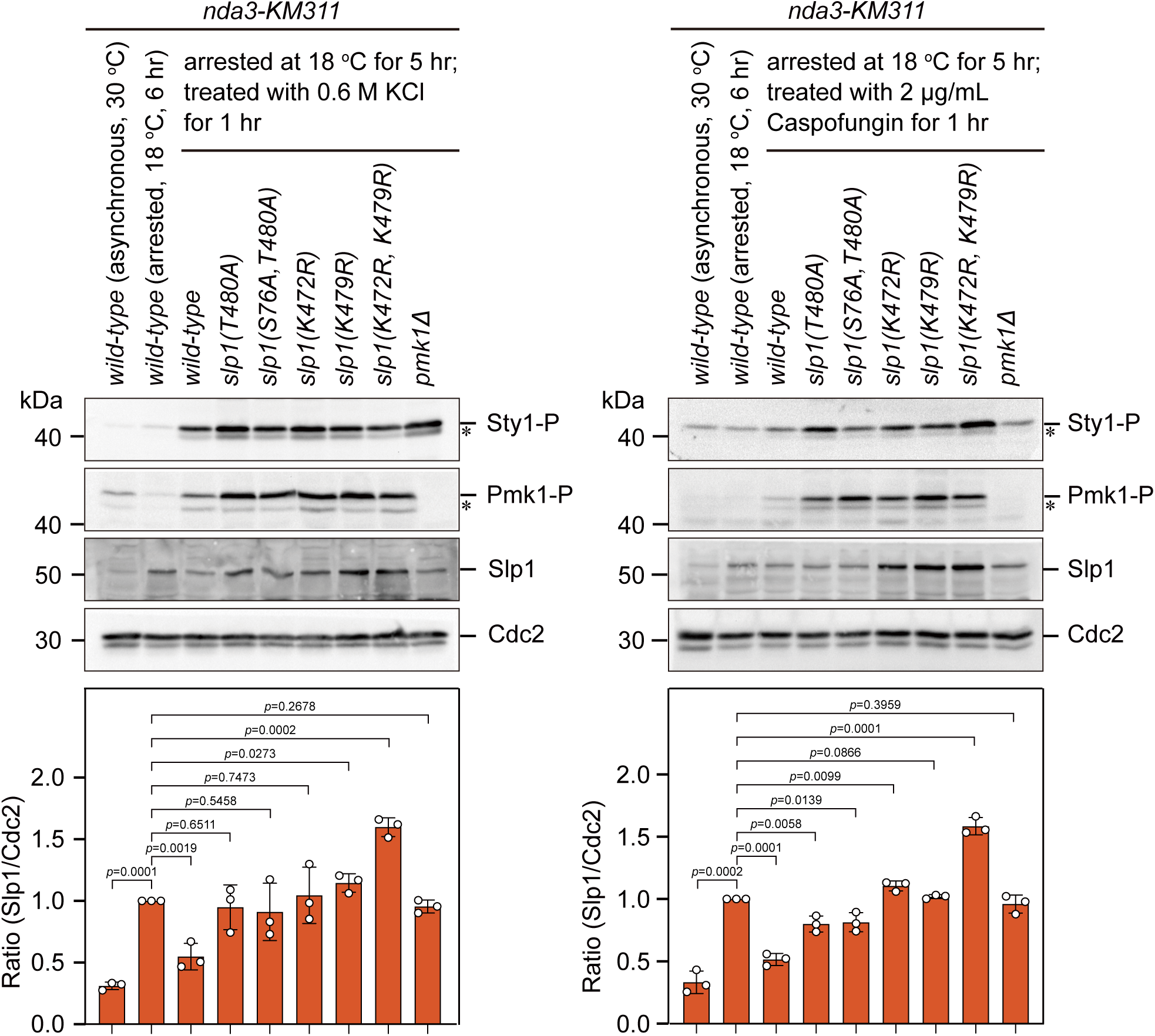
Immunoblot analysis of activation of MAPKs and Slp1^Cdc20^ protein levels in Pmk1 phosphorylation-deficient and ubiquitylation-deficient *slp1* mutants under stress. *nda3-KM311* strains expressing wild type *slp1^+^* or *slp1* with the indicated mutations were first synchronized with HU and then arrested at metaphase by treatment at 18 °C for 5 hours. Then, a final concentration of 0.6 M KCl or 2 μg/mL caspofungin was added into cultures and left at 18 °C for 1 hour before harvesting. *nda3-KM311 slp1^+^* cells either grown asynchronously at 30 °C or synchronized with HU and arrested at metaphase by treatment at 18 °C for 6 hours served as controls. Protein samples were subjected to SDS-PAGE and immunoblotted with anti-phospho p42/44 and anti-phospho p38 antibodies as indicative of phosphorylated Pmk1 (Pmk1-P) or phosphorylated Sty1 (Sty1-P), respectively. Slp1Cdc20 levels were detected with anti-Slp1 antibodies and anti-PSTAIRE (detecting Cdc2) was used as loading control. The experiment was repeated 3 times. Slp1^Cdc20^ levels were quantified with the relative ratio between Slp1^Cdc20^ and Cdc2 in wild-type strain synchronized and arrested at 18 °C for 6 hours set as 1.0. The mean value for each sample was calculated from three independent experiments, and *p* values were calculated against wild-type cells synchronized and arrested at 18 °C for 6 hours.

## Supplementary file legend

**Supplementary file 1.** Yeast strains used in this study.

## Source Data Legends

Figure 1-Source Data [raw data of time-course analyses of Cdc13-GFP at SPB for Figure 1D]

Figure 2-Source Data 1 [uncropped blots for Figure 2A-2C]

Figure 2-Source Data 2 [raw data of co-immunoprecipitation rate of Mad2/Mad3/Slp1, Slp1 level measurement, and RT-qPCR for Figure 2A-2D]

Figure 3-Source Data 1 [uncropped blots for Figure 3A, 3C and 3D]

Figure 3-Source Data 2 [raw data of Slp1 level measurement and time-course analyses of Cdc13-GFP at SPB for Figure 3D and 3E]

Figure 4-Source Data 1 [uncropped blots for Figure 4A and 4C-4G]

Figure 4-Source Data 2 [raw data of Slp1 level measurement and time-course analyses of Cdc13-GFP at SPB for Figure 4G and 4H]

Figure 5-Source Data 1 [uncropped blots for Figure 5A, 5C and 5F]

Figure 5-Source Data 2 [raw data of Slp1 level measurement and time-course analyses of Cdc13-GFP at SPB for Figure 5A-5D]

Figure 6-Source Data 1 [uncropped blots for Figure 6B and 6C]

Figure 6-Source Data 2 [raw data of time-course analyses of Cdc13-GFP at SPB for Figure 6D and 6E]

Figure 7-Source Data [raw data of defective chromosome segregation analyses for Figure 7C]

Figure 1-figure supplement 1-Source Data 1 [uncropped blots for Figure 1-figure supplement 1A]

Figure 1-figure supplement 1-Source Data 2 [raw data of time-course analyses of Cdc13-GFP at SPB for Figure 1-figure supplement 1B and 1C]

Figure 1-figure supplement 2-Source Data 1 [uncropped blots for Figure 1-figure supplement 2A]

Figure 1-figure supplement 2-Source Data 2 [raw data of time-course analyses of Cdc13-GFP at SPB for Figure 1-figure supplement 2C and 2D]

Figure 1-figure supplement 3-Source Data [raw data of quantitative analyses of cells with short spindles in response to Mad2 overexpression for Figure 1-figure supplement 3B]

Figure 2-figure supplement 1-Source Data 1 [uncropped blots for Figure 2-figure supplement 1C]

Figure 2-figure supplement 1-Source Data 2 [raw data of time-course analyses of arrest-and-release of *cdc25-22* mutants for Figure 2-figure supplement 1B]

Figure 3-figure supplement 2-Source Data [uncropped gels for Figure 3-figure supplement 2B]

Figure 3-figure supplement 3-Source Data [raw data of time-course analyses of Cdc13-GFP at SPB for Figure 3-figure supplement 3]

Figure 4-figure supplement 1-Source Data [uncropped gels for Figure 4-figure supplement 1A]

Figure 4-figure supplement 3-Source Data 1 [uncropped blots for Figure 4-figure supplement 3C]

Figure 4-figure supplement 3-Source Data 2 [raw data of quantitative analysis of Slp1 T480 phosphorylation levels for Figure 4-figure supplement 3C]

Figure 4-figure supplement 5-Source Data 1 [uncropped blots for Figure 4-figure supplement 5]

Figure 4-figure supplement 5-Source Data 2 [raw data of Slp1 level measurement for Figure 4-figure supplement 5]

Figure 4-figure supplement 6-Source Data [raw data of time-course analyses of Cdc13-GFP at SPB for Figure 4-figure supplement 6A and 6B]

Figure 5-figure supplement 1-Source Data [raw data of time-course analyses of Cdc13-GFP at SPB for Figure 5-figure supplement 1]

Figure 5-figure supplement 2-Source Data 1 [uncropped blots for Figure 5-figure supplement 2]

Figure 5-figure supplement 2-Source Data 2 [raw data of Slp1 level measurement for Figure 5-figure supplement 2]

Figure 5-figure supplement 5-Source Data [uncropped blots for Figure 5-figure supplement 5]

Figure 6-figure supplement 1-Source Data [uncropped blots for Figure 6-figure supplement 1]

Figure 6-figure supplement 2-Source Data [raw data of time-course analyses of Cdc13-GFP at SPB for Figure 6-figure supplement 2]

Figure 6-figure supplement 3-Source Data 1 [uncropped blots for Figure 6-figure supplement 3]

Figure 6-figure supplement 3-Source Data 2 [raw data of Slp1 level measurement for Figure 6-figure supplement 3]

## References

Akera, T., Goto, Y., Sato, M., Yamamoto, M., and Watanabe, Y. (2015). Mad1 promotes chromosome congression by anchoring a kinesin motor to the kinetochore. Nat Cell Biol 17, 1124–1133.

Alfieri, C., Chang, L., Zhang, Z., Yang, J., Maslen, S., Skehel, M., and Barford, D. (2016). Molecular basis of APC/C regulation by the spindle assembly checkpoint. Nature 536, 431–436.

Bai, S., Sun, L., Wang, X., Wang, S.M., Luo, Z.Q., Wang, Y.M., and Jin, Q.W. (2022). Recovery from spindle checkpoint-mediated arrest requires a novel Dnt1-dependent APC/C activation mechanism. Plos Genetics 18, e1010397.

Bardwell, A.J., Flatauer, L.J., Matsukuma, K., Thorner, J., and Bardwell, L. (2001). A conserved docking site in MEKs mediates high-affinity binding to MAP kinases and cooperates with a scaffold protein to enhance signal transmission. J Biol Chem 276, 10374–10386.

Bardwell, L., and Thorner, J. (1996). A conserved motif at the amino termini of MEKs might mediate high-affinity interaction with the cognate MAPKs. Trends in biochemical sciences 21, 373–374.

Cansado, J., Soto, T., Franco, A., Vicente-Soler, J., and Madrid, M. (2021). The Fission Yeast Cell Integrity Pathway: A Functional Hub for Cell Survival upon Stress and Beyond. J Fungi (Basel) 8, 1–32.

Chao, W.C., Kulkarni, K., Zhang, Z., Kong, E.H., and Barford, D. (2012). Structure of the mitotic checkpoint complex. Nature 484, 208–213.

Chen, R.H. (2004). Phosphorylation and activation of Bub1 on unattached chromosomes facilitate the spindle checkpoint. EMBO J 23, 3113–3121.

Chen, Y.H., Wang, G.Y., Hao, H.C., Chao, C.J., Wang, Y., and Jin, Q.W. (2017). Facile manipulation of protein localization in fission yeast through binding of GFP-binding protein to GFP. Journal of Cell Science 130, 1003–1015.

Chi, J.J., Li, H., Zhou, Z., Izquierdo-Ferrer, J., Xue, Y., Wavelet, C.M., Schiltz, G.E., Zhang, B., Cristofanilli, M., Lu, X., et al. (2019). A novel strategy to block mitotic progression for targeted therapy. EBioMedicine 49, 40–54.

Chung, E., and Chen, R.H. (2003). Phosphorylation of Cdc20 is required for its inhibition by the spindle checkpoint. Nat Cell Biol 5, 748–753.

D’Angiolella, V., Mari, C., Nocera, D., Rametti, L., and Grieco, D. (2003). The spindle checkpoint requires cyclin-dependent kinase activity. Genes Dev 17, 2520–2525.

Danielsen, J.M.R., Sylvestersen, K.B., Bekker-Jensen, S., Szklarczyk, D., Poulsen, J.W., Horn, H., Jensen, L.J., Mailand, N., and Nielsen, M.L. (2011). Mass Spectrometric Analysis of Lysine Ubiquitylation Reveals Promiscuity at Site Level. Molecular & Cellular Proteomics 10.

Edreira, T., Celador, R., Manjón, E., and Sánchez, Y. (2020). A novel checkpoint pathway controls actomyosin ring constriction trigger in fission yeast. eLife 9.

Fujimitsu, K., Grimaldi, M., and Yamano, H. (2016). Cyclin-dependent kinase 1-dependent activation of APC/C ubiquitin ligase. Science 352, 1121–1124.

Gomez-Gil, E., Martin-Garcia, R., Vicente-Soler, J., Franco, A., Vazquez-Marin, B., Prieto-Ruiz, F., Soto, T., Perez, P., Madrid, M., and Cansado, J. (2020). Stress-activated MAPK signaling controls fission yeast actomyosin ring integrity by modulating formin For3 levels. eLife 9, e57951.

Greil, C., Engelhardt, M., and Wasch, R. (2022). The Role of the APC/C and Its Coactivators Cdh1 and Cdc20 in Cancer Development and Therapy. Front Genet 13, 941565.

Hartmuth, S., and Petersen, J. (2009). Fission yeast Tor1 functions as part of TORC1 to control mitotic entry through the stress MAPK pathway following nutrient stress. J Cell Sci 122, 1737–1746.

Heinrich, S., Geissen, E.M., Kamenz, J., Trautmann, S., Widmer, C., Drewe, P., Knop, M., Radde, N., Hasenauer, J., and Hauf, S. (2013). Determinants of robustness in spindle assembly checkpoint signalling. Nat Cell Biol 15, 1328–1339.

Hiraoka, Y., Toda, T., and Yanagida, M. (1984). The NDA3 gene of fission yeast encodes beta-tubulin: a cold-sensitive nda3 mutation reversibly blocks spindle formation and chromosome movement in mitosis. Cell 39, 349–358.

Hjerpe, R., Aillet, F., Lopitz-Otsoa, F., Lang, V., England, P., and Rodriguez, M.S. (2009). Efficient protection and isolation of ubiquitylated proteins using tandem ubiquitin-binding entities. EMBO Rep 10, 1250–1258.

Izawa, D., and Pines, J. (2015). The mitotic checkpoint complex binds a second CDC20 to inhibit active APC/C. Nature 517, 631–634.

Jeong, S.M., Bui, Q.T., Kwak, M., Lee, J.Y., and Lee, P.C. (2022). Targeting Cdc20 for cancer therapy. Biochim Biophys Acta Rev Cancer 1877, 188824.

Jia, L., Li, B., and Yu, H. (2016). The Bub1-Plk1 kinase complex promotes spindle checkpoint signalling through Cdc20 phosphorylation. Nature communications 7, 10818.

Kapanidou, M., Curtis, N.L., and Bolanos-Garcia, V.M. (2017). Cdc20: At the Crossroads between Chromosome Segregation and Mitotic Exit. Trends in biochemical sciences 42, 193–205.

Kim, D.U., Hayles, J., Kim, D., Wood, V., Park, H.O., Won, M., Yoo, H.S., Duhig, T., Nam, M., Palmer, G., et al. (2010). Analysis of a genome-wide set of gene deletions in the fission yeast Schizosaccharomyces pombe. Nat Biotechnol 28, 617–623.

Kim, S.H., Lin, D.P., Matsumoto, S., Kitazono, A., and Matsumoto, T. (1998). Fission yeast Slp1: an effector of the Mad2-dependent spindle checkpoint. Science 279, 1045–1047.

Kim, T., Lara-Gonzalez, P., Prevo, B., Meitinger, F., Cheerambathur, D.K., Oegema, K., and Desai, A. (2017). Kinetochores accelerate or delay APC/C activation by directing Cdc20 to opposing fates. Genes Dev 31, 1089–1094.

Klomp, J.A., Klomp, J.E., Stalnecker, C.A., Bryant, K.L., Edwards, A.C., Drizyte-Miller, K., Hibshman, P.S., Diehl, J.N., Lee, Y.S., Morales, A.J., et al. (2024). Defining the KRAS- and ERK-dependent transcriptome in KRAS-mutant cancers. Science 384, eadk0775.

Kraft, C., Herzog, F., Gieffers, C., Mechtler, K., Hagting, A., Pines, J., and Peters, J.M. (2003). Mitotic regulation of the human anaphase-promoting complex by phosphorylation. EMBO J 22, 6598–6609.

Kramer, E.R., Scheuringer, N., Podtelejnikov, A.V., Mann, M., and Peters, J.M. (2000). Mitotic regulation of the APC activator proteins CDC20 and CDH1. Mol Biol Cell 11, 1555–1569.

Labit, H., Fujimitsu, K., Bayin, N.S., Takaki, T., Gannon, J., and Yamano, H. (2012). Dephosphorylation of Cdc20 is required for its C-box-dependent activation of the APC/C. EMBO J 31, 3351–3362.

Lopez-Aviles, S., Grande, M., Gonzalez, M., Helgesen, A.L., Alemany, V., Sanchez-Piris, M., Bachs, O., Millar, J.B., and Aligue, R. (2005). Inactivation of the Cdc25 phosphatase by the stress-activated Srk1 kinase in fission yeast. Mol Cell 17, 49–59.

Lopez-Aviles, S., Lambea, E., Moldon, A., Grande, M., Fajardo, A., Rodriguez-Gabriel, M.A., Hidalgo, E., and Aligue, R. (2008). Activation of Srk1 by the mitogen-activated protein kinase Sty1/Spc1 precedes its dissociation from the kinase and signals its degradation. Mol Biol Cell 19, 1670–1679.

Madrid, M., Soto, T., Khong, H.K., Franco, A., Vicente, J., Pérez, P., Gacto, M., and Cansado, J. (2006). Stress-induced response, localization, and regulation of the Pmk1 cell integrity pathway in. Journal of Biological Chemistry 281, 2033–2043.

Madrid, M., Vázquez-Marín, B., Franco, A., Soto, T., Vicente-Soler, J., Gacto, M., and Cansado, J. (2016). Multiple crosstalk between TOR and the cell integrity MAPK signaling pathway in fission yeast. Sci Rep-Uk 6.

Mansfeld, J., Collin, P., Collins, M.O., Choudhary, J.S., and Pines, J. (2011). APC15 drives the turnover of MCC-CDC20 to make the spindle assembly checkpoint responsive to kinetochore attachment. Nat Cell Biol 13, 1234–1243.

May, K.M., Paldi, F., and Hardwick, K.G. (2017). Fission Yeast Apc15 Stabilizes MCC-Cdc20-APC/C Complexes, Ensuring Efficient Cdc20 Ubiquitination and Checkpoint Arrest. Curr Biol 27, 1221–1228.

McAinsh, A.D., and Kops, G.J.P.L. (2023). Principles and dynamics of spindle assembly checkpoint signalling. Nat Rev Mol Cell Bio 24, 543–559.

Minshull, J., Sun, H., Tonks, N.K., and Murray, A.W. (1994). A MAP kinase-dependent spindle assembly checkpoint in Xenopus egg extracts. Cell 79, 475–486.

Murray, A.W. (2011). A brief history of error. Nat Cell Biol 13, 1178–1182.

Musacchio, A. (2015). The Molecular Biology of Spindle Assembly Checkpoint Signaling Dynamics. Curr Biol 25, R1002–1018.

Ozoe, F., Kurokawa, R., Kobayashi, Y., Jeong, H.T., Tanaka, K., Sen, K., Nakagawa, T., Matsuda, H., and Kawamukai, M. (2002). The 14–3–3 proteins Rad24 and Rad25 negatively regulate Byr2 by affecting its localization in Schizosaccharomyces pombe. . Molecular and Cellular Biology 22, 15.

Pan, J., and Chen, R.H. (2004). Spindle checkpoint regulates Cdc20p stability in Saccharomyces cerevisiae. Genes Dev 18, 1439–1451.

Perez, P., and Cansado, J. (2010). Cell integrity signaling and response to stress in fission yeast. Curr Protein Pept Sci 11, 680–692.

Peters, J.M. (2006). The anaphase promoting complex/cyclosome: a machine designed to destroy. Nat Rev Mol Cell Biol 7, 644–656.

Petersen, J., and Hagan, I.M. (2005). Polo kinase links the stress pathway to cell cycle control and tip growth in fission yeast. Nature 435, 507–512.

Petersen, J., and Nurse, P. (2007). TOR signalling regulates mitotic commitment through the stress MAP kinase pathway and the Polo and Cdc2 kinases. Nat Cell Biol 9, 1263–1272.

Pinna, L.A., and Ruzzene, M. (1996). How do protein kinases recognize their substrates? Bba-Mol Cell Res 1314, 191–225.

Plotnikov, A., Zehorai, E., Procaccia, S., and Seger, R. (2011). The MAPK cascades: signaling components, nuclear roles and mechanisms of nuclear translocation. Biochim Biophys Acta 1813, 1619–1633.

Primorac, I., and Musacchio, A. (2013). Panta rhei: the APC/C at steady state. J Cell Biol 201, 177–189.

Qiao, R., Weissmann, F., Yamaguchi, M., Brown, N.G., VanderLinden, R., Imre, R., Jarvis, M.A., Brunner, M.R., Davidson, I.F., Litos, G., et al. (2016). Mechanism of APC/CCDC20 activation by mitotic phosphorylation. Proc Natl Acad Sci U S A 113, E2570–2578.

Repetto, M.V., Winters, M.J., Bush, A., Reiter, W., Hollenstein, D.M., Ammerer, G., Pryciak, P.M., and Colman-Lerner, A. (2018). CDK and MAPK Synergistically Regulate Signaling Dynamics via a Shared Multi-site Phosphorylation Region on the Scaffold Protein Ste5. Molecular Cell 69, 938-+.

Ronkina, N., and Gaestel, M. (2022). MAPK-Activated Protein Kinases: Servant or Partner? Annu Rev Biochem, 12.11–12.36.

Rothbauer, U., Zolghadr, K., Tillib, S., Nowak, D., Schermelleh, L., Gahl, A., Backmann, N., Conrath, K., Muyldermans, S., Cardoso, M.C., et al. (2006). Targeting and tracing antigens in live cells with fluorescent nanobodies. Nat Methods 3, 887–889.

Sacristan-Reviriego, A., Madrid, M., Cansado, J., Martin, H., and Molina, M. (2014). A conserved non-canonical docking mechanism regulates the binding of dual specificity phosphatases to cell integrity mitogen-activated protein kinases (MAPKs) in budding and fission yeasts. PLoS One 9, e85390.

Saitoh, S., Takahashi, K., and Yanagida, M. (1997). Mis6, a fission yeast inner centromere protein, acts during G1/S and forms specialized chromatin required for equal segregation. Cell 90, 131–143.

Sczaniecka, M., Feoktistova, A., May, K.M., Chen, J.S., Blyth, J., Gould, K.L., and Hardwick, K.G. (2008). The spindle checkpoint functions of Mad3 and Mad2 depend on a Mad3 KEN box-mediated interaction with Cdc20-anaphase-promoting complex (APC/C). J Biol Chem 283, 23039–23047.

Sewart, K., and Hauf, S. (2017). Different Functionality of Cdc20 Binding Sites within the Mitotic Checkpoint Complex. Curr Biol 27, 1213–1220.

Shiozaki, K., and Russell, P. (1995a). Cell-cycle control linked to extracellular environment by MAP kinase pathway in fission yeast. Nature 378, 739–743.

Shiozaki, K., and Russell, P. (1995b). Counteractive roles of protein phosphatase 2C (PP2C) and a MAP kinase kinase homolog in the osmoregulation of fission yeast. Embo J 14, 492–502.

Shiozaki, K., Shiozaki, M., and Russell, P. (1998). Heat stress activates fission yeast Spc1/StyI MAPK by a MEKK-independent mechanism. Mol Biol Cell 9, 1339–1349.

Smith, D.A., Toone, W.M., Chen, D., Bahler, J., Jones, N., Morgan, B.A., and Quinn, J. (2002). The Srk1 protein kinase is a target for the Sty1 stress-activated MAPK in fission yeast. J Biol Chem 277, 33411–33421.

Steen, J.A., Steen, H., Georgi, A., Parker, K., Springer, M., Kirchner, M., Hamprecht, F., and Kirschner, M.W. (2008). Different phosphorylation states of the anaphase promoting complex in response to antimitotic drugs: a quantitative proteomic analysis. Proc Natl Acad Sci U S A 105, 6069–6074.

Sudakin, V., Chan, G.K., and Yen, T.J. (2001). Checkpoint inhibition of the APC/C in HeLa cells is mediated by a complex of BUBR1, BUB3, CDC20, and MAD2. J Cell Biol 154, 925–936.

Sugiura, R., Toda, T., Dhut, S., Shuntoh, H., and Kuno, T. (1999). The MAPK kinase Pek1 acts as a phosphorylation-dependent molecular switch. Nature 399, 479–483.

Sullivan, M., and Morgan, D.O. (2007). Finishing mitosis, one step at a time. Nat Rev Mol Cell Biol 8, 894–903.

Swaffer, M.P., Jones, A.W., Flynn, H.R., Snijders, A.P., and Nurse, P. (2018). Quantitative Phosphoproteomics Reveals the Signaling Dynamics of Cell-Cycle Kinases in the Fission Yeast. Cell Reports 24, 503–514.

Takenaka, K., Gotoh, Y., and Nishida, E. (1997). MAP kinase is required for the spindle assembly checkpoint but is dispensable for the normal M phase entry and exit in Xenopus egg cell cycle extracts. J Cell Biol 136, 1091–1097.

Tang, Z., Shu, H., Oncel, D., Chen, S., and Yu, H. (2004). Phosphorylation of Cdc20 by Bub1 provides a catalytic mechanism for APC/C inhibition by the spindle checkpoint. Mol Cell 16, 387–397.

Uzunova, K., Dye, B.T., Schutz, H., Ladurner, R., Petzold, G., Toyoda, Y., Jarvis, M.A., Brown, N.G., Poser, I., Novatchkova, M., et al. (2012). APC15 mediates CDC20 autoubiquitylation by APC/C(MCC) and disassembly of the mitotic checkpoint complex. Nat Struct Mol Biol 19, 1116–1123.

Vanoosthuyse, V., and Hardwick, K.G. (2009). A novel protein phosphatase 1-dependent spindle checkpoint silencing mechanism. Curr Biol 19, 1176–1181.

Wang, Y., Li, W.Z., Johnson, A.E., Luo, Z.Q., Sun, X.L., Feoktistova, A., McDonald, W.H., McLeod, I., Yates, J.R., 3rd, Gould, K.L., et al. (2012). Dnt1 acts as a mitotic inhibitor of the spindle checkpoint protein dma1 in fission yeast. Mol Biol Cell 23, 3348–3356.

Yamada, H.Y., Matsumoto, S., and Matsumoto, T. (2000). High dosage expression of a zinc finger protein, Grt1, suppresses a mutant of fission yeast slp1(+), a homolog of CDC20/p55CDC/Fizzy. J Cell Sci 113 (*Pt 22*), 3989–3999.

Yamaguchi, M., VanderLinden, R., Weissmann, F., Qiao, R., Dube, P., Brown, N.G., Haselbach, D., Zhang, W., Sidhu, S.S., Peters, J.M., et al. (2016). Cryo-EM of Mitotic Checkpoint Complex-Bound APC/C Reveals Reciprocal and Conformational Regulation of Ubiquitin Ligation. Mol Cell 63, 593–607.

Yamano, H. (2019). APC/C: current understanding and future perspectives. F1000Res 8, 725.

Yen, A.H., and Yang, J.L. (2010). Cdc20 Proteolysis Requires p38 MAPK Signaling and Cdh1-Independent APC/C Ubiquitination During Spindle Assembly Checkpoint Activation by Cadmium. Journal of Cellular Physiology 223, 327–334.

Yoon, H.J., Feoktistova, A., Wolfe, B.A., Jennings, J.L., Link, A.J., and Gould, K.L. (2002). Proteomics analysis identifies new components of the fission and budding yeast anaphase-promoting complexes. Curr Biol 12, 2048–2054.

Yu, H. (2007). Cdc20: a WD40 activator for a cell cycle degradation machine. Mol Cell 27, 3–16.

Yudkovsky, Y., Shteinberg, M., Listovsky, T., Brandeis, M., and Hershko, A. (2000). Phosphorylation of Cdc20/fizzy negatively regulates the mammalian cyclosome/APC in the mitotic checkpoint. Biochem Biophys Res Commun 271, 299–304.

Zhang, S., Chang, L., Alfieri, C., Zhang, Z., Yang, J., Maslen, S., Skehel, M., and Barford, D. (2016). Molecular mechanism of APC/C activation by mitotic phosphorylation. Nature 533, 260–264.

Zhao, Y., and Chen, R.H. (2006). Mps1 phosphorylation by MAP kinase is required for kinetochore localization of spindle-checkpoint proteins. Curr Biol 16, 1764–1769.

Zuin, A., Carmona, M., Morales-Ivorra, I., Gabrielli, N., Vivancos, A.P., Ayte, J., and Hidalgo, E. (2010). Lifespan extension by calorie restriction relies on the Sty1 MAP kinase stress pathway. EMBO J 29, 981–991.

