## supplemental Table 1 for "Negative regulation of APC/C activation by MAPK-mediated attenuation of Cdc20^Slp1^ under stress"

Supplementary file 1. Yeast strains used in this study.

| Strain | Genotype |
| --- | --- |
| JY3 | *h^-^ ade6-216 leu1-32 ura4-D18* |
| JY4 | *h^+^ ade6-216 leu1-32 ura4-D18* |
| JY21 | *h^-^ cdc25-22 ura4-D18* |
| JY77 | *h^-^ mad2Δ::ura4^+^ ura4^-^ leu1^-^* |
| JY96 | *h^+^ mts3-1 ura4-D18 leu1-32* |
| JY227 | *h^-^ nda3-KM311 leu1-32 ura4-D18 ade6-21X* |
| JY2162 | *h^-^ sty1Δ::ura4^+^ ura4-D18 leu1-32* |
| JY3058 | *h^-^ nda3-KM311 cdc13-117::cdc13-GFP:LEU2 ark1-as3**::hygR ura4^-^* |
| JY4022 | *h^?^ nda3-KM311 lid1-TAP::kanR mad2-GFP::kanR mad3-GFP::his3^+^ leu1-32 ura4-D18 his3-D1 ade6-216* |
| JY4773 | *h^-^ pmk1Δ ::ura4^+^ ura4-D18 leu1-32* |
| JY4782 | *h^-^ spk1Δ::ura4^+^ ura4-D18 leu1-32* |
| JY5407 | *h^+^ nda3-KM311 cdc13-117::cdc13-GFP::LEU2 ura4^-^* |
| JY8607 | *h^+^ nda3-KM311 cdc13-117::cdc13-GFP::LEU2 pDUAL-P_slp1(long2)_-slp1-T_slp1_::leu1^+^* |
| JY8731 | *h^-^ nda3-KM311 cdc13-117::cdc13-GFP::LEU2 lys1Δ::-P_slp1(1504)_-slp1::hygR* |
| JY9122 | *h^?^ nda3-KM311 apc15Δ::KanR lid1-TAP-kanR mad2-GFP::kanR mad3-GFP-his3^+^* |
| JY9150 | *h^+^ nda3-KM311 cdc13-117::cdc13-GFP::LEU2 pmk1Δ::ura4 ura4? leu1**?* |
| JY9177 | *h^+^ nda3-KM311 mad3-GFP-his3^+^ mad2-13myc::hygR* |
| JY9182 | *h^?^ nda3-KM311 apc15Δ::kanR* |
| JY9212 | *h^+^ nda3-KM311 sty1-3HA::kanR* |
| JY9241 | *h^-^ nda3-KM311 pmk1-HA-6His::ura4^+^ ura4-D18 leu1-32* |
| JY9245 | *h^?^ nda3-KM311 pmk1Δ::ura4^+^* |
| JY9371 | *h^?^ nda3-KM311 lid1-TAP-kanR mad2-GFP::kanR mad3-GFP-his3^+^ pmk1Δ::ura4^+^* |
| JY9400 | *h^+^ nda3-KM311 cdc13-117::cdc13-GFP::LEU2 sty1-T97A leu1? ade6^-^ ura4^-^* |
| JY9401 | *h^-^ nda3-KM311 sty1-T97A* |
| JY9585 | *h^+^ lys1Δ::P_adh11_-6×HA-pek1(DD)[S234D, T238D]::hygR ura4-D18 leu1-32* |
| JY9586 | *h^+^ lys1Δ::P_adh21_-6×HA-pek1(DD)[S234D, T238D]::hygR ura4-D18 leu1-32* |
| JY9589 | *h^+^ nda3-KM311 cdc13-117::cdc13-GFP::LEU2*  *lys1Δ::P_adh11_-6×HA-pek1(DD)[S234D,T238D]::hygR* |
| JY9590 | *h^+^ nda3-KM311 cdc13-117::cdc13-GFP::LEU2*  *lys1Δ::P_adh21_-6×HA-pek1(DD)[S234D,T238D]::hygR* |
| JY9655 | *h^?^ nda3-KM311 pmk1Δ::ura4^+^ sty1-T97A* |
| JY9668 | *h^?^ nda3-KM311 lys1Δ::P_adh21_-6×HA-pek1(DD)[S234D, T238D]::hygR* |
| JY9672 | *h^?^ nda3-KM311 lys1Δ::P_adh11_-6×HA-pek1(DD)[S234D, T238D]::hygR* |
| JY9685 | *h^+^ nda3-KM311 mad3-GFP-his3^+^ mad2-13myc::hygR wis1-DD::ura4^+^* |
| JY9686 | *h^-^ nda3-KM311 lid1-TAP-kanR mad2-GFP::kanR mad3-GFP-his3^+^ apc15Δ::kanR lys1Δ::P_adh11_-6×HA-wis1(DD)[S469D;T473D]::hygR* |
| JY9720 | *h^+^ nda3-KM311 mad3-GFP-his3^+^ mad2-13myc::hygR sty1-T97A* |
| JY9777 | *h^+^ nda3-KM311 slp1Δ::ura4^+^ lys1Δ::P_slp1(1504)_-slp1-hygR ura4-D18 leu1-32* |
| JY9790 | *h^-^ nda3-KM311 lid1-TAP-kanR mad2-GFP::kanR mad3-GFP-his3^+^ sty1-T97A* |
| JY9837 | *h^?^ nda3-KM311 slp1Δ::ura4^+^ pmk1Δ::ura4^+^*  *lys1::P_slp1(1504)_-slp1-hygR ura4-D18 leu1* |
| JY9840 | *h^+^ nda3-KM311 lid1-TAP-kanR mad2-GFP::kanR mad3-GFP-his3^+^ lys1Δ::P_adh21_-6×HA-pek1(DD)[S234D, T238D]::hygR* |
| JY9841 | *h^?^ nda3-KM311 cdc13-GFP::kanR cdc2-as1(F84G)* |
| JY9860 | *h^+^ nda3-KM311 lid1-TAP-kanR mad2-GFP::kanR mad3-GFP-his3^+^ lys1Δ::P_adh11_-6×HA-pek1(DD)[S234D, T238D]::hygR* |
| JY9890 | *h^-^ nda3-KM311 slp1Δ::ura4^+^ cdc13-117::cdc13-GFP::LEU2 lys1Δ::P_slp1(1504_)-slp1::hygR ura4^-^* |
| JY9892 | *h^+^ nda3-KM311 cdc13-117::cdc13-GFP::LEU2 slp1Δ::ura4^+^ lys1Δ::P_slp1(1504)_-slp1::hygR ura4^-^* |
| JY9898 | *h^+^ nda3-KM311 slp1Δ::ura4^+^ pmk1Δ::ura4^+^ sty1-T97A lys1Δ::P_slp1(1504)_-slp1-hygR cdc13-117::cdc13-GFP::LEU2 leu1^-^ ura4^-^* |
| JY9914 | *h^-^ nda3-KM311 lys1Δ::P_adh21_-6×HA-wis1(DD)[S469D;T473D]::hygR* |
| JY9915 | *h^-^ nda3-KM311 lys1Δ::P_adh11_-6×HA-wis1(DD)[S469D;T473D]::hygR* |
| JY9916 | *h^-^ nda3-KM311 lid1-TAP-kanR mad2-GFP::kanR mad3-GFP-his3^+^ lys1Δ::P_adh21_-6×HA-wis1(DD)[S469D;T473D]::hygR* |
| JY9928 | *h^+^ nda3-KM311 cdc13-117::cdc13-GFP::LEU2 lys1Δ::P_adh21_-6×HA-wis1(DD)[S469D,T473D]::hygR ura4^-^* |
| JY9929 | *h^+^ nda3-KM311 cdc13-117::cdc13-GFP::LEU2 lys1Δ::P_adh11_-6×HA-wis1(DD)[S469D,T473D]::hygR ura4^-^* |
| JY9934 | *h^-^ nda3-KM311 lid1-TAP-kanR mad2-GFP::kanR mad3-GFP-his3^+^ lys1Δ::P_adh11_-6×HA-wis1(DD)[S469D;T473D]::hygR* |
| JY9955 | *h^+^ lys1Δ::P_adh21_-6×HA-wis1-DD[S469D, T473D]::hygR ura4-D18 leu1-32* |
| JY9957 | *h^+^ lys1Δ::P_adh11_-6×HA-wis1-DD[S469D, T473D]::hygR ura4-D18 leu1-32* |
| JY10080 | *h^?^ nda3-KM311 apc15Δ::kanR cdc13-117::cdc13-GFP::LEU2*  *lys1Δ:: P_adh11_-6×HA-pek1(DD)[S234D,T238D]::hygR* |
| JY10094 | *h^?^ nda3-KM311 apc15Δ::kanR cdc13-117::cdc13-GFP::LEU2* |
| JY10095 | *h^?^ nda3-KM311 apc15Δ::kanR cdc13-117::cdc13-GFP::LEU2*  *lys1Δ::P_adh11_-6×HA-wis1(DD)[S469D,T473D]::hygR* |
| JY10104 | *h^?^ nda3-KM311 slp1Δ::ura4^+^ lys1Δ::P_slp1(1504)_-slp1::hygR*  *cdc13-117::cdc13-GFP::LEU2 Z::P_adh11_-6×HA-pek1(DD)[S234D,T238D]::hygR* |
| JY10106 | *h^+^* *nda3-KM311 mad2Δ::ura4^+^ cdc13-117::cdc13-GFP::LEU2 ura4^-^* |
| JY10112 | *h^?^ nda3-KM311 cdc13-117::cdc13-GFP:LEU2 ura4::P_adh11_-6xHA-pek1(DD)::kanR* |
| JY10121 | *h^-^ mad2∆::ura4 his3-D1 ura4-D18 leu1-32* |
| JY10123 | *h^+^ nda3-KM311 lid1-TAP-kanR mad2Δ::ura4^+^ mad3-GFP-his3^+^* |
| JY10133 | *h^-^ nda3-KM311 mad2Δ::ura4^+^ cdc13-117::cdc13-GFP::LEU2 lys1Δ::P_adh11_-6×HA-wis1(DD)[S469D,T473D]::hygR ura4^-^* |
| JY10135 | *h^?^ nda3-KM311 slp1Δ::ura4^+^ lys1Δ::P_slp1(1504)_-slp1::hygR cdc13-117::cdc13-GFP::LEU2 ura4:: P_adh11_-6×HA-wis1(DD)[S469D,T473D]::hygR* |
| JY10175 | *h^-^ nda3-KM311 mad2Δ::ura4^+^ cdc13-117::cdc13-GFP::LEU2*  *lys1Δ::P_adh11_-6×HA-wis1(DD) [S469D,T473D]::hygR Z::P_mad2_-mad2::kanR ura4^-^* |
| JY10181 | *h^+^* *nda3-KM311 mad2Δ::ura4^+^ cdc13-117::cdc13-GFP::LEU2 Z::P_mad2_-mad2::kanR ura4^-^* |
| JY10259 | *h^+^ nda3-KM311 mad2Δ::ura4^+^ cdc13-117::cdc13-GFP::LEU2 ade6^+^::P_adh21_-mad2::natR* |
| JY10366 | *h^?^ nda3-KM311 cdc13-117::cdc13-GFP::LEU2 slp1Δ::ura4^+^*  *lys1Δ::P_slp1(1504)_-slp1(S28A,T31A)-T_adh1_::hygR* |
| JY10368 | *h^+^ nda3-KM311 apc15-13myc:: kanR* |
| JY10452 | *h^-^ nda3-KM311 cdc13-117::cdc13-GFP::LEU2 slp1Δ::ura4^+^*  *lys1Δ::P_slp1(1504)_-slp1(K19E,K20E,R21E)-hygR ura4^-^* |
| JY10454 | *h^+^ nda3-KM311 cdc13-117::cdc13-GFP::LEU2 slp1Δ::ura4^+^ lys1Δ::P_slp1(1504)_-slp1(K47E,R48E)-hygR ura4^-^* |
| JY10456 | *h^-^ nda3-KM311 cdc13-117::cdc13-GFP::LEU2 slp1Δ::ura4^+^ lys1Δ::P_slp1(1504)_-slp1(K19E,K20E,R21E,K47E,R48E)-hygR ura4^-^* |
| JY10978 | *h^+^ nda3-KM311 cdc13-117:cdc13-GFP:LEU2 lys1Δ::P_slp1(1504)_-slp1(R21E,R48E)::hygR* |
| JY11000 | *h^+^ nda3-KM311 cdc13-117:cdc13-GFP:LEU2 slp1Δ::ura4^+^*  *lys1Δ::P_slp1(1504)_-slp1(T480A)::hygR* |
| JY11001 | *h^+^ nda3-KM311 cdc13-117::cdc13-GFP:LEU2 slp1Δ::ura4^+^ lys1Δ::P_slp1(1504)_-slp1(S76A ,T480A)::hygR* |
| JY11002 | *h^+^ nda3-KM311 cdc13-117::cdc13-GFP::LEU2 slp1Δ::ura4^+^*  *lys1Δ::P_slp1(1504)_-slp1(K479R)-T_adh1_::hygR* |
| JY11007 | *h+ nda3-KM311 cdc13-117:cdc13-GFP:LEU2 slp1Δ::ura4^+^*  *lys1Δ::P_slp1(1504)_-slp1(R21E,R48E)::hygR ura4^-^* |
| JY11039 | *h^+^ nda3-KM311 apc15-13myc:: kanR cdc2-asM17-bsd leu1-32 ura4-D18 ade6-M216* |
| JY11057 | *h^+^ nda3-KM311 cdc13-117::cdc13-GFP::LEU2 slp1Δ::natR*  *lys1Δ::P_slp1(1504)_-slp1(K472R,K479R)-T_adh1_::hygR* |
| JY11060 | *h^+^ nda3-KM311 cdc13-117::cdc13-GFP::LEU2 slp1Δ::ura4^+^*  *lys1Δ::P_slp1(1504)_-slp1(K472R,K479R)-T_adh1_::hygR* |
| JY11073 | *h^-^ nda3-KM311 cdc13-117:cdc13-GFP:LEU2 slp1Δ::ura4+*  *lys1::P_slp1(1504)_-slp1::natR ura4^-^* |
| JY11133 | *h^?^ nda3-KM311 apc15-13myc:: kanR slp1Δ::ura4^+^*  *lys1Δ::P_slp1(1504_)-slp1(S28A,T31A)::hygR ura4^-^* |
| JY11154 | *h^+^* *nda3-KM311 cdc13-117:cdc13-GFP:LEU2*  *lys1**Δ::P_slp1(1504)_-slp1(K479R)-T_slp1_::hygR* |
| JY11157 | *h^-^ nda3-KM311 slp1Δ::ura4+ pmk1Δ::ura4+ lys1::P_slp1(1504)_-slp1::natR*  *cdc13-117:cdc13-GFP:LEU2 ura4-D18* |
| JY11158 | *h^-^ nda3-KM311 cdc13-117::cdc13-GFP::LEU2 slp1Δ::ura4^+^*  *lys1::P_slp1(1504)_-slp1-T_adh1_::natR Z<<P_adh11_-6×HA-pek1(DD)[S234D,T238D]::kanR* |
| JY11177 | *h^+^ nda3-KM311 cdc13-117::cdc13-GFP::LEU2 slp1Δ::ura4^+^*  *lys1Δ::P_slp1(1504)_-slp1(K479R)-T_adh1_::hygR*  *Z<<P_adh11_-6×HA-pek1(DD)[S234D,T238D]::kanR* |
| JY11181 | *h^?^ nda3-KM311 cdc13-117:cdc13-GFP:LEU2 slp1Δ::ura4^+^*  *lys1Δ::P_slp1(1504)_-slp1-T_slp1_::hygR ura4^-^* |
| JY11182 | *h^?^ nda3-KM311 cdc13-117:cdc13-GFP:LEU2 slp1Δ::ura4^+^*  *lys1Δ::P_slp1(1504)_-slp1(K479R)-T_slp1_::hygR ura4^-^* |
| JY11183 | *h^+^ nda3-KM311 cdc13-117:cdc13-GFP:LEU2 slp1Δ::ura4^+^*  *lys1Δ::P_slp1(1504)_-slp1(T480A)-T_slp1_::hygR ura4^-^* |
| JY11185 | *h^-^ nda3-KM311 slp1Δ::ura4^+^cdc13-117::cdc13-GFP:LEU2*  *lys1Δ::P_slp1(1504)_-slp1(S28A,T31A,S59A)-T_adh1_::hygR* |
| JY11189 | *h^?^ nda3-KM311 cdc13-117::cdc13-GFP::LEU2 slp1Δ::ura4^+^*  *lys1Δ::P_slp1(1504)_-slp1(S28E,T31E,S59E)-T_adh1_::hygR* |
| JY11190 | *h^?^ nda3-KM311 cdc13-117::cdc13-GFP::LEU2 slp1Δ::ura4^+^ pmk1Δ::ura4^+^*  *lys1Δ::P_slp1(1504)_-slp1(S28E,T31E,S59E)-T_adh1_::hygR* |
| JY11191 | *h^?^ nda3-KM311 cdc13-117::cdc13-GFP::LEU2 slp1Δ::ura4^+^*  *lys1Δ::P_slp1(1504)_-slp1(S76E,T480E)-T_adh1_::hygR* |
| JY11192 | *h^?^ nda3-KM311 cdc13-117::cdc13-GFP::LEU2 slp1Δ::ura4^+^ pmk1Δ::ura4^+^*  *lys1Δ::P_slp1(1504)_-slp1(S76E,T480E)-T_adh1_::hygR* |
| JY11193 | *h^+^ nda3-KM311 cdc13-117:cdc13-GFP:LEU2*  *lys1Δ::P_slp1(1504)_-slp1(K472R)-T_adh1_::hygR* |
| JY11194 | *h^?^ nda3-KM311 cdc13-117:cdc13-GFP:LEU2 slp1Δ::ura4^+^*  *lys1Δ::P_slp1(1504)_-slp1(PDSP-K19A,K20A,R21A,K47A,R48A)-T_adh1_::hygR* |
| JY11195 | *h^?^ nda3-KM311 cdc13-117::cdc13-GFP:LEU2 slp1Δ::ura4^+^ pmk1Δ::ura4^+^*  *lys1Δ::P_slp1(1504)_-slp1(PDSP-K19A,K20A,R21A,K47A,R48A)**-T_adh1_::hygR* |
| JY11196 | *h^?^ nda3-KM311 cdc13-117::cdc13-GFP::LEU2 slp1Δ::ura4^+^*  *lys1Δ::P_slp1(1504)_-slp1(K19A,K20A,R21A,K47A,R48A)-T_adh1_::hygR* |
| JY11197 | *h^?^ nda3-KM311 cdc13-117::cdc13-GFP::LEU2 slp1Δ::ura4^+^*  *lys1Δ::P_slp1(1504)_-slp1(K19A,K20A,R21A,K47A,R48A)-T_adh1_::hygR*  *Z<<P_adh11_-6×HA-pek1(DD)[S234D,T238D]::kanR* |
| JY11198 | *h^?^ nda3-KM311 cdc13-117::cdc13-GFP::LEU2* *lys1Δ::P_slp1(1504)_-slp1(K19A,K20A,R21A,K47A,R48A)-T_adh1_::hygR Z<<P_adh11_-6xHA-pek1(DD)[(S234D, T238D)] ::kanR* |
| JY11199 | *h^?^ nda3-KM311 cdc13-117::cdc13-GFP:LEU slp1Δ::ura4^+^ pmk1Δ::ura4^+^ lys1Δ::P_slp1(1504)_-slp1::natR Z<<P_adh11_-6xHA-pek1(DD)[(S234D, T238D)]::kanR* |
| JY11200 | *h^?^ nda3-KM311 cdc13-117::cdc13-GFP:LEU2 slp1Δ::ura4^+^*  *lys1Δ::P_slp1(1504)_-slp1(S28A,T31A, S59A)-T_adh1_::hygR Z<<P_adh11_-6xHA-pek1(DD)[(S234D, T238D)]::kanR* |
| JY11201 | *h^?^ nda3-KM311 cdc13-117::cdc13-GFP:LEU2 slp1Δ::ura4^+^ lys1Δ::P_slp1(1504)_-slp1(S28A,T31A,S59A)-T_adh1_::hygR* |
| JY11202 | *h^?^ nda3-KM311 cdc13-117::cdc13-GFP:LEU2 slp1Δ::ura4^+^ lys1Δ::P_slp1(1504)_-slp1(S28A,T31A,S59A)-T_adh1_::hygR Z<<P_adh11_-6xHA-pek1(DD)[(S234D, T238D)]::kanR* |
| JY11203 | *h^?^ nda3-KM311 cdc13-117::cdc13-GFP:LEU2 slp1Δ::ura4^+^ lys1Δ::P_slp1(1504)_-slp1(S28A,T31A,S59A,S76A)-T_adh1_::hygR* |
| JY11204 | *h^?^ nda3-KM311 cdc13-117::cdc13-GFP:LEU2 slp1Δ::ura4^+^ lys1Δ::P_slp1(1504)_-slp1(S28A,T31A,S59A,S76A)-T_adh1_::hygR Z<<P_adh11_-6xHA-pek1(DD)[(S234D, T238D)]::kanR* |
| JY11205 | *h^?^ nda3-KM311 cdc13-117::cdc13-GFP:LEU2 slp1Δ::ura4^+^ lys1Δ::P_slp1(1504)_-slp1(S76A,T480A)-T_adh1_::hygR Z<<P_adh11_-6xHA-pek1(DD)[(S234D, T238D)]::kanR* |
| JY11206 | *h^?^ nda3-KM311 cdc13-117::cdc13-GFP:LEU2 slp1Δ::ura4^+^ lys1Δ::P_slp1(1504)_-slp1(S28A,T31A,S59A,S76A,T480A)-T_adh1_::hygR* |
| JY11207 | *h^?^ nda3-KM311 cdc13-117::cdc13-GFP:LEU2 slp1Δ::ura4^+^ lys1Δ::P_slp1(1504)_-slp1(S28A,T31A,S59A,S76A,T480A)-T_adh1_::hygR Z<<P_adh11_-6xHA-pek1(DD)[(S234D, T238D)]::kanR* |
| JY11208 | *h^?^ nda3-KM311 lys1Δ::P_adh11_-6xHA-pek1(DD)[(S234D, T238D)]::hygR*  *atf1Δ::natR cdc13-117:cdc13-GFP::LEU2* |
| JY11209 | *h^?^ nda3-KM311 lys1Δ::P_adh11_-6xHA-pek1(DD)[(S234D, T238D)]::hygR pmk1Δ::ura4^+^ cdc13-117::cdc13-GFP::LEU2* |
| JY11210 | *h^?^ nda3-KM311 apc15-13myc:: kanR slp1Δ::ura4^+^*  *lys1Δ::P_slp1(1504_)-slp1::hygR ura4^-^* |
| JY11222 | *h^?^ nda3-KM311 cdc13-117::cdc13-GFP::LEU2 slp1Δ::ura4^+^*  *lys1Δ::P_slp1(1504)_-slp1(T480S)-T_adh1_::hygR* |
| JY11227 | *h^?^ nda3-KM311 cdc13-117::cdc13-GFP::LEU2 slp1Δ::ura4^+^*  *lys1Δ::P_slp1(1504)_-slp1(K472R,K479R)-T_adh1_::hygR*  *Z<<P_adh11_-6×HA-pek1(DD)[S234D,T238D]::kanR* |
| JY11229 | *h^?^ nda3-KM311 cdc13-117::cdc13-GFP::LEU2 slp1Δ::ura4^+^*  *lys1Δ::P_slp1(1504)_-slp1(K472R)-T_adh1_::hygR* |
| JY11230 | *h^?^ nda3-KM311 cdc13-117::cdc13-GFP::LEU2 slp1Δ::ura4^+^*  *lys1Δ::P_slp1(1504)_-slp1(K472R)-T_adh1_::hygR*  *Z<<P_adh11_-6×HA-pek1(DD)[S234D,T238D]::kanR* |
| JY11263 | *h^?^ nda3-KM311 slp1Δ::ura4^+^ cdc13-117::cdc13-GFP:LEU2 lys1Δ::P_slp1(1504)_-slp1(T480A)::hygR Z<<P_adh11_-6xHA-pek1(DD)[(S234D, T238D)] ::kanR* |
| JY11289 | *h^?^ nda3-KM311 slp1Δ::ura4^+^ atf1Δ::natR lys1Δ::P_slp1(1504)_-slp1::natR cdc13-117::cdc13-GFP:LEU2 Z<<P_adh11_-6xHA-pek1(DD)[(S234D, T238D)]::kanR* |
| JY11295 | *h^+^ mts3-1 ura4^-^ leu1^-^ ade6^-^* |
| JY11296 | *h^?^ nda3-KM311 cdc13-117:cdc13-GFP:LEU2 slp1Δ::ura4^+^*  *lys1Δ::P_slp1(1504)_-slp1(S28A;T31A)-T_slp1_::hygR ura4^-^* |
| JY11297 | *h^?^ nda3-KM311 cdc13-117:cdc13-GFP:LEU2 slp1Δ::ura4^+^*  *lys1Δ::P_slp1(1504)_-slp1(S28A;T31A;S59A)-T_slp1_::hygR ura4^-^* |
| JY11307 | *h^?^ nda3-KM311 cdc13-117::cdc13-GFP::LEU2 slp1Δ::ura4^+^*  *lys1Δ::P_slp1(1504)_-slp1(S28AE,T31A)-T_adh1_::hygR*  *Z<<P_adh11_-6×HA-pek1(DD)[S234D,T238D]::kanR* |
| JY11321 | *h^?^ mts3-1 lys1Δ::P _slp1(1504_)-sfGFP-slp1(K472R)::hygR ura4^-^ leu1^-^ ade6^-^* |
| JY11322 | *h^+^ mts3-1 lys1Δ::P_slp1(1504)_-sfGFP-slp1(K472R;K479R)::hygR*  *ura4^-^ leu1^-^ ade6^-^* |
| JY11323 | *h^?^ nda3-KM311 atf1Δ::natMX6 cdc13-117::cdc13-GFP:LEU2 ura4::P_adh11_-6xHA-pek1(DD)::kanR* |
| JY11333 | *h^+^* *mts3-1 lys1Δ::P_slp1(1504)_-sfGFP-slp1(K19A,K20A,R21A,K47A,R48A)-T_adh1_::hygR* |
| JY11334 | *h^+^* *mts3-1 lys1Δ::P_slp1(1504)_-sfGFP-slp1(PDSP-K19A,K20A,R21A,K47A,R48A)-T_adh1_::hygR* |
| JY11335 | *h^+^* *mts3-1 lys1Δ::P_slp1(1504)_-sfGFP-slp1(T480A)-T_adh1_::hygR::lys1* |
| JY11336 | *h^+^* *mts3-1 lys1Δ::P_slp1(1504)_-sfGFP-slp1(S76A,T480A)-T_adh1_::hygR::lys1* |
| JY11337 | *h^+^* *mts3-1 lys1Δ::P_slp1(1504)_-* *sfGFP-slp1(S28A,T31A,S59A)-T_adh1_::hygR::lys1* |
| JY11338 | *h^+^* *mts3-1Δ:ura4^+^*  *lys1Δ::P_slp1(1504)_-* *sfGFP-slp1(S28A,T31A,S59A,S76A)-T_adh1_::hygR::lys1* |
| JY11339 | *h^+^* *mts3-1 lys1Δ::P_slp1(1504)_-sfGFP-slp1(S28A,T31A,S59A,S76A,T480A)-T_adh1_::hygR::lys1* |
| JY11340 | *h^?^ mts3-1 lys1Δ::P_slp1(1504)_-sfGFP-slp1(K479R)-T_adh1_::hygR::lys1* |
| JY11355 | *h^+^ mts3-1 slp1::P_slp1_-sfGFP-slp1 ade6::P_adh1_-GST-slp1(456-488aa)-natR* |
| JY11356 | *h^+^ mts3-1 slp1::P_slp1_-sfGFP-slp1 pmk1Δ::ura4^+^ ade6::P_adh1_-GST-slp1(456-488aa)::natR* |
| JY11357 | *h^-^ mts3-1 slp1::P_slp1_-sfGFP-slp1 ade6::P_adh1_-GST-slp1(456-488aa)-natR lys1Δ::P_adh11_-6xHA-pek1(DD)::hygR* |
| JY11368 | *h^?^mts3-1 his5∆::P_adh11_-6xHA-pek1(DD)::natR lys1∆::P_slp1(1504)_-sfGFP-slp1(K19E,K20E,R21E,K47E,R48E)-T_adh1_::hygR ura4- leu1- ade6-* |
| JY11369 | *h^?^ mts3-1 his5∆::Padh11-6xHA-pek1(DD)::natR lys1∆::P_slp1(1504)_-sfGFP-slp1(PDSP-5E)-T_adh1_::hygR ura4- leu1- ade6-* |
| JY11370 | *h^?^ mts3-1 his5∆::Padh11-6xHA-pek1(DD)::natR lys1∆::P_slp1(1504)_-sfGFP-slp1(K479R)-T_adh1_::hygR ura4- leu1- ade6-* |
| JY11371 | *h^?^ mts3-1 his5∆::Padh11-6xHA-pek1(DD)::natR lys1∆::P_slp1(1504)_-sfGFP-slp1(K472R)-T_adh1_::hygR ura4- leu1- ade6-* |
| JY11372 | *h^?^ mts3-1 his5∆::Padh11-6xHA-pek1(DD)::natR lys1∆::pUC119-P_slp1(1504)_-sfGFP-slp1(K472R,K479R)-T_adh1_::hygR ura4- leu1- ade6-* |
| JY11373 | *h^?^ mts3-1 his5∆::Padh11-6xHA-pek1(DD)::natR lys1Δ::P_slp1(1504)_-sfGFP-slp1(T480A)-T_adh1_::hygR ura4- leu1- ade6-* |
| JY11374 | *h^?^ mts3-1 his5∆::Padh11-6xHA-pek1(DD)::natR lys1Δ::P_slp1(1504)_-sfGFP-slp1(S76A,T480A)-T_adh1_::hygR ura4- leu1- ade6-* |
| JY11375 | *h^?^ mts3-1 his5∆::Padh11-6xHA-pek1(DD)::natR lys1Δ::P_slp1(1504)_-sfGFP-slp1(S28A,T31A,S59A)-T_adh1_::hygR ura4- leu1- ade6-* |
| JY11376 | *h^?^ mts3-1 his5∆::Padh11-6xHA-pek1(DD)::natR lys1Δ::P_slp1(1504)_-sfGFP-slp1(S28A,T31A,S59A,S50A,S76A)-T_adh1_::hygR ura4- leu1- ade6-* |
| JY11377 | *h^?^ mts3-1 his5∆::Padh11-6xHA-pek1(DD)::natR lys1Δ::P_slp1(1504)_-sfGFP-slp1(S28A,T31A,S59A,S50A,S76A,T480A)-T_adh1_::hygR ura4- leu1- ade6-* |
| JY11378 | *h^?^ mts3-1 his5Δ::P_adh11_-6xHA-pek1(DD)::natR*  *lys1Δ::P _slp1(1504)_-sfGFP-slp1::hygR ura4^-^ leu1^-^ ade6^-^* |
| JY11379 | *h^?^ mts3-1 lys1Δ::P_slp1(1504)_-sfGFP-slp1(WT)-T_adh1_::hygR::lys1* |
| JY11399 | *h^?^ mts3-1 pmk1Δ::ura4^+^ lys1Δ::P _slp1(1504)_-sfGFP-slp1::hygR* |
| JY11400 | *h^?^ mts3-1 his5Δ::P_adh11_-6xHA-pek1(DD)::natR lys1Δ::P_slp1(1504)_-sfGFP-slp1(wt)-T_adh1_::hygR pmk1Δ::kanR ura4- leu1- ade6-* |
| JY11421 | *h^?^ mts3-1 ade6::P_adh1_-slp1(1-60aa)-mEGFP-2xNLS-GST-slp1(456-488aa)::natR* |
| JY11422 | *h^?^ mts3-1 lys1∆::P_adh11_-pmk1-GBP-mCherry::hygR ade6::P_adh1_-slp1(1-60aa)-mEGFP-2xNLS-GST-slp1(456-488aa)::natR* |
| JY11423 | *h^?^ mts3-1 pmk1∆::ura4 ade6::P_adh1_-slp1(1-60aa)-mEGFP-2xNLS-GST-slp1(456-488aa)::natR* |
| JY11424 | *h^?^ mts3-1 ura4::P_adh11_-6xHA-pek1(DD)::kanR ade6::P_adh1_-slp1(1-60aa)-mEGFP-2xNLS-GST-slp1(456-488aa)::natR* |
| JY11425 | *h^?^ mts3-1 ura4::P_adh11_-6xHA-pek1(DD)::kanR lys1∆::P_adh11_-pmk1-GBP-mCherry::hygR ade6::P_adh1_-slp1(1-60aa)-mEGFP-2xNLS-GST-slp1(456-488aa)::natR* |
| JY11494 | *h^+^ ura4-D18 leu1-32 ade6-M210* |
| JY11548 | *h^+^ cdc25-22 pmk1Δ::ura4^+^* |
| JY11552 | *h^?^ P_nmt1_-mad2::leu1 GFP-atb2^+^::kanR* |
| JY11555 | *h^-^ P_nmt1_-mad2::leu1 GFP-atb2^+^::kanR spk1∆::kanR* |
| JY11556 | *h^+^ lys1Δ::P_adh21_-6xHA-byr1(DD)::hygR ura4-D18 leu1-32 ade6-M210* |
| JY11557 | *h^+^ lys1Δ::P_adh11_-6xHA-byr1(DD)::hygR ura4-D18 leu1-32 ade6-M210* |
| JY11558 | *h^?^ nda3-KM311 cdc13-117:cdc13-GFP:LEU2*  *lys1Δ::P_adh21_-6xHA-byr1(DD)::hygR ura4^-^* |
| JY11559 | *h^?^ nda3-KM311 cdc13-117:cdc13-GFP:LEU2*  *lys1Δ::P_adh11_-6xHA-byr1(DD)::hygR ura4^-^* |
| JY11562 | *h^?^ cdc25-22 lys1Δ::P_adh11_-6xHA-pek1(DD)[(S234D, T238D)]::hygR* |
| JY11563 | *h^?^ cdc25-22 slp1Δ::ura4 lys1∆::P_slp1(1504)_-slp1-T_slp1_::hygR ura4^-^* |
| JY11564 | *h^?^ cdc25-22 slp1Δ::ura4 lys1∆::P_slp1(1504)_-slp1(T480A)-T_slp1_::hygR ura4^-^* |
| JY11571 | *h^?^ cdc25-22 GFP-atb2^+^::kanR* |
| JY11572 | *h^?^ cdc25-22 GFP-atb2^+^::kanR pmk1∆::ura4^+^* |
| JY11591 | *h^?^ P_nmt1_-mad2::leu1^+^ GFP-atb2^+^::kanR pmk1Δ::ura4^+^* |
| JY11592 | *h^?^ P_nmt1_-mad2::leu1^+^ GFP-atb2^+^::kanR sty1-T97A::natR* |
